# The Avocado Genome Informs Deep Angiosperm Phylogeny, Highlights Introgressive Hybridization, and Reveals Pathogen-Influenced Gene Space Adaptation

**DOI:** 10.1101/654285

**Authors:** Martha Rendón-Anaya, Enrique Ibarra-Laclette, Alfonso Méndez Bravo, Tianying Lan, Chunfang Zheng, Lorenzo Carretero-Paulet, Claudia Anahí Perez-Torres, Alejandra Chacón-López, Gustavo Hernandez-Guzmán, Tien-Hao Chang, Kimberly M. Farr, W. Brad Barbazuk, Srikar Chamala, Marek Mutwil, Devendra Shivhare, David Alvarez-Ponce, Neena Mitter, Alice Hayward, Stephen Fletcher, Julio Rozas, Alejandro Sánchez Gracia, David Kuhn, Alejandro F. Barrientos-Priego, Jarkko Salojärvi, Pablo Librado, David Sankoff, Alfredo Herrera-Estrella, Victor A. Albert, Luis Herrera-Estrella

## Abstract

The avocado, *Persea americana*, is a fruit crop of immense importance to Mexican agriculture with an increasing demand worldwide. Avocado lies in the anciently-diverged magnoliid clade of angiosperms, which has a controversial phylogenetic position relative to eudicots and monocots. We sequenced the nuclear genomes of the Mexican avocado race, *P. americana* var. *drymifolia*, and the most commercially popular hybrid cultivar, Hass, and anchored the latter to chromosomes using a genetic map. Resequencing of Guatemalan and West Indian varieties revealed that ∼39% of the Hass genome represents Guatemalan source regions introgressed into a Mexican race background. Some introgressed blocks are extremely large, consistent with the recent origin of the cultivar. The avocado lineage experienced two lineage-specific polyploidy events during its evolutionary history. Although gene-tree/species-tree phylogenomic results are inconclusive, syntenic ortholog distances to other species place avocado as sister to the enormous monocot and eudicot lineages combined. Duplicate genes descending from polyploidy augmented the transcription factor diversity of avocado, while tandem duplicates enhanced the secondary metabolism of the species. Phenylpropanoid biosynthesis, known to be elicited by *Colletotrichum* (anthracnose) pathogen infection in avocado, is one enriched function among tandems. Furthermore, transcriptome data show that tandem duplicates are significantly up- and down-regulated in response to anthracnose infection, whereas polyploid duplicates are not, supporting the general view that collections of tandem duplicates contribute evolutionarily recent “tuning knobs” in the genome adaptive landscapes of given species.

**SIGNIFICANCE STATEMENT:** Avocado is a nutritious, economically important fruit species that occupies an unresolved position near the earliest evolutionary branchings of flowering plants. Our nuclear genome sequences of Mexican and Hass variety avocados inform ancient evolutionary relationships and genome doublings, the admixed nature of Hass, and provide a look at how pathogen interactions have shaped avocado’s more recent genomic evolutionary history.

## INTRODUCTION

The avocado, *Persea americana*, is a commercially important tree fruit species in the Lauraceae family, otherwise known for the spices cinnamon, bay leaves, and sassafras (gumbo filé) (1). Lauraceae is contained within the early diverging magnoliid lineage of angiosperms, which at about 11,000 total species is minuscule in comparison to the dominant eudicot and monocot flowering plant lineages, comprising about 285,000 species combined (2). Avocados are a vital crop for Mexico, from which almost 50% of all avocado exports originate, valued at about $2.5 billion U.S. dollars *. Although the avocado has an ancient cultivation history in Mexico and Central to South America (3), its extreme worldwide popularity as an oily, nutty-flavored fruit with highly beneficial nutritional properties dates mainly from the early 20^th^ century (4). Cultivated avocados occur in three landraces with possibly independent cultivation origins that reflect their current distribution: the Mexican, Guatemalan, and West Indian varieties, respectively (4). The principal industrial avocado cultivar is known as Hass, after the grower who first patented it in 1935. Hass represents a hybrid between the Guatemalan and Mexican races, but its precise breeding history is unknown (4, 5). Here, we generate and analyze the complete genome sequences of a Hass individual and a representative of the highland Mexican landrace, *Persea americana* var. *drymifolia*. We also study genome resequencing data for other Mexican individuals, as well as Guatemalan and West Indian accessions. We use these data to study the admixed origin of the Hass cultivar, and demonstrate its race-wise parentage more precisely. We evaluate the phylogenetic origin of avocado among angiosperms and provide information on avocado’s unique polyploid ancestry. The adaptive landscape of the avocado genome in terms of its duplicate gene functional diversity was also explored. We further evaluate gene expression patterns during the defense response of Hass avocado to anthracnose disease, and how this is partitioned by gene duplication mechanisms.

## RESULTS AND DISCUSSION

### Plant Material, Genome Assembly, and Annotation

Due to growing market demand, 90% of cultivated avocado corresponds to the cultivar Hass, which in Mexico is commonly grafted on Mexican race (*P. americana* var. *drymifolia*) rootstock (4). This practice makes it possible to maintain high productivity as the indigenous race is well adapted to Mexican highland soils. The Hass cultivar and Mexican race were chosen to generate reference genomes (Table S1; Supplementary information section 1). Additionally, to explore the genetic diversity available in avocado, we resequenced representative individuals from the three avocado botanical varieties (vars. *drymifolia, guatemalensis* and *americana*), including the disease-resistant rootstock Velvick (6), an additional Hass specimen and the early flowering/fruiting Hass somatic mutant Carmen Hass cultivar (otherwise known as Mendez No. 1 (4, 7)), as well as wild avocados of the West Indian variety (*P. americana* var. *costaricensis*). *Persea shiedeana* (the edible coyo), a species relatively closely related to *P. americana* (8), was also included (Tables S1,S2).

*De novo* and evidence-directed annotation revealed a similar number of protein coding genes in each genome: 22,441 from the Mexican race and 24,616 from Hass (Table 1, Tables S8, S9; Supplementary information section 2). We next used the BUSCO software to estimate the presence of 1,440 conserved embryophyte single copy genes (9) in the annotations, leading to estimated completeness percentages of 85% and 86.3% for Hass and Mexican avocado, respectively (Table 1). The Mexican race was sequenced using the short read, high coverage, Illumina sequencing platform, while the Hass genome was sequenced using the long-read Pacific Biosciences sequencing technology. Given the similar BUSCO scores, we used the larger Hass genome assembly for downstream SNP calling, as PacBio technology lowers the probability of contig misassembly and permits incorporation of substantially more repetitive DNA sequence and genes lying within it into the assembled genome, which might have otherwise been missed.

**Table 1.** General statistics of the avocado assemblies and their annotations (Tables S8-S15).

|  | *Persea americana* assembly |  |
| --- | --- | --- |
|  | var. *drymifolia* | 'Hass' cultivar |
| Number of contigs | 99,957 | 8,135 |
| Total length of contigs (bp) | 668,137,248 | **912,697,600** |
| Number of scaffolds | 42,722 | -- |
| Total length of scaffolds (bp) | **823,419,498** | -- |
| Longest contig / scaffold (bp) | 254,240 / 4,610,966 | 2,811,280 / -- |
| Mean contig / scaffold length (bp) | 6,684 / 19,274 | 112,194 / -- |
| N50 contig / scaffold length (bp) | 11,724 / 323,854 | 296,371 / -- |
| Assembly in scaffolded contigs | 87.6% | 0% |
| Assembly in unscaffolded contigs | 12.4% | 100% |
| Protein coding genes (% BUSCO completeness) | 22,441 (86.3%) | 24,616 (85%) |
| ncRNAs | 4,413 | 8,815 |
| TEs | 11,372 | 20,611 |

We also anchored the Hass genome to an avocado genetic map (10). Two large mapping populations of 1,339 trees were genotyped with 5,050 single nucleotide polymorphism (SNP) markers from transcribed genes, and the resulting map was used to order the Hass scaffolds into twelve linkage groups, matching the avocado haploid chromosome number (see, e.g., chromosome 4, Fig. 2A). The total length of the anchored genome accounts for 46.2% of the Hass assembly, and represents 915 scaffolds, 361 of which could be oriented (Tables S5-S7; Figs. S1-S12).

**Figure 1.**
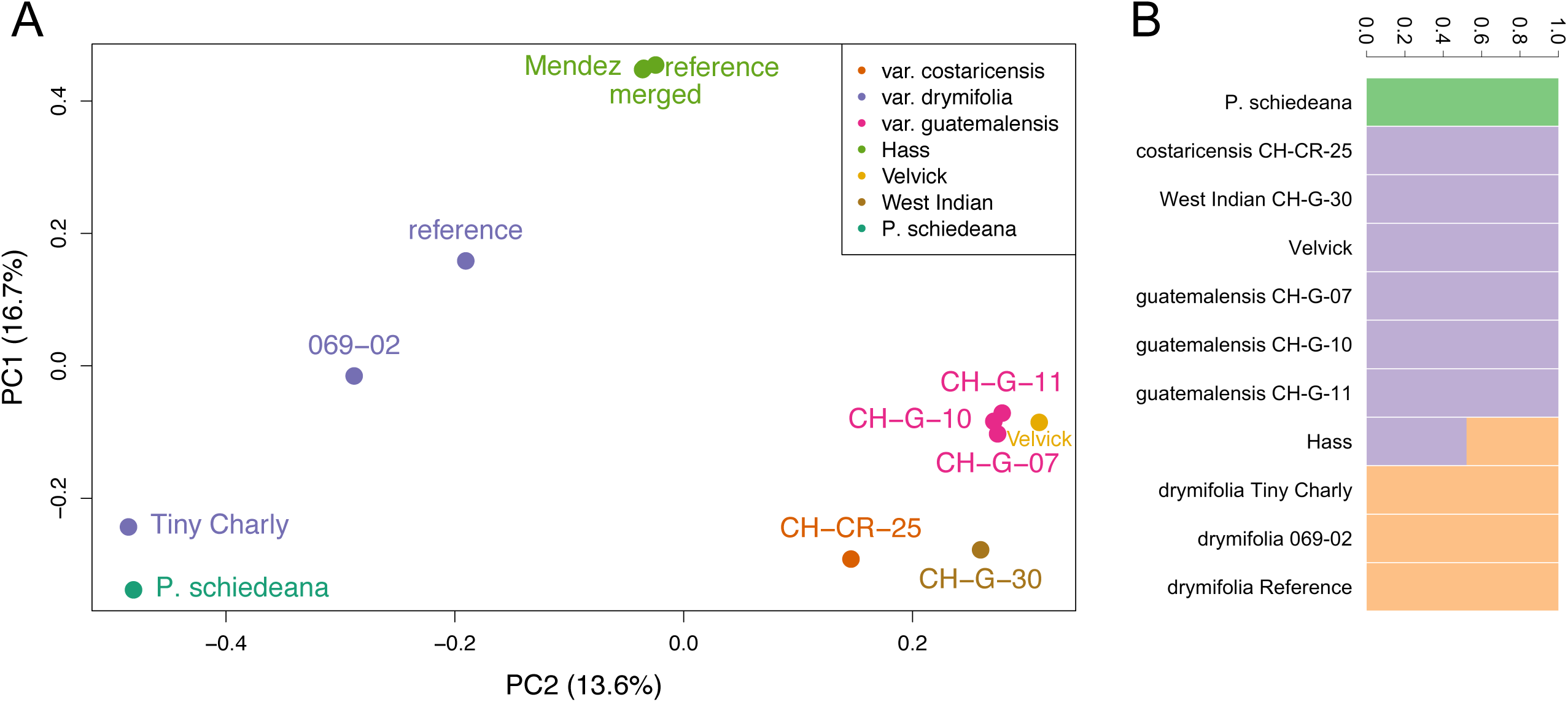
Population genomic structure of avocado. **A**. Principal component analysis (PCA) of genome-wide SNPs reveals population groupings among races and varieties. The Guatemalan and West Indian/Costa-Rican accessions are closely related, while the Mexican (*P. americana* v. *drymifolia*) specimens are more diverse, with the unusual individual Tiny Charly drawn toward the outgroup species *P. schiedeana* by PC2. Hass and its sport Mendez are tightly clustered and intermediate between Mexican and Guatemalan and West Indian/Costa-Rican on PC2. **B**. NGSAdmix analysis reveals similar population structure at K=3. The *Persea schiedeana* outgroup is distinct, and the Hass reference genome is revealed to be admixed between Guatemalan-West Indian and Mexican source populations, the Mexican source clearly contributing greater than 50%.

**Figure 2.**
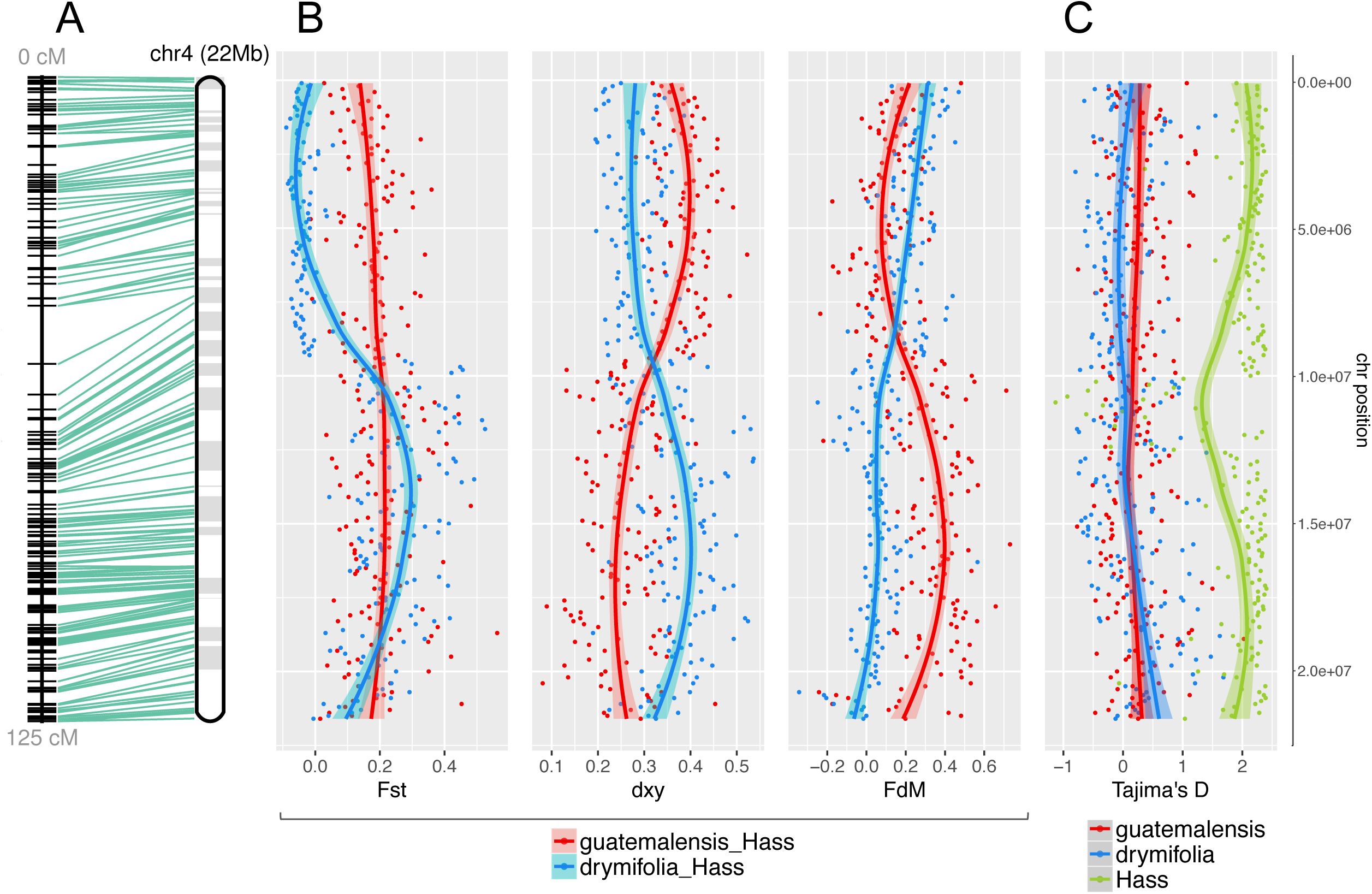
SNP diversity analysis reveals the hybrid genomic background of Hass avocado. **A**. Twenty-two megabases (Mb) of anchored DNA on chromosome 4 exemplify the hybrid nature of Hass, in which genomic introgression from the Guatemalan avocado race (variety *guatemalensis*) occurred into a Mexican (variety *drymifolia*) genetic background. **B**. While in the top chromosome arm the blue trend line shows a low differentiation index (F_ST_) between Hass and the Mexican subpopulation as well as a high introgression signal 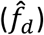 from var. *drymifolia* into Hass, these signals should not be misinterpreted as introgression events, since the absolute genetic divergence (*d*_*XY*_) between both sets of accessions does not vary along the chromosome. The lower arm of the chromosome, however, has inverted trends, where our estimators describe an elevated introgression signal from var. *guatemalensis* into Hass, as confirmed by the decay in *d*_*XY*_ (red trend line), and higher F_ST_ between Hass and Mexican accessions. **C**. No evidence for selective sweeps or domestication signatures were identified; Mexican and Guatemalan subpopulations displayed neutral D values while Hass maintained extreme D values at the theoretical upper limit of the estimator (≈ 2). Such positive values reflect a “bottlenecked” origin with clonal expansion after the very recent foundation of the cultivar only a few decades ago. Each dot in the plots corresponds to statistics for SNP data in non-overlapping 100 Kb windows (confidence interval of 0.90 for graphical smoothed conditional means). Apparent centromeric regions are located at around 10 Mb, where 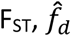 and *d*_*XY*_ intersect and Tajima’s D for Hass decreases.

### Single Nucleotide Polymorphisms, Population Structure, and the Parentage of Hass

To study avocado from a population genomic perspective, we resequenced accessions of different races and cultivars and mapped the reads against the Hass reference genome assembly (Table S2). The estimated depth of coverage ranged from 3.3 - 39X, with breadth of coverage between 70% - 92% (Table S11; Supplementary information section 3.2). Given the uneven sequencing coverage, we used ANGSD to call SNPs across the entire (unanchored) genome assembly, followed by a stringent pruning based on per-site depth, minor allele frequencies and linkage disequilibrium, that resulted in 179,029 high quality SNP variants (Supplementary data 2). Phylogenetic, principal component and identity-by-state (IBS) analyses derived from this data set (Fig. 1A; Figs. S13-S16) cluster the samples belonging to the Hass cultivar and Guatemalan variety into two groups as expected according to their genetic background. Principal component analysis of genome-wide SNPs showed relative uniformity in Costa-Rican/West Indian/Guatemalan group, but strong heterogeneity within the Mexican subpopulation, wherein the unusual accession Tiny Charly is a divergent sample (Fig. S15). SNPhylo (11) results reflected the poor fit of the SNP data to a bifurcating tree by embedding the hybrid Hass within an otherwise Mexican clade, one known parent of this hybrid cultivar (Fig. S13). Furthermore, in that lineage’s sister group, Guatemalan accessions were derived within an otherwise Costa-Rican/West Indian lineage, suggesting an admixed origin involving Guatemalan and other sources. Phylogenetic patterns generated from chromosome-wise SNP subsets (based only on contigs anchored to chromosomes) recapitulated these relationships for 7 of avocado’s 12 chromosomes, whereas one chromosome supported Hass to be sister to the Costa-Rican/West Indian lineage, perhaps reflective of chromosomal differences in the admixture proportions of this hybrid cultivar (Fig. S14; Supplementary data 2). Furthermore, IBS clustering placed Hass intermediate between the Guatemalan and Mexican subpopulations, agreeing with the hybrid nature of this variety (Fig. S16).

To account for further evidence of admixture in the Hass reference genome, we used NGSAdmix (12) modeling different possible numbers of source populations (K = 1-6) (Fig. S17; Supplementary information section 5). The Akaike information criterion (AIC) indicated K = 1 as the preferred model, reflecting poor population structuring within avocado as a whole. However, since we know Hass is admixed *a priori*, we chose the smallest (most parsimonious) K for which Hass admixture appears (K = 3). This criterion predicts the following three populations: (i) *Persea schiedeana*, (ii) the West Indian plus the Guatemalan varieties, and the (iii) Mexican accessions (Fig. 1B). Combining the IBS and NGSAdmix observations, we specifically calculated the contribution of Guatemalan and Mexican backgrounds into the Hass subpopulation. EIGMIX (13) revealed that the greatest admixture proportion, 61%, stemmed from the Mexican race (Fig. S18; Table S13).

Although based on ~46% anchoring of scaffolds to chromosomes, we investigated chromosome-wise signatures of admixture in the Hass genome (Supplementary section 5.2). We calculated the 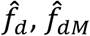 and *d*_*XY*_ estimators of introgression and divergence (Fig. S19; Table S14) according to Martin et al. (14) in non-overlapping 100Kb windows, controlling the directionality of gene-flow from the Guatemalan race into Hass versus the Mexican race into Hass, setting *P. schiedeana* as the outgroup and leaving Tiny Charly out of the Mexican subpopulation to avoid the bias this divergent accession could introduce into calculations (Fig. 2B). Genomic regions that behave as 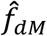 outliers can be distinguished as introgressed from ancestral variation if the absolute genetic distance *d*_*XY*_ is also reduced between donor (P_3_) and receptor (P_2_). In the presence of gene flow, genomic windows coalesce more recently than the lineage split, so the magnitude of reduction in P_2_-P_3_*d*_*XY*_ is greater than in the case where recombination and hybridization are absent. We evaluated several 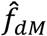 cut-offs (Q50, 75, 90; Fig. S20) and observed a remarkable reduction of genetic divergence in the scenario where gene flow occurs from the Guatemalan race into Hass. Considering those blocks with 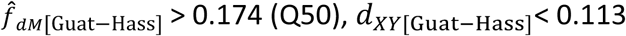 (mean divergence between subpopulations) and 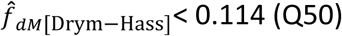, we were able to define 840 high-confidence regions of Guatemalan origin across the 12 chromosomes (Supplementary information section 5.2; Figs. 2 and S21,23-34). Chromosome 4 illustrates these analyses well, demonstrating that a huge Guatemalan block, which could encompass an entire chromosome arm, is present in the Hass genome (Fig. 2A). The length of this Guatemalan-derived block, uninterrupted by recombination, reflects the extremely recent hybrid origin of the cultivar.

We also calculated the level of nucleotide diversity (π (15, 16)) in each population (Mexican, Guatemalan, Hass), the F_ST_ index (17) to determine regions of high differentiation between varieties, and Tajima’s D (18) in order to evaluate any deviations from neutral evolution (Table S14; Supplementary information section 5.3). We observed that Hass has the lowest nucleotide diversity (π = 0.06) and very high Tajima’s D in all chromosomes (genomic average of 1.5), as expected for individuals derived from a recent founder event and clonally propagated; these values contrast with the low, positive Tajima’s D values in the Mexican and Guatemalan populations (genomic averages of 0.19 and 0.11, respectively; Fig. S22). In the case of chromosome 4, the F_ST_ index between the Mexican race and Hass corroborates our previous conclusions on admixture, showing that approximately half of the chromosome corresponds to the Mexican background whereas the other half has its origin in the Guatemalan population (Fig. 2B).

### Whole Genome Duplication History

Next, we investigated *Persea americana* genome structural history and relatedness of the species to other major groups of angiosperms (Supplementary information section 6). We used the CoGe SynMap tool (19) to examine avocado self:self and avocado:*Amborella* synteny. *Amborella*, the single living representative of the sister lineage to all other angiosperms, is known to show a 1:3 syntenic block relationship compared with *Vitis* (grapevine) and its ancient hexaploid structure shared with all core eudicots (20-23). As such, the *Amborella* genome displays no additional whole genome duplications (WGDs) since the angiosperm last common ancestor (LCA). Consequently, paralogous syntenic blocks discovered within self:self and avocado:*Amborella* plots could reflect WGDs unique to the avocado lineage that occurred since last common ancestry with *Amborella*. Using these approaches, we discovered that the avocado genomes show evidence for two ancient polyploidy events (Fig. 3A-C). We investigated the relative timing of these events with respect to the gamma hexaploidy (21, 22) and species splits using K_s_ density plots of orthologous and paralogous gene pairs between avocado, *Amborella*, and *Vitis* (Supplementary information section 6.2). Avocado polyploidy events are lineage-specific as both postdate the divergence of the avocado lineage from common ancestry with either *Amborella* or *Vitis* (Figs. 3C and S36). Considerable fractionation (alternative deletion of duplicated genes between polyploid subgenomes) since these two polyploidy events is observable in blockwise relationships of about 4:1 (Figs. 3B, S35). The blockwise relationship of 4:1 for avocado:*Amborella* suggests although not proves that the two events were WGDs and not triplications. Further quantitative analysis using well-conserved orthologous syntenic “superblocks” (24) between avocado and 15 other angiosperms strongly supported the conclusion that the most recent polyploidy event in avocado was a WGD and not a triplication, as in the ancient gamma hexaploidy event (Supplementary information section 6.3).

**Figure 3.**
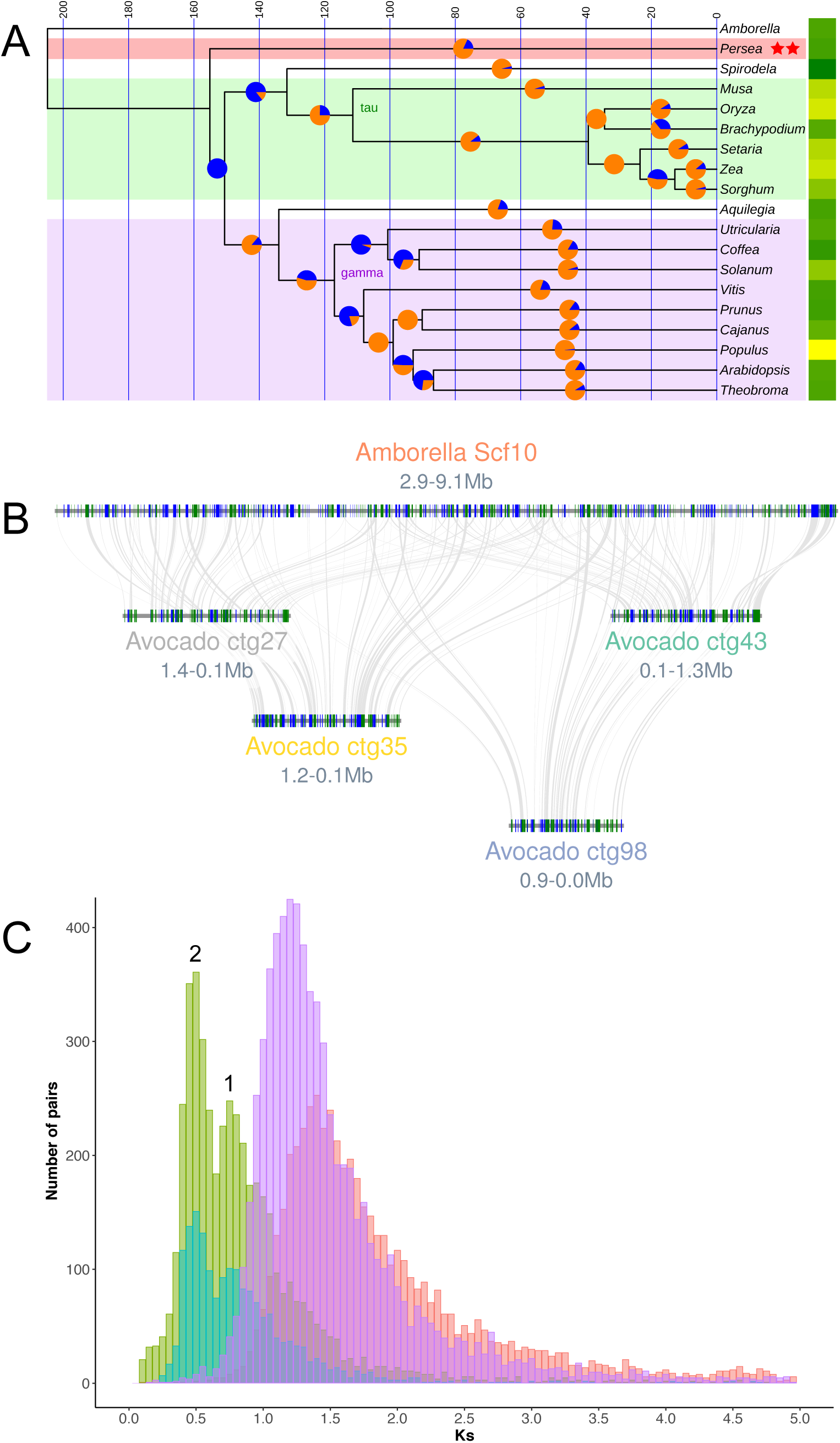
Phylogenomic and whole genome duplication history of avocado. **A**. An ultrametric time-tree based on universally present single-copy protein sequences depicts one of 3 common resolutions of *Persea* (Magnoliidae) relationships to other flowering plants. This topology, showing avocado sister to monocots plus eudicots, mirrors phylogenetic relationships derived from syntenic distances. Here, the split time between the last common ancestor of avocado and the monocot/eudicot crown group is less than 4 million years. Pie charts at 50% positions on branches show proportions of gene gains (orange) versus losses (blue) as determined by BadiRate’s birth-death-innovation model. Yellow-green (greater-lesser) heatmap to the right of the tree depicts relative numbers of genes in the modern genomes. Syntenic analysis revealed two independent WGD events (red stars) during avocado’s evolutionary history. **B**. Hass avocado (bottom 4 genomic blocks) shows 4:1 intercalated syntenic relationships with *Amborella* (upper block). **C**. Syntenic homologs in avocado show a bimodal K_s_ distribution suggestive of 2 polyploidy events (numbered 1 and 2; cyan: Hass:Hass paralogs; green: Hass:*drymifolia* homologs) following the split between magnoliids and *Amborella* (red syntenic homologs). These events postdate the species split between *Vitis* and avocado (purple syntenic homologs) and so are independent of the gamma triplication that underlies *Vitis*.

### Phylogenomic Analysis of Avocado’s Placement Among Angiosperms

To further corroborate the placement of these two polyploidy events as specific to the avocado lineage, we generated phylogenetic trees of representatives of the major angiosperm lineages using two data forms: coding sequence alignments, and modal distances within large collections of syntenic orthologs between species pairs (Supplementary information section 7).

Single-copy gene families (presumed unambiguous orthologs, those that returned to single copy following duplicate deletions after the various polyploidy events in flowering plant history (25)) were retrieved from orthogroup classification of 19 angiosperm proteomes, including those of avocado, *Amborella*, and representatives of monocots and eudicots (Table S10; Supplementary information section 3.1). Phylogenetic trees based on 176 stringently-filtered single-copy gene alignments (Supplementary information section 7.1 and supplementary data 3) gave different results for amino acid versus inferred codon data. Based on protein sequences, avocado was resolved as sister to monocots plus eudicots (i.e., branching before their divergence from each other; cf. (26, 27)), whereas from coding sequences, avocado was placed as sister to monocots only (cf. (28)) (Figs. S40, S41, respectively). In a different analysis we included *Gnetum* (a gymnosperm) and *Selaginella* (a non-seed plant) in orthogroup classification to generate a rooted species tree from all gene trees (4,694) that contained one or more (i.e., paralogous) gene copies from all species (Supplementary information section 7.2). Here, avocado was resolved as sister to eudicots only (Fig. S42), a result similarly found in transcriptome-based analyses of large numbers of species (24, 29). In an altogether different approach (30, 31), we generated a neighbor-joining tree based on modal dissimilarity scores from thousands of syntenically-validated ortholog pairs generated by the SynMap function on the CoGe platform (19) (Supplementary information section 7.3). Here, avocado was again placed as sister to monocots plus eudicots, as in Fig. 3A (Fig. S44).

Apparently, the early branching orders of the angiosperms are extremely difficult to determine using protein coding sequences. This problem is due in part to sequence parallelism/reversal over deep time, limitations in taxon sampling (including unknown extinctions), biases in sequence-based ortholog versus paralog determination, but clearly also to the relatively coincident branching times of the species involved (see (32), their Fig. 6, and also below). Rapid species divergences can lead to real gene-tree/species-tree discordances through enhanced occurrence of incomplete lineage sorting (ILS), wherein polymorphic allele states in ancestral populations do not have enough time to fix according to the species tree %(33-35).

In an experimental approach to the problem, we further investigated the possible role of ILS using gene family turnover analysis as incorporated in BadiRate (36) (Supplementary information section 7.4). Trees with the three alternative placements of avocado were converted into time-calibrated ultrametric trees, and the likelihoods of duplicate gains versus losses were evaluated under four different branch models (Supplementary information section 7.4). The AIC clearly favored Free-Rates (FR) models, supporting heterogeneous rates of multi-gene family evolution across lineages (Table S17). Interestingly, such uneven rates of gene turnover cannot be entirely explained by lineage-specific WGD/whole genome triplication (WGT) events, given that FR models fit multi-gene family data better than WGD/WGT models alone. Additionally, allowing turnover rates to vary in each short branch (<10 my) also improved likelihood and AIC values, although the fit was still worse than under the FR model (Table S17). That FR models fit gene count data significantly better could be explained by their flexibility to accommodate variation that is not explicitly accounted for by current turnover models, such as gene copy variation within species. Intraspecific variation, segregating in an ancestral population, can be inherited differently by two splitting lineages, which will thus start diverging with a significant fraction of differentiation. This predicts that divergence will be inflated for short branches, and that this bias will become negligible as divergence times increase, because its relative contribution to the total divergence tends to be comparatively small over time (37, 38). We observe a correlation between turnover rates and branch lengths at the multigene family level (Fig. S46), suggesting pervasive copy number variation (CNV) in the ancestral species, possibly exacerbated by WGD and subsequent fractionation processes. Short phylogenetic branches, representing rapid speciation events, increase the incidence of ILS in phylogeny reconstruction since extinctions of alternative duplicate copies within ancestral populations (e.g., unfixed CNVs) further break up branches that are nearly time-coincident already (39). According to BadiRate estimates, the temporal impact of ILS on turnover rates extended well beyond 10 Mya (Table S18, Fig. S46), a time-frame exceeding the branch length of the lineage that existed immediately prior to avocado divergence from other species, which varied in age from only 7.4 to as little as 3.8 million years (Table S18; Figs. 3A, S46, S47). This implies that the 3 different placements of avocado among angiosperms may be impossible to discriminate among for purely biological reasons (cf. (40)). Yet, one of the three different tree topologies was preferred based on AIC contrasts under the FR model: the topology wherein magnoliids are sister to monocots plus eudicots (Figs. 3A, S47).

### Functional Enrichments in Duplicate Gene Space

Duplicate gene collections within plant genomes mainly derive from two processes, local and ongoing tandem duplication events, many of which may be recent, and global and often ancient polyploidy events wherein entire gene complements are duplicated (41). Subfunctionalization and/or neofunctionalization of duplicate gene copies (42) results in retained descendants of duplication events that have differentially escaped the otherwise usual fate of duplicates – pesudogenization – through functional divergence. Tandem duplication is problematic for genes part of dosage-sensitive transcriptional regulatory networks, or for genes that code for parts of multiprotein complexes (43). Such functions are more likely to remain among the surviving duplicate complements stemming from precisely dosage-balancing polyploidy events (44). On the other hand, dosage responsive functions such as secondary metabolism (including biochemical pathway addition) are among those most likely to survive as sub- or neofunctionalized tandem duplicates (43). These patterns have been repeatedly observed among plant genomes, wherein secondary metabolic function is most prevalent among tandems, and transcriptional function is enriched among polyploid duplicates (e.g., (44, 45)). The avocado genome provides no exception to this rule; we identified precisely these overrepresentation patterns among GO and KEGG categories for these different classes of gene duplicates, separated using the CoGe platform (Supplementary information section 8). Among 2433 total polyploid duplicates, ‘regulation of transcription, DNA-templated’ was significantly overrepresented by 352 genes (Table S20). Enriched functions among tandem duplicates was highly illustrative of the secondary metabolic landscape particular to avocado (Table S20). We show that ‘phenylpropanoid biosynthesis’ and closely related KEGG pathways (Table S21) are significantly enriched among tandem duplicates (*p* = 2.08e-08; Fisher’s exact test, Bonferroni corrected). This functional enrichment in a long-lived tree may have evolved in response to pathogen infection, including *Colletotrichum* (anthracnose) and *Phytophthora cinnamomi* (avocado root rot), both of which are reported to activate the phenylpropanoid biosynthetic pathways in avocado %(46-48). Several GO functional enrichments among avocado tandems (for example, ‘1,3-beta-D-glucan synthase activity’ and ‘regulation of cell shape’; *p* = 1.64e-05 and 0.00258, respectively; Fisher’s exact test, Bonferroni corrected) relate to callose synthase activity (49, 50), a recently discovered avocado defense mechanism against *P. cinnamomi* (51). Other significantly enriched GOs include ‘phenylpropanoid metabolic process‘, ‘lignin biosynthetic process’ and ‘UDP-glycosyltransferase activity’ (*p* = 0.00142, 7.36e-07 and 5.16e-07, respectively; Fisher’s exact test, Bonferroni corrected,), categories directly or closely related to phenylpropanoid biosynthesis (52, 53). The lignin functional enrichment, for example, includes diverse tandemly duplicated genes involved in many pathway-interrelated processes, including homologs of both biosynthetic and regulatory genes encoding HYDROXYCINNAMOYL-COA SHIKIMATE/QUINATE HYDROXYCINNAMOYL TRANSFERASE (HCT), CINNAMYL ALCOHOL DEHYDROGENASE 5 (CAD5), LACCASE 17 (LAC17), CAFFEATE O-METHYLTRANSFERASE 1 (COMT1), PEROXIDASE 52 (PRX52), NAC DOMAIN CONTAINING PROTEIN 12 (NAC012), and NAC SECONDARY WALL THICKENING PROMOTING FACTOR 1 (NST1). As could be expected from the above, the GOs ‘defense response’ and ‘defense response to fungus’ are significantly enriched among tandem duplicates (*p* = 0.000165 and 0.0167, respectively; Fisher’s exact test, Bonferroni corrected), as has been discovered for other plant genomes, and involving many different gene families and responses. Tandem O-methyltransferases homologous to COMT1 may also contribute to synthesis of the phenylpropanoid derivative and insecticide estragole (54), which is largely responsible for the anise-like leaf scent and fruit taste of many avocado cultivars, particularly of the Mexican race (55). Another relevant enriched GO category among tandems is ‘ethylene-activated signaling pathway’ (*p* = 0.000463; Fisher’s exact test, Bonferroni corrected), which annotates many different transcription factor duplicates. Ethylene signaling factors such as ERF1 (represented by 2 homologs) are heavily involved in pathogen-induced responses, including to infection by *Colletotrichum* and other necrotrophic fungi %(56-59). Also identified are 3 homologs of EIN3, a transcription factor that initiates downstream ethylene responses, including fruit ripening (60). Avocado fruit matures on the tree in a process that involves ethylene synthesis and signaling, while it does not ripen until harvested - a desirable trait that allows growers to delay harvesting for several months (61).

Given the ancient derivation of avocado’s retained polyploid duplicates, most tandem duplicates in the genome are expected to be of more recent origin, having been generated by ongoing gene birth-death-innovation processes that operate in all eukaryotic genomes. As such, sub- or neofunctionalized tandem duplicates that survive the usual fate of duplicated genes – pseudogenization – should be enriched in functions that fine-tune a given species’ recent selective environment. In the case of avocado, response to fungal pathogens is precisely reflected in its tandemly duplicated gene complement.

### Differential Expression of Tandem Versus Polyploid Duplicates

Following our prediction that many tandem duplicates fixed in the avocado genome may have evolved under relatively recent pathogen pressure, we examined differential expression of Hass genes after treatment with the anthracnose causal agent (62) (Supplementary information section 9). Hass transcriptome reads for untreated control vs. pathogen-treated were mapped to Hass gene models using Kallisto (63), normalized to transcript-per-million (TPM) values and thresholded by identifying genes with treatment/control log2 fold-change outside of the [2,-2] interval. Tandems were significantly enriched among upregulated (*p* = 3.536e-09; Fisher exact test) and downregulated genes (*p* = 7.274e-07), whereas polyploid duplicates did not show enrichment (Supplementary information section 9). We interpret these results to indicate that tandem duplicates are the most dynamic component of the avocado duplicate gene space under pathogen treatment.

We also examined functional enrichments for up-vs. downregulated tandem duplicates (Table S24). The only significantly enriched category was xyloglucan:xyloglucosyl transferase activity (*p* = 0.038984; Fisher’s exact test, Bonferroni corrected). Among genes with this annotation were 4 homologs of *XYLOGLUCAN ENDOTRANSGLUCOSYLASE/HYDROLASE 22* (*XTH22*; also known as *TOUCH 4* (*TCH4*) (64)). *XTH22* and similar genes encoding cell-wall modifying proteins have been shown to up- or down-regulate after Citrus Huanglongbing infection (65), whitefly infestation (63), and herbivore (66) or mechanical stimulation (67), the latter provoking a *Botrytis*-protective response. In a different pathogen response, upregulation of *XTH22* occurs in concert with pectin digestion in *Pseudomonas*-sensitive Arabidopsis lines that overexpress *IDA-LIKE 6* (*IDL6* (68)).

## CONCLUSIONS

Our new genomes of Mexican and Hass avocados provide the requisite resources for genome-wide association studies to identify important traits among natural avocado genetic diversity present in Mesoamerica, to develop genome-assisted breeding and genetic modification efforts crucial for the improvement of this long life-cycle crop, to fight threatening avocado diseases and optimize growth and desirable phenotypic traits. We anchored almost half of the sequenced Hass genome to a genetic map, providing linkage information for genetic variation on 12 chromosomes. We resequenced 10 genomes representing small populations of Guatemalan, West Indian, Mexican and Hass-related cultivars – and the genome of the closely related species *Persea schiedeana* – in order to call SNPs and study genetic diversity among these chromosomes. Analyses of admixture and introgression clearly highlighted the hybrid origin of Hass avocado, pointed to its Mexican and Guatemalan progenitor races, and showed Hass to contain Guatemalan introgression in approximately one-third of its genome. Introgressed blocks of chromosome arm size matched expectation based on Hass’s recent (20^th^ Century) origin. We uncovered two ancient polyploidy events that occurred in the lineage leading to avocado, and conclude that these were independent from genome duplications or triplications known to have occurred in other angiosperm clades. We contributed to solving the problem of magnoliid phylogenetic relationships to other major angiosperm clades by showing that thousands of syntenic orthologs among 14 species support an arrangement wherein the magnoliid clade branched off before the split between monocots and eudicots. However, this resolution is tentative, with coding sequence phylogenomics inconclusive and gene family birth/death analysis suggesting appreciable duplicate gene turnover – and therefore enhanced possibility for incomplete lineage sorting - during what appears to have been a nearly coincident radiation of the major angiosperm clades. We also studied the adaptive landscape of the avocado genome through functional enrichment analyses of its mechanistically-distinct duplicate gene collections, i.e., tandem vs. polyploid duplicates. Tandem duplicates were enriched with many potentially important metabolic responses that may include relatively recent adaptation against fungal pathogens. In contrast, ancient polyploid duplicates, which originated in two distinct waves, were enriched with transcriptional regulatory functions reflective of core physiological and developmental processes. We discovered that tandem duplicates were more dynamically transcribed following anthracnose infection, and that some of the upregulated genes could be related to defense responses. In sum, our work paves the way for genomics-assisted avocado improvement (1).

## Data availability

Bioproject: PRJNA508502. Biosamples: SAMN10523735; SAMN10523720; SAMN10523736; SAMN10523738; SAMN10523739; SAMN10523746; SAMN10523747; SAMN10523748; SAMN10523749; SAMN10523750; SAMN10523752; SAMN10523753; SAMN10523756. SRA submission: SUB4878870. Whole Genome Shotgun projects have been deposited at DDBJ/ENA/GenBank under the accessions SDXN00000000 and SDSS00000000. The versions described in this paper are version SDXN01000000 and SDSS01000000 (*P. americana* var. *drymifolia* and *P. americana* cultivar Hass, respectively). The genome assemblies and annotations are available at https://genomevolution.org/CoGe/SearchResults.pl?s=29305&p=genome and https://genomevolution.org/coge/SearchResults.pl?s=29302&p=genome (*P. americana* var. *drymifolia* and *P. americana* cultivar Hass, respectively).

## Acknowledgements

This project was funded in part by a grant from the SAGARPA/CONACyT sectorial program to LHE (Grant 00126261), grants 0922742 and 1442190 to VAA from the U.S. National Science Foundation, as well as (to NM and AH) Horticulture Innovation Australia Ltd (HIA) and the Australian Bureau of Agricultural and Resource Economics and Sciences (ABARES; Australian Government). We thank the Fundación Salvador Sánchez Colín-CICTAMEX, S.C. for providing some avocado specimens. We also thank Araceli Fernandez and Emanuel Villafan, administrators of the high-performance computing systems at LANGEBIO and INECOL, respectively.

## Author contributions

LHE conceived research; LHE, VAA and AHE led research; EIL, AMB, CAPT, ACL, GHG, NM, AH, SF, DK, and AFBP provided materials and/or collected data; MRA, EIL, TL, CZ, LCP, THC, KMF, WBB, SC, MM, DSh, JR, ASG, DK, JS, PL, DSa, and VAA analyzed data; MRA, EIL, PL, DSa provided text for the supplement; and VAA and LHE wrote the main text. All authors approved the final manuscript.

## Footnotes

* https://www.statista.com/topics/3108/avocado-industry/ and http://www.freshplaza.com/article/178230/Mexico-Avocado-exports-generate-2.5-billion-dollars

## Notes

#### Summary of Updates

Figure 3C and caption revised. Missing Supplementary text and figures PDF added.

