## Supplementary information (text, figures, tables) for "The Avocado Genome Informs Deep Angiosperm Phylogeny, Highlights Introgressive Hybridization, and Reveals Pathogen-Influenced Gene Space Adaptation"

### 1. Sequencing and assembly of the *Persea americana* genomes

#### 1.1. Plant material

Plant material for the *P. americana* var. *drymifolia* (001-01) reference genome and a second *drymifolia* resequenced individual were obtained from the germplasm bank of the 'Instituto Nacional de Investigaciones Forestales y Agropecuarias (INIFAP)' in Uruapan, Michoacan, Mexico. The remaining resequenced accessions (including the outgroup species *P. schiedeana*) were obtained from the 'Fundación Salvador Sánchez Colín (CICTAMEX, S.C) high altitude *Persea* germplasm bank located at La Cruz, Experimental Center at Coatepec Harinas in the State of Mexico. The materials for the Hass reference genome and Carmen Hass cultivar were collected from a commercial orchard in Tingambato, Michoacan, Mexico. Finally, 'Velvick' rootstock was provided by the University of Queensland, Australia. (Table S1,2).

#### 1.2. Flow cytometric analysis and genome size estimation

Genome size for *P. americana* was assessed using flow cytometry. Young leaves were finely chopped with a razor blade in Petri dishes with 500µL of nuclei extraction buffer (Cystain ultraviolet Precise P Nuclei Extraction Buffer; Partec GmbH, Münster Germany). The suspension was passed through Partec Cell Trics disposable filters with a pore size of 50 µm. The nuclei were stained with 1.5 mL of 4,6-diamidino-2 phenylindole (DAPI). DNA content of at least 5,000 stained nuclei was determined for each sample using a PARTEC CA II Cytometer (Partec GmbH, Münster Germany) after UV excitation of DAPI. *Arabidopsis thaliana* (2C = 0.39 pg) and *Pisum sativum* (2C = 9.09 pg) were used as internal standards (1, 2). Using the conversion factor of 1 pg DNA = 978 Mbp (3), the genome size of *P. americana* was estimated to be 980 Mb.

Previous flow cytometric estimates suggested that the genome of *P. americana* is approximately 911 Mb (4) while our assemblies for the Hass cultivar and var. *drymifolia* suggest sizes of 823.4 Mb and 912.7 Mb, respectively (section 1.4). Analysis of k-*mer* frequencies (5) suggested a genome size of 937-951 Mb. Thus, sequence-based methods suggest that the *P. americana* genome size is closer to 920 Mb, which represents an average of these three estimation approaches.

#### 1.3. High molecular weight DNA preparation, library construction and sequencing

Prior to sequencing, the genetic background of the trees was confirmed using a subset of 9 EST-SSR markers previously described (6). Due to their high polymorphism, these EST-SSR markers are sufficient to clearly distinguish botanical varieties and cultivars such as Hass. The *Persea americana* var. *drymifolia* reference individual (001-01) was selected to minimize heterozygosity. In order to minimize chloroplast and mitochondrial DNA contamination, high molecular weight DNA was prepared from nuclei of young leaves according to previously described protocols (7). First, isolated nuclei were collected from a 60% Percoll (Invitrogen) density gradient following low-speed centrifugation (4000g for 10 min at 4°C), then high-quality megabase-sized DNA was isolated (8) and sheared (Covaris® M220 Focused-ultrasonicator™) to obtain DNA fragments ranked according to the size required for sequencing libraries (~0.5 Kb, 1 Kb, 3 Kb, 5 Kb, or 8 Kb). DNA libraries were constructed and sequenced (from one end [single-reads] or from both ends [paired-end reads]) using the 454 (Roche) and HiSeq (Illumina) platforms at the Genomic Services Laboratory of LANGEBIO-CINVESTAV, Mexico. A BAC library was constructed at the Arizona Genomic Institute following established methods (9). Briefly, DNA was isolated from nuclei collected in agarose plugs and DNA digestion was performed with HindIII, followed by ligation to the pAGIBAC1 vector (a modified pIndigoBAC536Blue with an additional SwaI site (10)). BAC genomic insert sizes were estimated by *Not*I digestion and pulsed-field gel electrophoresis (CHEF-DRIII system, Bio-Rad) as described elsewhere (9). Using a genome-wide shotgun strategy, a total of 472.6 million 454 single and paired-end reads and 55,824 Sanger BAC-end reads were generated, representing ~186x coverage of the 920-Mb genome (see section 1.2). Additional Illumina sequencing data (~31x) were used to improve the assembly (Table S3). Sequence reads were mainly assembled with Newbler v2.6 (see section 1.4).

For the reference genome of the Hass cultivar, high-quality megabase-sized DNA isolated from nuclei of young expanding leaves was submitted to the National Center for Genome Resources (NCGR) for PacBio single-molecule real-time (SMRT) sequencing. A single library was prepared and run on 102 SMRT cells. With a genome size of approximately 920 Mb, PacBio SMRT sequencing provided approximately 80x coverage of the entire genome. SMRT sequencing of the Hass cultivar genome initially resulted in 5.5 million raw reads, with a mean read length of 13.8 Kb, totaling 75.3 Gb.

Finally, from each of the individuals selected for resequencing, libraries with an insert size of ~375 bp were constructed and sequenced on an Illumina HiSeq2000 sequencing machine. The paired-end reads generated from each dataset represent coverage ranged from ~3 to ~16x with respect to the avocado genome size (see section 3.2). From Hass, we generated ~80x genome coverage from two short-insert paired-end libraries (~350 bp and ~550 bp, respectively). These libraries were generated from the same tree selected to generate the reference genome, and in the assembly process were only used in the base correction step. Finally, two additional paired-end libraries (~39x coverage each) were made by pooling equimolar amounts of DNA isolated from four different trees of Hass and Carmen Hass cultivars (Table S2). This pool of individuals was included as an additional Hass sample – labeled as Hass2 – in downstream analyses (i.e., phylogenetic reconstruction and population genomics).

Finally, in the case of the Velvick rootstock cultivar, genomic DNA was extracted (11) from clone A998 maintained at the Maroochy Research Facility, Queensland Government Department of Agriculture and Fisheries. Illumina shotgun DNA libraries with Pippin Prep size selection (250 bp insert) were sequenced (HiSeq 2x100 bp PE) at the Australian Genome Research Facility (AGRF).

Table S1: List of sequenced accessions of *Persea schiedeana* and *Persea americana* varieties.

| | | | **Accession** | **Number of tree** | **Collection Site** | **Common name** | **Germplasm bank** |
|---|---|---|---|---|---|---|---|
| ***Persea schiedeana* (Chinene)** | | | CH-GU-01 | 17 | Mazatenango, Suchitepequez, Guatemala | Otrabanda | Fundacion Salvador Sánchez Colín (CICTAMEX, S.C); Coatepec Harinas in the State of Mexico |
| ***Persea americana*** | **Botanical varieties** | **var. *drymifolia*** | 001-01 | --- | --- | | Instituto Nacional de Investigaciones Forestales, (INIFAP); in Uruapan, Michoacan, Mexico |
| | | | 069-02 | --- | --- | | |
| | | | (Tiny-Charly) | 104 | unknown | Tiny Charly (provided by 'Colegio de Postgraduados' (CP) germplasm bank. Puebla, Mexico | Fundacion Salvador Sánchez Colín (CICTAMEX, S.C); Coatepec Harinas in the State of Mexico |
| | | **var. *guatemalensis*** | CH-G-07 | 63 | San Cristobal de las Casas, Chiapas, Mexico | SCrMer 7S1 | |
| | | | CH-G-10 | 80 | Olanca, Chiapas, Mexico | Olanca 2S3 | |
| | | | CH-G-11 | 116 | Olanca, Chiapas, Mexico | Olanca 3S1 | |
| | | **var. West Indian** | 263-C | | Hunucma, Yucatan, Mexico | Hunucmá 09 | |
| | **Wild** | **var. *costarricensis*** | CH-CR-25 | 105 | Matapalo, Puntarenas, Costa Rica | Las Nubes 06 | |
| | **Cultivars** | **Commercial Varieties** | Hass | --- | Tingambato, Michoacan, Mexico | Hass | Commercial orchard; Tingambato, Michoacan, Mexico |
| | | | Mendez | --- | | Carmen/ Hass (Mendez) | |
| | **Phytophthora-tolerant rootstock** | | --- | --- | University of Queensland, Australia | Velvick | --- |

Table S2: Summary of *Persea* resequencing data.

| Species | Group | Subgroup | Variety | Accession | Number of Illumina paired-end reads | Estimate of paired-end distance (derived from mapping process) | Standard deviation of estimated distances |
|---|---|---|---|---|---|---|---|
| ***Persea schiedeana* (Chinene)** | | | | CH-GU-01 | 104,521,517 | 369.10 | 87.61 |
| ***Persea schiedeana* (Chinene)** | | | | CH-GU-01 | 62,917,160 | 508.85 | 159.11 |
| ***Persea americana*** | **Botanical varieties (Horticultural races)** | | **var. *drymifolia* (Mexican race)** | 069-02 | 140,485,158 | 290.23 | 62.93 |
| ***Persea americana*** | **Botanical varieties (Horticultural races)** | | **var. *drymifolia* (Mexican race)** | (Tiny Charly) | 120,886,270 | 420.42 | 111.83 |
| ***Persea americana*** | **Botanical varieties (Horticultural races)** | | **var. *guatemalensis* (Guatemalan race)** | CH-G-07 | 97,086,017 | 305.35 | 66.65 |
| ***Persea americana*** | **Botanical varieties (Horticultural races)** | | **var. *guatemalensis* (Guatemalan race)** | CH-G-10 | 76,413,139 | 280.31 | 66.33 |
| ***Persea americana*** | **Botanical varieties (Horticultural races)** | | **var. *guatemalensis* (Guatemalan race)** | CH-G-11 | 99,872,804 | 293.31 | 76.32 |
| ***Persea americana*** | **Botanical varieties (Horticultural races)** | | **var. *americana* (West Indian race)** | 263-C | 76,188,605 | 309.76 | 67.81 |
| ***Persea americana*** | **Wild type** | | **var. *costaricensis*** | CH-CR-25 | 74,918,362 | 269.54 | 70.42 |
| ***Persea americana*** | **Commercial varieties** | **Cultivars** | **cv. Hass (Guatemalan x Mexican race hybrid)** | Hass | 207,829,207 | 396.11 | 97.21 |
| ***Persea americana*** | **Commercial varieties** | **Cultivars** | **cv. Hass (Guatemalan x Mexican race hybrid)** | Hass | 176,532,977 | 577.09 | 154.99 |
| ***Persea americana*** | **Commercial varieties** | **Cultivars** | **cv. Hass (Guatemalan x Mexican race hybrid)** | Hass | 179,112,055 | 423.06 | 103.58 |
| ***Persea americana*** | **Commercial varieties** | **Cultivars** | **cv. Carmen/Hass** | Carmen/Hass (Mendez) | 177,401,667 | 417.95 | 103.82 |
| ***Persea americana*** | **Commercial varieties** | **Rootstock** | **Velvick (West Indian × Guatemalan race hybrid)** | Velvick | 254,083,516 | | |
| ***Persea americana*** | **Commercial varieties** | **Rootstock** | **Velvick (West Indian × Guatemalan race hybrid)** | Velvick | 258,697,634 | | |

Table S3: Summary of sequencing data used in *Persea americana* var. *drymifolia* genome assembly.

| Type of sequences \| platform | High quality reads[1] | Average length | Number of bases | Coverage[2] (Mb) | Estimate of paired-end distance (derived from assembly process) | Standard deviation of estimated distances |
|---|---|---|---|---|---|---|
| BAC-ends (120K) \|3730xl AB | 55,824 (x2) | 595.64 | 66,502,015 | 0.08 | 114,276.50 | 28,569.10 |
| Single-ended \| 454-$XLR_{plus}$ | 11,226,071 | 609.49 | 6,842,207,104 | 8.31 | NA | NA |
| Single-ended \| 454-$FLX_{titanium}$ | 31,749,065 | 356.33 | 11,313,133,140 | 13.75 | NA | NA |
| Paired-end (3Kb) \| 454-$FLX_{titanium}$ | 63,133,796 | 356.59 | 22,513,183,417 | 27.36 | 2,124.80 | 531.2 |
| Paired-end (5Kb) \| 454-$FLX_{titanium}$ | 124,066,522 | 355.81 | 44,144,221,067 | 53.64 | 5,721.70 | 1,430.40 |
| Paired-end (8Kb) \| 454-$FLX_{titanium}$ | 242,431,346 | 355.28 | 86,131,581,833 | 104.66 | 8,482.58 | 2,145.98 |
| Paired-end (650 bp)\| HiSeq2000 | 101,288,690 (x2) | 100 | 20,257,738,000 | 24.61 | 550.00 | 275 |
| Paired-end (1Kb) \| HiSeq2000 | 62,080,646 (x2) | 100 | 12,416,129,200 | 15.09 | 950.00 | 475 |
| Single-ended \| HiSeq2000 | 121,180,202 (x1) | 100 | 12,118,020,200 | 14.72 | NA | NA |
| **Total** | | | **215,802,715,976** | **262.21** | | |

[1] Selected based on the stringency parameters: -q 30 (Minimum quality score to keep), -p 95 (Minimum percent of bases that must have [-q] quality) and -a 30 (the average quality of paired-end or single-ended reads).
[2] Considering the final Newbler assembly size of 823 Mb.

### 1.4. Genome assembly

#### *P. americana* var. *drymifolia*

454 reads originating from chloroplast and mitochondrial genomes were filtered from all sequence data prior to assembly. Additionally, using the CD-HIT pipeline (12) reads were screened to remove artificial duplicates originating from PCR errors. Filtered 454 reads and Sanger BAC ends were assembled using Newbler v2.6. From the initial 472.6 million reads, about 80.20% were assembled. The resulting 106,547 contigs were assembled and linked into 50,037 scaffolds. In addition, using Illumina single and paired-end reads, the generated assembly was subsequently scaffolded and gap-closed using SSPACE (13) and GapFiller (14), respectively. The resulting assembly consists of 100,563 contigs merged

into 42,722 scaffolds, spanning 823.4 Mb including embedded gaps (18.82%). The contig N50 was 11.7 Kb, and the scaffold N50 was 323.8 Kb (Table S4). The cumulative scaffold size was about 10% smaller than the estimated genome size of 920 Mb (section 1.2). The final assembly was corrected using iCORN (15), which was run to correct single base and short indel errors through re-alignment of the Illumina reads.

***P. americana*, Hass cultivar**
PacBio long reads were assembled using the FALCON assembler (https://github.com/PacificBiosciences/FALCON). The resulting assembly consists of 8,135 contigs (≥ 2Kb), spanning 912.6 Mb, which represents 99.2% of the estimated genome size. The N50 length was 2.38 Mb across 770 contigs (Table S4). The per-base error rate of the *de novo* assembly was subsequently reduced using iCORN software (15) through of the alignment of the Illumina read pairs from the 350 bp an 550 bp libraries.

To assess and compare the quality of the assemblies, similar metrics and statistics were calculated using the Assemblathon software (16). Using the PacBio technology we produced an assembly with 815 contigs smaller than 10 Kb (4.9 Mb), while the hybrid strategy used for the *drymifolia* variety produced a total of 35,241 (116.5 Mb). In both cases, the number of sequences longer than 10 Kb produced was similar (7,320 and 8,022, respectively), however, approximately twice more sequences up to 100 Kb were produced in the Hass cultivar assembly (Table S4). Differences between the assembled genomes of Hass and *drymifolia* are not limited only to the number and size of the sequences produced, because the absence of gaps in the PacBio contigs (even with significantly lower coverage of 80x) provides a higher quality assembly.

Table S4. Statistics of assemblies of *P. americana* genomes. Statistics and metrics were calculated with Assemblathon software.

| | ***P. americana* genome** | |
|---|---|---|
| | **var. *drymifolia*** | **Hass** |
| Number of scaffolds | 42,722 | NA |
| Total size of scaffolds | 823,419,498 | NA |
| Longest scaffold | 4,610,966 | NA |
| Shortest scaffold | 1,712 | NA |
| Number of scaffolds > 1K nt | 42,722 (100%) | NA |
| Number of scaffolds > 10K nt | 7,890 (18.5%) | NA |
| Number of scaffolds > 100K nt | 1,143 (2.6%) | NA |
| Number of scaffolds > 1M nt | 121 (0.3%) | NA |
| Mean scaffold size | 19,274 | NA |
| Median scaffold size | 2,987 | NA |
| N50 scaffold length | 323,854 | NA |
| L50 scaffold count | 502 | NA |
| scaffold %A | 24.73 | NA |
| scaffold %C | 15.73 | NA |
| scaffold %G | 15.77 | NA |

| | | |
|---|---|---|
| scaffold %T | 24.90 | NA |
| scaffold %N | 18.87 | NA |
| scaffold %non-ACGTN | 0.00 | NA |
| Number of scaffold non-ACGTN nt | 0 | NA |
| Percentage of assembly in scaffolded contigs | 87.60 | 0.00 |
| Percentage of assembly in unscaffolded contigs | 12.40 | 100.00 |
| Average number of contigs per scaffold | 2.30 | 1.00 |
| Average length of break (>25 Ns) between contigs in scaffold | 2,712 | 0 |
| Number of contigs | 99,957 | 8,135 |
| Number of contigs in scaffolds | 67,163 | 0 |
| Number of contigs not in scaffolds | 32,794 | 8,135 |
| Total size of contigs | 668,137,248 | 912,697,600 |
| Longest contig | 254,240 | 2,811,280 |
| Shortest contig | 500 | 2,013 |
| Number of contigs > 1K nt | 98,149 (100%) | 8,135 (100%) |
| Number of contigs > 10K nt | 17,143 (17.1%) | 7,320 (90.0%) |
| Number of contigs > 100K nt | 82 (0.1%) | 2,157 (26.5%) |
| Number of contigs > 1M nt | 0 | 84 (1.0%) |
| Mean contig size | 6,684 | 112,194 |
| Median contig size | 3,438 | 44,946 |
| N50 contig length | 11,724 | 296,371 |
| L50 contig count | 14,226 | 770 |
| contig %A | 30.48 | 30.43 |
| contig %C | 19.38 | 19.54 |
| contig %G | 19.44 | 19.57 |
| contig %T | 30.69 | 30.46 |
| contig %N | 0.00 | 0.00 |
| contig %non-ACGTN | 0.00 | 0.00 |
| Number of contig non-ACGTN nt | 0 | 0 |

### 1.5. Scaffold anchoring into pseudochromosomes

Two large mapping populations of avocado consisting of 1339 trees were genotyped with 5050 SNP markers from transcribed genes using an Illumina Infinium SNP chip (17). A Florida mapping population consisted of 527 progeny from Tonnage x Simmonds and 249 from Simmonds x Tonnage. A California mapping population consisted of 576 progeny from Hass x Bacon and 230 progeny from Bacon x Hass. Microsatellite marker data from the Florida mapping populations that had been used to produce a moderately resolved genetic recombination map (18) were included with the SNP data to anchor the new maps to the old. Individual maps were created for each population and joined using JoinMap4.1.

The resulting saturated map was then used to order the Hass scaffolds into twelve linkage groups as follows. The 3457 probes (121mers) designed for the Illumina chip were mapped against the Hass reference genome assembly using blastn with highly stringent parameters: only those probes with unique hits on the genome displaying >95% identity and 100% coverage, or, probes with no more than two hits on the genome displaying 100% identity and coverage were considered and manually curated.

The calculated genetic distances of the SNPs from each probe, together with the physical distance in the Hass scaffolds were formatted to use as input for Allmaps (v. 08.17;(19)) to reconstruct the linkage groups. Twelve pseudo-chromosomes were assembled with a correlation between genetic and physical distances of 1 (Figure S1-S12). Overall, 2688 unique markers were considered, having an average of 6.2 markers per Mb. The total length of the anchored genome accounts for 46.2% of the Hass assembly, and represents 915 scaffolds, 361 of which could be oriented, and 271 estimated gaps (Table S5-7).

Table S5. AllMaps summary for consensus map.

| | **Map** | **Anchored** | **Oriented** | **Unplaced** |
|---|---|---|---|---|
| **Linkage Groups** | 12 | | | |
| **Markers (unique)** | 2,688 | 2,643 | 1,740 | 45 |
| **Markers per Mb** | 6.2 | 6.3 | 7.1 | 0.1 |
| **N50 Scaffolds** | 548 | 537 | 294 | 233 |
| **Scaffolds** | 947 | 915 | 361 | 7,220 |
| **Scaffolds with 1 marker** | 415 | 390 | 0 | 25 |
| **Scaffolds with 2 markers** | 189 | 184 | 130 | 5 |
| **Scaffolds with 3 markers** | 111 | 111 | 40 | 0 |
| **Scaffolds with >= 4 markers** | 232 | 230 | 191 | 2 |
| **Total bases** | 430,938,895 (47.2%) | 421,428,294 (46.2%) | 243,553,840 (26.7%) | 491,269,306 (53.8%) |

Table S6. Consensus map length.

| | **Length (bp)** | | | | |
|---|---|---|---|---|---|
| Chromosome | **contigs** | **contigs + connectors** | **estimated gap length** | **ctg+gaps + connectors** | **contigs** |
| **chr1** | 62,796,911 | 62,805,911 | 3,918,359 | 66,722,070 | 91 |
| **chr2** | 55,958,243 | 55,969,643 | 6,685,340 | 62,651,183 | 115 |
| **chr3** | 49,438,558 | 49,451,358 | 8,372,500 | 57,819,958 | 129 |
| **chr4** | 21,589,134 | 21,594,134 | 2,042,400 | 23,635,334 | 51 |
| **chr5** | 42,831,826 | 42,843,126 | 5,995,465 | 48,834,791 | 114 |
| **chr6** | 26,412,953 | 26,418,753 | 2,652,552 | 29,069,405 | 59 |
| **chr7** | 27,800,660 | 27,807,660 | 4,040,497 | 31,845,957 | 71 |
| **chr8** | 32,101,031 | 32,106,731 | 2,642,167 | 34,747,698 | 58 |

| | | | | | |
|---|---|---|---|---|---|
| **chr9** | 24,970,589 | 24,977,689 | 3,887,705 | 28,862,994 | 72 |
| **chr10** | 25,753,982 | 25,758,782 | 2,170,842 | 27,927,924 | 49 |
| **chr11** | 28,371,989 | 28,377,589 | 2,755,426 | 31,131,715 | 57 |
| **chr12** | 23,402,418 | 23,407,218 | 3,753,180 | 27,158,898 | 49 |
| **Total** | **421,428,294** | **421,518,594** | **48,916,433** | **470,407,927** | **915** |

Table S7. (See additional file.) Order/position of contigs and estimated gaps in chromosomes.

Figure S1. Anchored chromosome 1. Genetic vs. physical distances and their correlation are displayed.

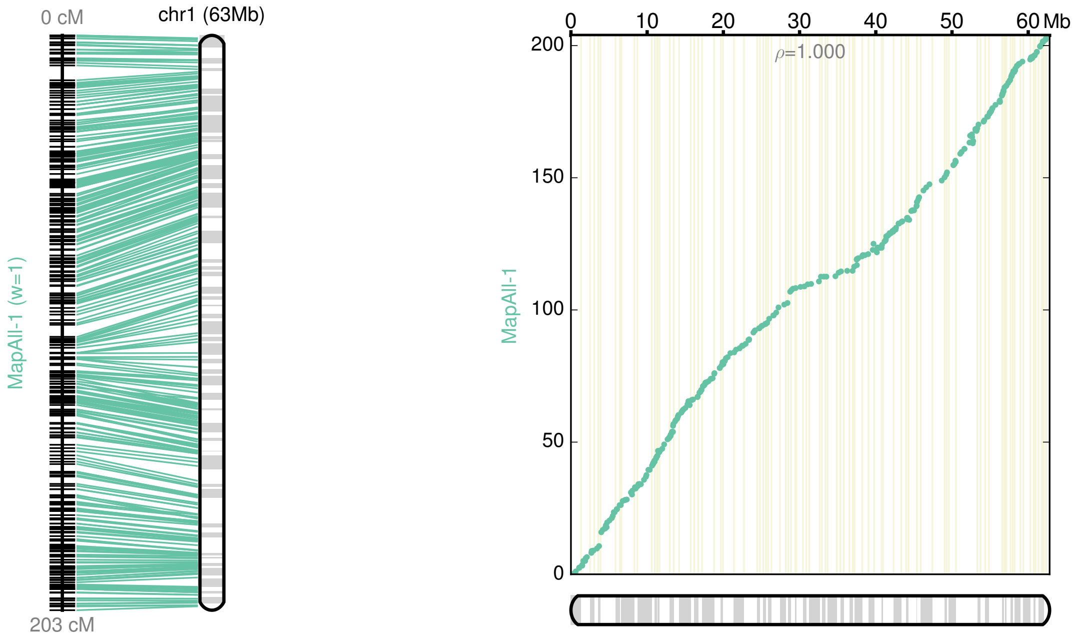

Figure S2. Anchored chromosome 2.

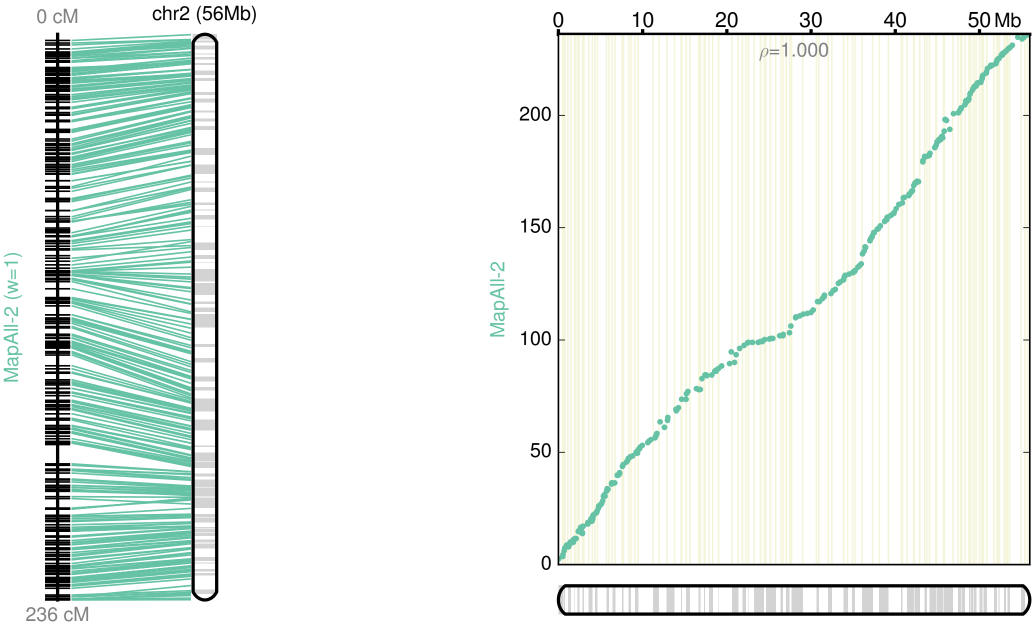

Figure S3. Anchored chromosome 3.

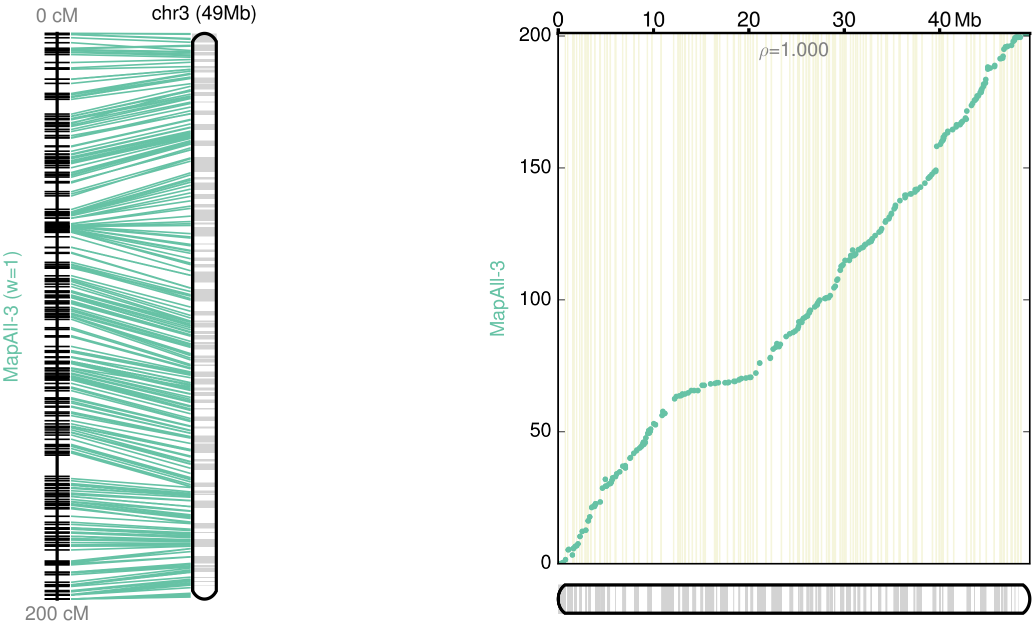

Figure S4. Anchored chromosome 4.

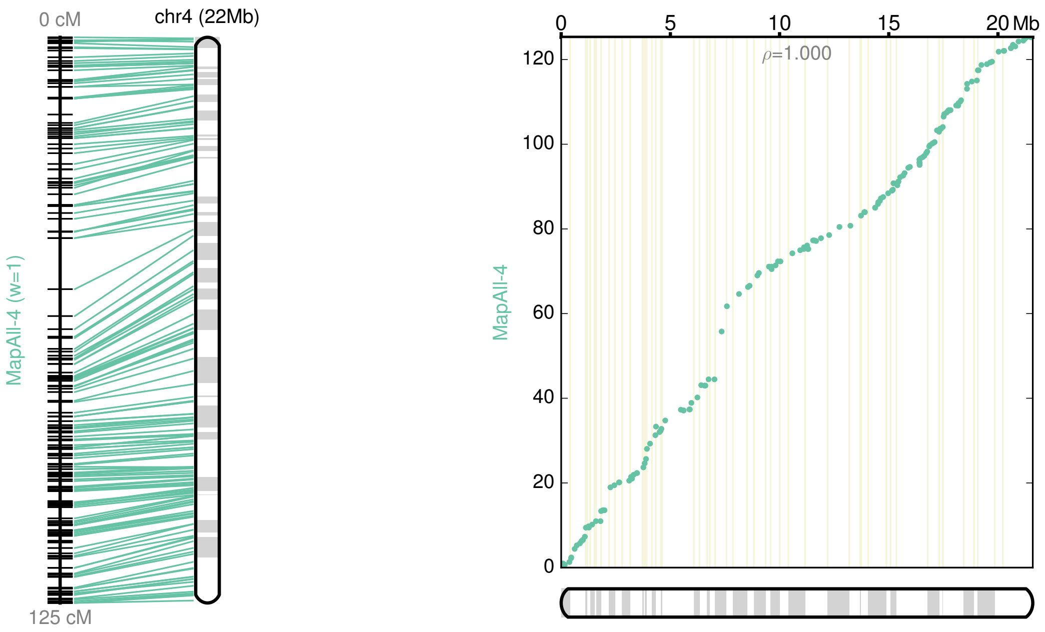

Figure S5. Anchored chromosome 5.

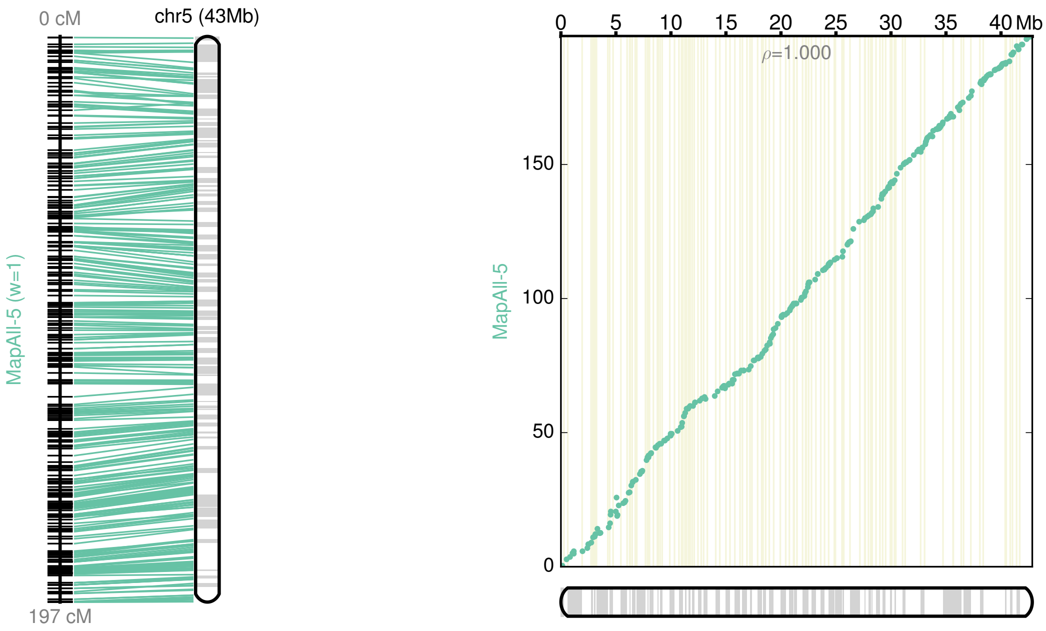

Figure S6. Anchored chromosome 6.

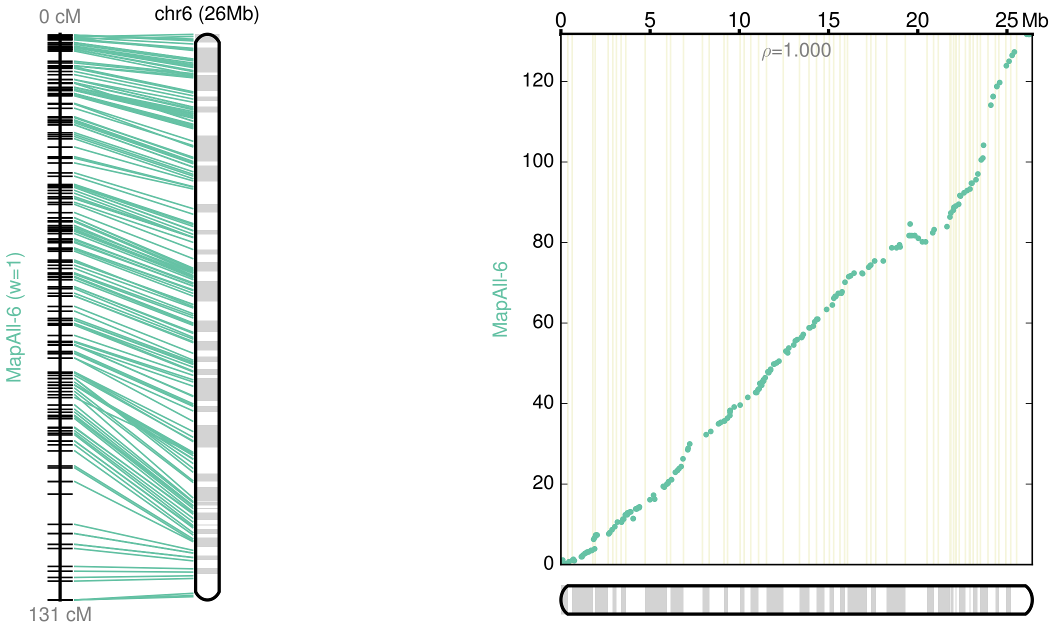

Figure S7. Anchored chromosome 7.

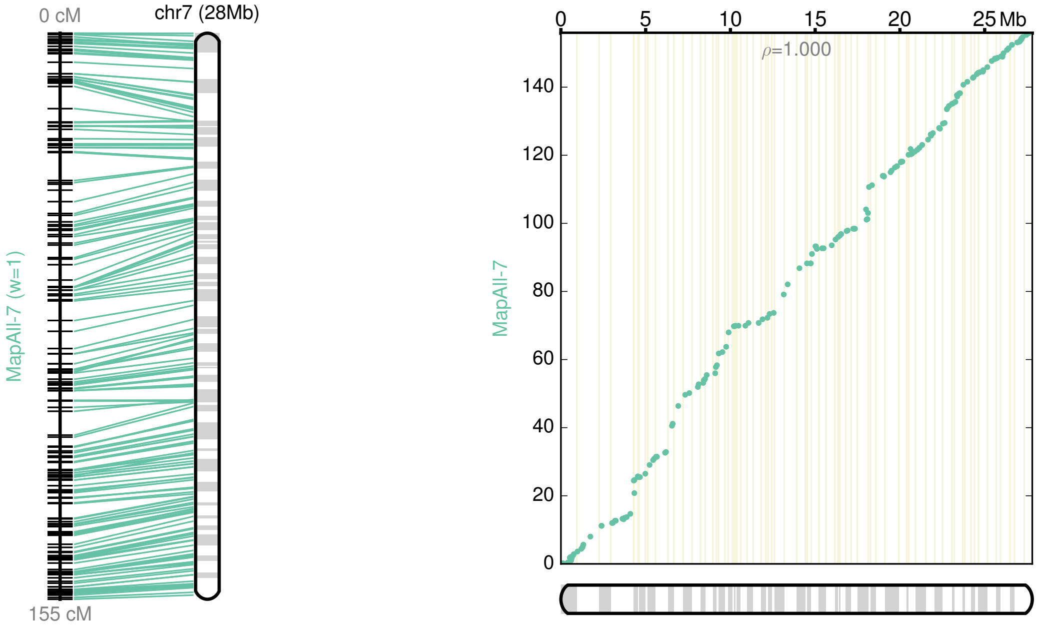

Figure S8. Anchored chromosome 8.

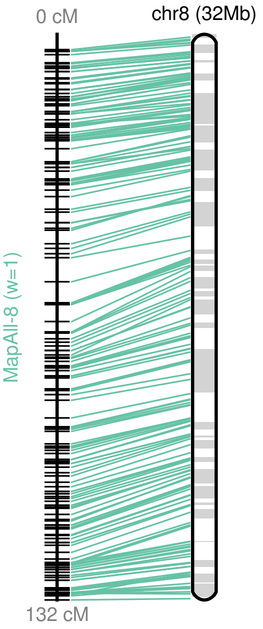

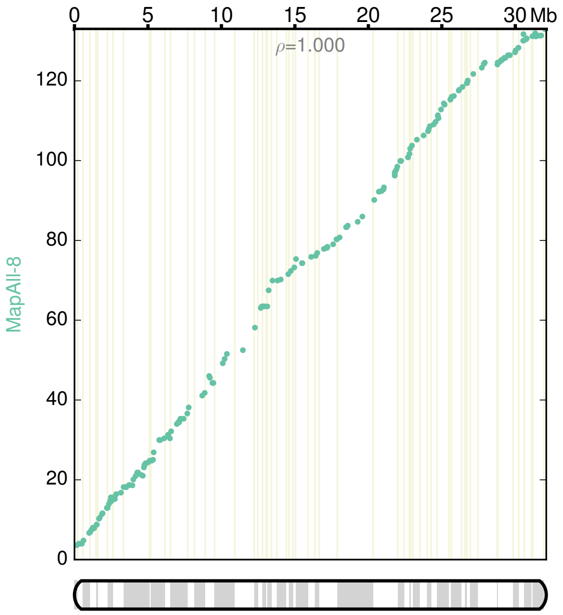

Figure S9. Anchored chromosome 9.

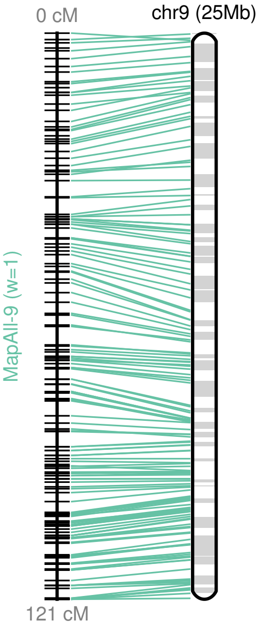

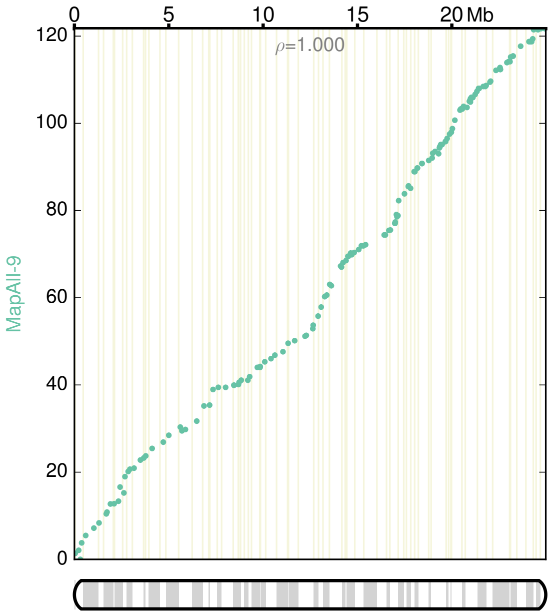

Figure S10. Anchored chromosome 10.

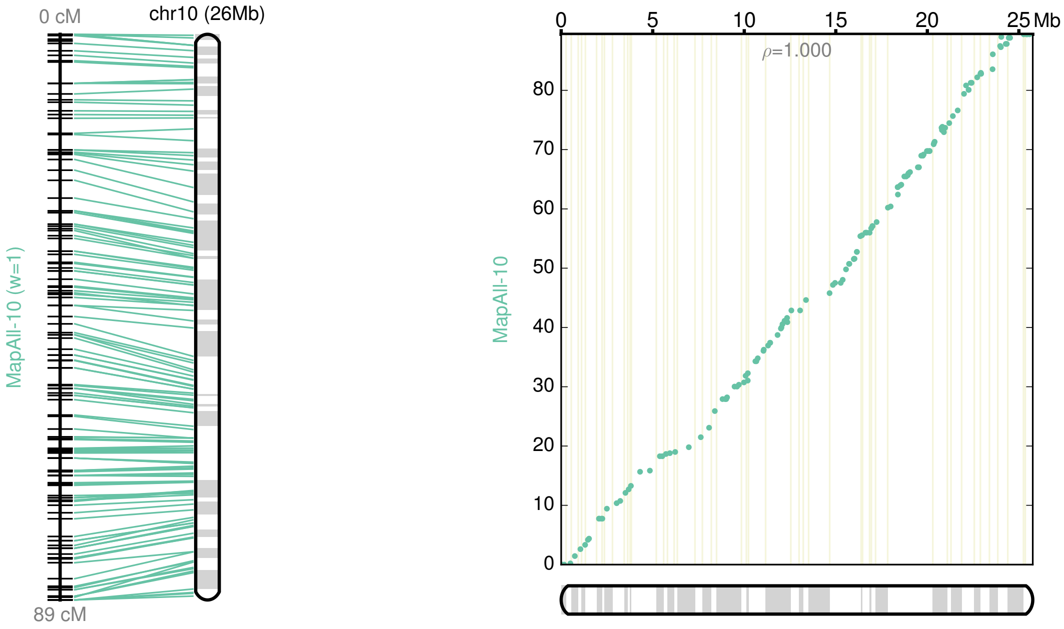

Figure S11. Anchored chromosome 11.

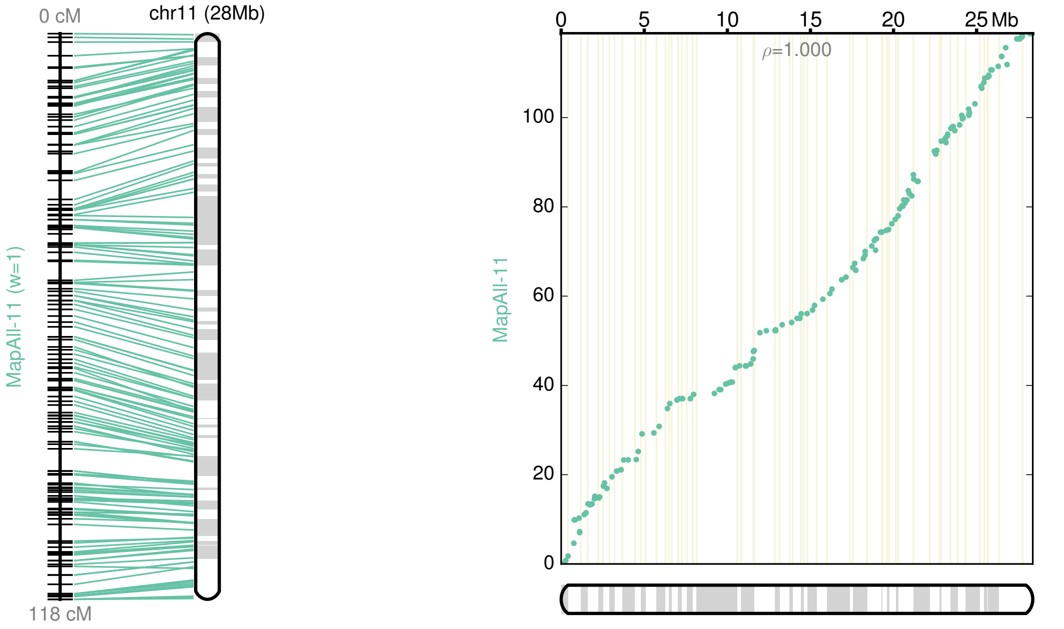

Figure S12. Anchored chromosome 12.

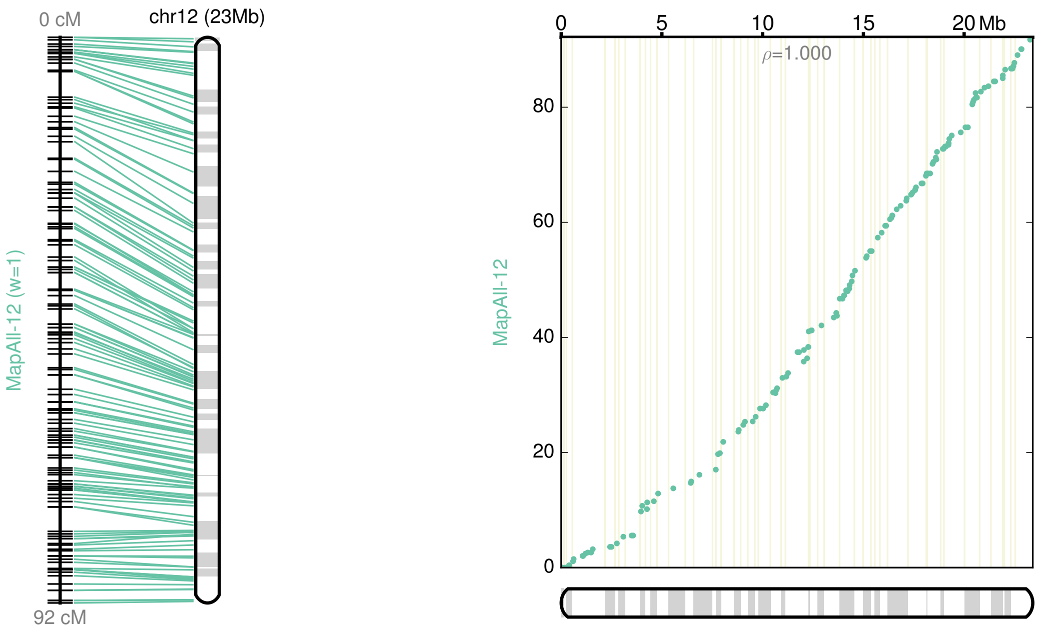

### 2. Annotation

#### 2.1. Repeat masking

The REPET v2.2 package (20) was used for *de novo* identification of repetitious sequences in both assembled *P. americana* genomes. REPET screens for structural features characteristic of transposable elements (TEs), searches for similarity with known TE sequences from RepBase (21), and probes for virtually all Pfam (22) hidden Markov models. Prior to gene prediction, the avocado genomes were masked using the RepeatMasker program (http://www.repeatmasker.org) and the *de novo* predicted TEs identified in each genome. Bases masked using default parameters represent around 50% of avocado genome sequence.

#### 2.2. Identification of protein-coding genes

The *ab initio* and evidence-directed predictor AUGUSTUS (23) was specifically trained for *P. americana* species using transcriptomics data and the available AUGUSTUS training web tool (24) (http://bioinf.uni-greifswald.de/webaugustus/). The transcriptome dataset was generated from two libraries (one from the Hass cultivar and one from the *drymifolia* variety). These libraries were prepared using the TruSeq RNA sample preparation kit and were sequenced on an Illumina HiSeq2000 sequencer. For the library preparation, RNA was isolated from several samples, including different organs in different development stages; these were pooled together in order to generate a unique library from each individual. Each library was sequenced in a single lane of a flow cell (unigenes can be found in Supplementary data 1). Additionally, a broad range of transcriptome sequences previously reported (25, 26) were also included in a new assembly process. Prior to assembly, the sequenced libraries for each individual were processed by using CASAVA version 1.8.2 to produce 100-bp paired-end sequence data in fastq format. To ensure the high quality of the reads, the fastq files were processed using a python script (https://github.com/Czh3/NGSTools/blob/master/qualityControl.py) with stringent quality parameters such as: -q 25 (Minimum quality score to keep), -p 95 (Minimum percent of bases that must have at least [-q] quality) and -a 30 (the average quality of the paired-end reads). The high quality R1-R2 read pairs were assembled using the Trinity software package (27). A total of 187,457 and 144,183 read pairs were generated from the Hass cultivar and *drymifolia* variety, respectively (see additional data file). These unigenes were used to train AUGUSTUS and estimate parameters to predict gene models in both sequenced genomes. In addition, the Maker-P pipeline (28) was used to improve the gene models predicted by AUGUSTUS. Inputs for Maker-P included the *de novo* draft genomes, the corresponding assembled transcriptomes, the species-specific repeat libraries predicted using REPET, and protein databases containing annotated proteins for *Amborella trichopoda* (a basal angiosperm), *Setaria italica* (foxtail millet, a monocot) and *Aquilegia coerulea* (columbine, a basal eudicot). Versions of these proteomes (the same as used for OrthoMCL; see section 3.1, below) were downloaded from the CoGe OrganismView database (http://genomevolution.org/CoGe/OrganismView.pl).

Based on this pipeline, a total of 26,954 gene models were identified in the *drymifolia* variety genome, while for the Hass cultivar, the gene number predicted was 33,378. The Maker-P pipeline, similarly to the majority of gene model predictors, has a tendency to overpredict genes, some of them considered as *ab intio* gene models due to lack of homology and/or transcriptional evidence. Considering the deep sequencing of avocado transcriptomes and the great number of reference proteins used, *ab initio* gene models were not included in future analyses. In this filtering, a similar number of evidence-based protein coding genes was identified in each genome, 22,441 from the *drymifolia* variety and 24,616 from the Hass cultivar. In both cases, around 70% of the proteins derived from the predicted gene models represent at least 70% of the homologous proteins identified in at least three of the five species selected to carry out the annotation process by the top-BLAST-hits method (see details in section 2.3).

#### 2.3. Homology search and functional annotation of *P. americana* protein-coding genes

The *P. americana* proteins predicted from each genome were BLASTed against the *Amborella trichopoda* proteome (AmTr v1.0; 26,811)(29), alongside proteomes from four other angiosperm species for which high quality of the predicted gene models is known. These highly curated genomes were selected in order to avoid the use of inaccurate gene models, which are common and difficult to avoid when a manual curation process has not been undertaken. The species used in these BLAST comparison were *Sorghum bicolor* (33,032 proteins)(30), *Vitis vinifera* (26,343 proteins)(31), *Solanum lycopersicum* (34,727 proteins)(32) and *Arabidopsis thaliana* (TAIR10, 27,416 proteins). Only ~5% of predicted *P. americana* proteins found no match in any of the five species that were selected for BLAST-based annotation (Supplementary Tables 11 and 12). Predicted gene models were considered as complete when the avocado protein derived from their corresponding gene model represents at least 70% of the length of three of five proteins identified as homologs

(Tables S14 and S15). Overall, these data suggest that the annotation is of comparable quality to the grape (31), tomato (32), coffee (33), cacao (34, 35) and other recently published plant genomes.

Functional domains in *P. americana* genes were identified by comparing their translated proteins against the Pfam database (22). Gene Ontology (GO) terms for each gene were obtained from the corresponding *Arabidopsis* homologs. Additionally, the avocado genes were also analyzed using the KEGG Automatic Annotation Server (KAAS; http://www.genome.jp/tools/kaas/) to provide annotations of KEGG Orthology (KO) codes. The bi-directional best hit (BBH) method was used. Enzyme Commission (EC) numbers were also assigned based on the annotations extracted from Kyoto Encyclopedia of Genes and Genomes (KEGG) (Tables S8 and S9).

Table S8: (See additional file.) Annotation of predicted gene models from *Persea americana* var. *drymifolia*.

Table S9: (See additional file.) Annotation of predicted gene models from *Persea americana* cv. Hass.

### 3. Comparative genomic analyses

#### 3.1. Identification of clusters of orthologous genes using OrthoMCL

The OrthoMCL program was used in order to compare the gene content of each *P. americana* genome against those contained in other genomes selected as representatives from specific clades of angiosperms. The following species were selected - the basal angiosperm *Amborella trichopoda* (CoGe genome version ID 19514); Monocots: *Musa acuminata* (banana; ID 11210), *Zea mays* (maize; ID 16904), *Sorghum bicolor* (sorghum; ID 24740), *Setaria italica* (foxtail millet; ID 23469), *Brachypodium distachyon* (ID 25040), *Oryza sativa* (rice; ID 8163) and *Spirodela polyrhiza* (greater duckweed; ID 24105); Eudicots: *Aquilegia coerulea* (columbine; ID 10706), *Coffea canephora* (coffee; ID 19443), *Solanum lycopersicum* (tomato; ID 12289), *Utricularia gibba* (bladderwort; ID 19475), *Vitis vinifera* (grape; ID 19990), *Cajanus cajan* (pigeon pea; ID 12470), *Prunus persica* (peach; ID 8400), *Populus trichocarpa* (Poplar; cotton wood; ID 8154), *Theobroma cacao* (chocolate; ID 10997) and *Arabidopsis thaliana* (from arabidopsis.org). First, TEs were screened and filtered out by BLASTP search (e-value $\leq 1\times10^{-6}$, bit score $\geq 50$) against RepBase (21). Splicing isoforms were also removed and only representative gene models were considered. Finally, an all-against-all comparison using BLASTP was performed with an e-value cut-off of $1\times10^{-10}$. Clustering was then performed based on a Markov cluster (MCL) algorithm using OrthoMCL v1.4(36) with inflation value of 1.5. 402,903 of 597,529 protein sequences (67.42%) were clustered into 44,151 orthologous groups (Table S10).

Table S10: (See additional file.) Changes in orthogroup size identified in selected angiosperm species.

#### 3.2. Mapping of re-sequenced accessions

Re-sequenced accessions described in section 1 were mapped against the Hass reference genome assembly and the previously reported plastid genome sequence (ID KX437771; (37)). The raw fastq reads were clipped of Illumina adapters and trimmed for low quality regions using trimmomatic (v.0.32;(38)) with the following parameters LEADING:20 TRAILING:20 SLIDINGWINDOW:3:15 MINLEN:35. The resulting paired and single end reads were mapped using bwa mem (v.0.7.12; (39)), filtered using samtools (v.1.3.1; (40)) for low quality mapping (-q 20) and for PCR duplicates (rmdup). The estimated breadth of coverage of the nuclear genome ranged between 70% up to 92%, while that of the plastid genome was constant around 30%, except for Velvick and the reference *drymifolia* accession (75%). Breadth and depth of coverage for each accession are contained in Table S11.

Table S11. (See additional file.) Sequencing coverage of the selected accessions.

### 4. Analysis of nuclear SNPs

Given the uneven and low coverage of the resequenced samples, SNPs were called using ANGSD v.0.923(41) with the following parameters for genotype calling and likelihood calculations: -GL 1 -doVcf 1 -doMaf 1 -SNP_pval 1e-6 -doMajorMinor 1 -doPost 1 -doCounts 1 -doGeno 1. The resulting vcf file was lifted over to the anchored chromosomes using picard LiftoverVcf (v2.4.1), and per-site information tags were filled using the plugin fill-tags of bcftools v1.5. Several pruning steps were taken to remove low quality sites: we kept sites with no more than 20% of missing data (--max-missing=0.8) and mean depth values (over all included individuals) in the range of 3 (--min-meanDP) to 21 (--max-meanDP), which corresponds to the average depth plus one standard deviation. Applying these filters, we obtained a total of 6.7e6 SNPs across the 12 chromosomes. Furthermore, we removed low frequency – singletons - and strongly linked variants in each chromosome using plink v1.9 (--maf=0.1; --indep-pairwise 100 10 0.4). With these stringent pruning steps, we generated a set of 179,029 SNPs (Table S13; Supplementary data 2).

Table S12. Number of filtered positions considered for independent phylogenetic reconstructions of each chromosome.

| | # sites | | # sites |
|---|---|---|---|
| chr1 | 24,839 | chr7 | 12,969 |
| chr2 | 23,310 | chr8 | 12,948 |
| chr3 | 21,728 | chr9 | 11,284 |
| chr4 | 9,181 | chr10 | 11,024 |
| chr5 | 17,560 | chr11 | 12,445 |
| chr6 | 11,387 | chr12 | 10,354 |

Despite a priori understanding that significant admixture likely occurred during avocado domestication, we investigated the fit of our SNP data to a bifurcating tree. Phylogenetic analysis was performed using SNPhylo on the 179K set of SNPs as well as on SNPs representing each individual chromosome (Table S12). High bootstrap support was achieved at a stringent minor allele frequency threshold of 0.4 (that is, 43,119 sites considered from the 179K set, and an average of ~3,600 sites per chromosome; Figure S13-14), as rare variants caused potential long branch attraction effects and poor topological resolution. The single tree from all SNPs (Supplementary data 2), when rooted with *P.*

*schiedeana*, revealed two main clades: (1) Mexican accessions, with Hass embedded sister to the var. drymifolia reference genome, and (2) Costa Rican/West Indian/Guatemalan accessions, the latter in a derived position sister to the Velvick cultivar. These relationships clearly reflect the admixed origin of Hass and its suspected closest ancestry to Mexican cultivars, and suggest as well that Costa Rican and West Indian accessions are admixed between Guatemalan and other sources. Chromosome-wise phylogenetic trees (Figure S14) support the same relationships across 7 of 12 chromosomes (chromosomes 2, 3, 4, 5, 7, 9, 11), a highly similar topology that groups West Indian and Velvick (chromosomes 8, 10), one topology that embeds Hass accessions within a Costa Rican/West Indian, Guatemalan clade (chromosome 1), and two other topologies within which Mexican and Costa Rican/West Indian, Guatemalan individuals show scrambled relationships (chromosomes 6, 12).

Figure S13. SNPhylo tree based on 43K polymorphic SNPs across the 12 chromosomes.

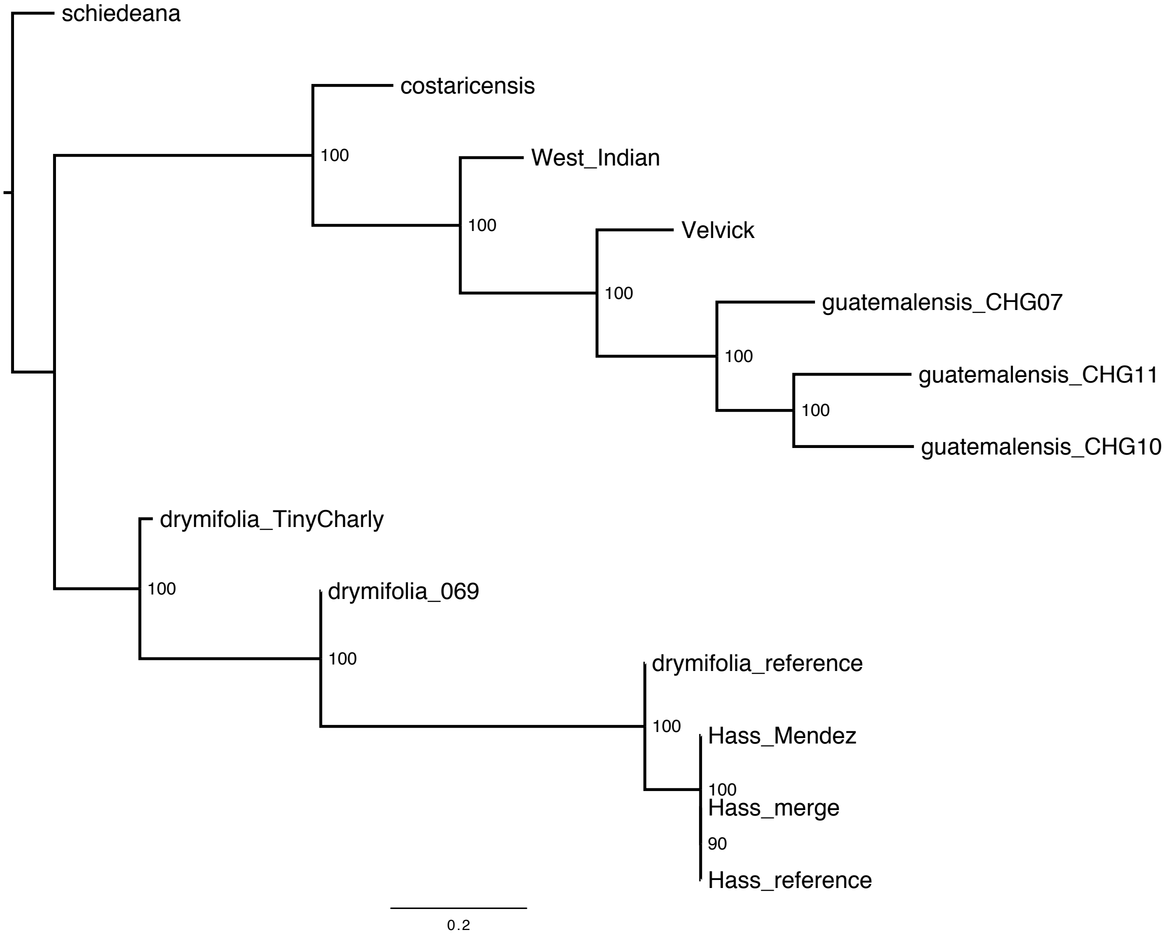

Figure S14. SNPhylo trees for each individual chromosome, 1-12.

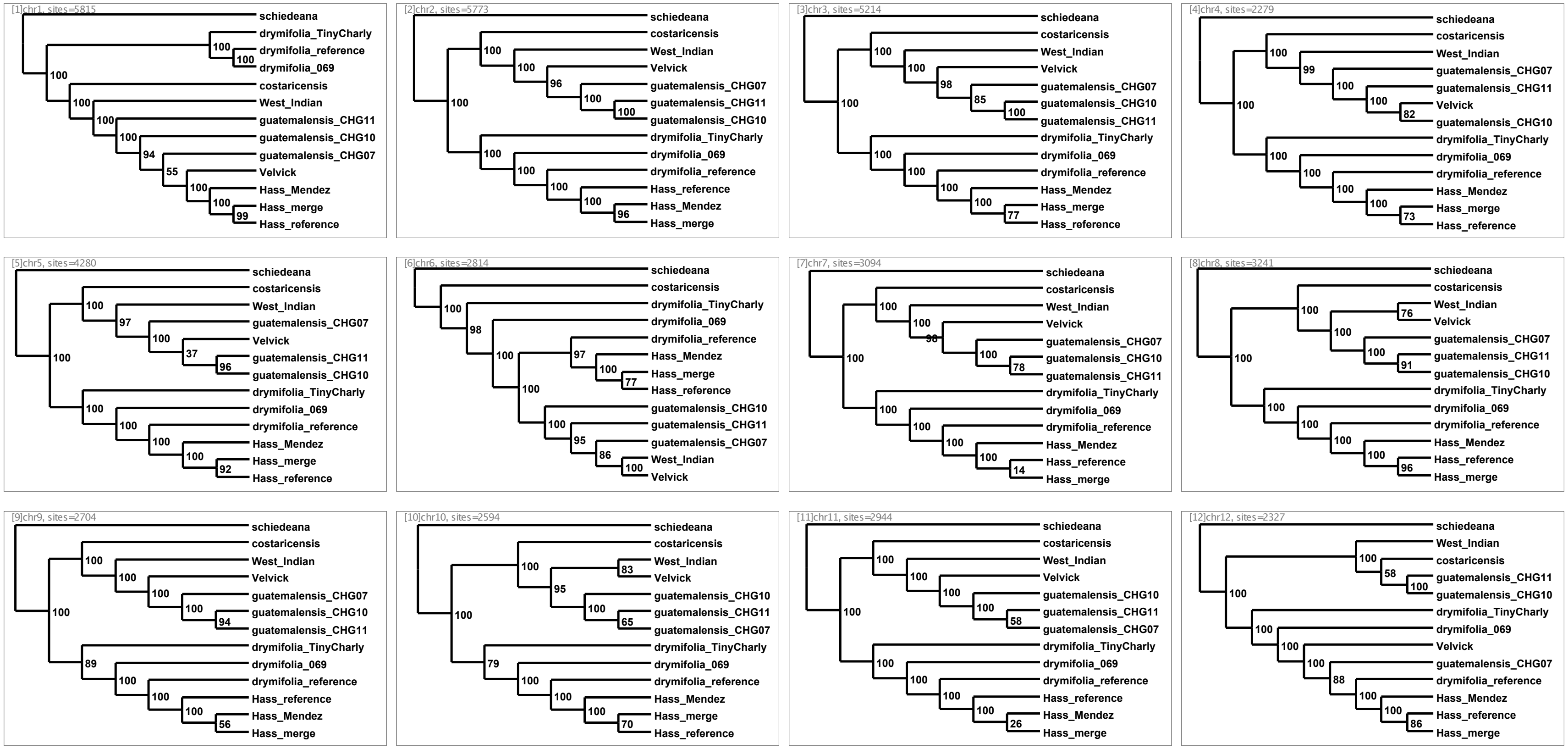

Given that purely cladogenetic behavior of SNP data was not expected among our avocado accessions, we also executed a principal component (PCA) and identity-by-state (IBS) analyses to ordinate the variation in the absence of a tree assumption. We used the functions in SNPRelate (42) for PCA (snpgdsPCA) and IBS (snpgdsIBS) on the 179K SNP set. The PCA resulting from all stringently pruned sites clusters the samples belonging to Hass, Guatemalan and Mexican varieties as expected according to their genetic background, except for Tiny Charly, which behaves as a more divergent Mexican accession (Figure S15). IBS analysis on the same data set not only placed Hass accessions together, but as an intermediate population between Guatemalan and Mexican subpopulations, agreeing with the hybrid nature of this variety. In addition, IBS placed Tiny Charly as an outlier together with *P. schiedeana* (Figure S16), suggesting that our Mexican subpopulation is strongly heterogeneous. Nevertheless, while performing cluster analysis on the matrix of genome-wide IBS pairwise distances (using the functions snpgdsHCluster and snpgdsCutTree in SNPRelate), only one group was determined by permutation score, which could be due to the small number of accessions in our sampling. Based in these observations, we removed Tiny Charly from the Mexican group for the downstream population genomics calculations described in section 5.

Figure S15. Principal component analysis based on the set of pruned 179K SNPs. Top, PCA1 versus PCA2; bottom, a matrix showing PCAs 1-6.

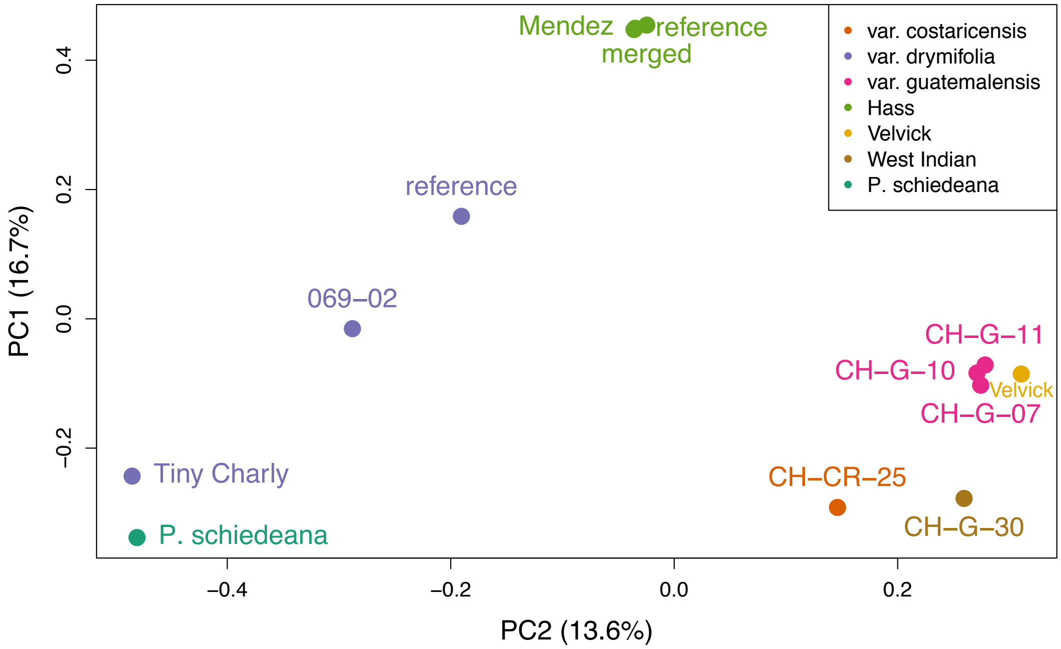

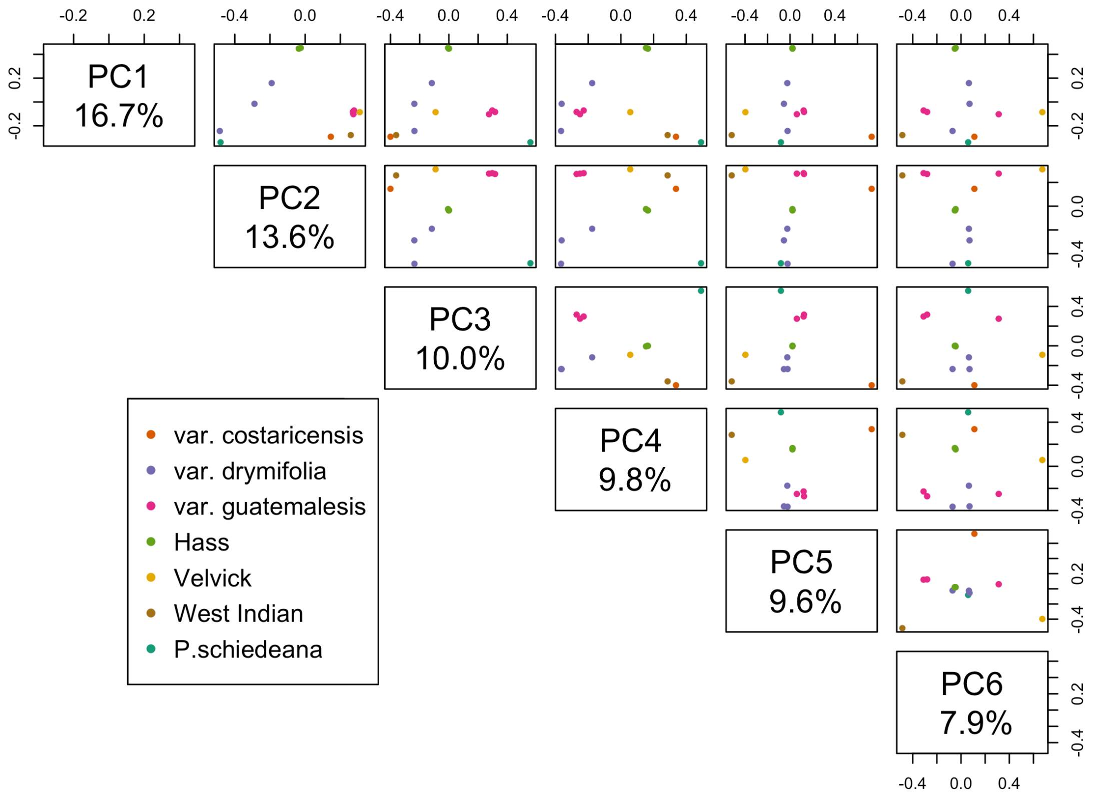

Figure S16. Identity-by-state analysis of the *Persea* samples based on the 179K SNPs set.

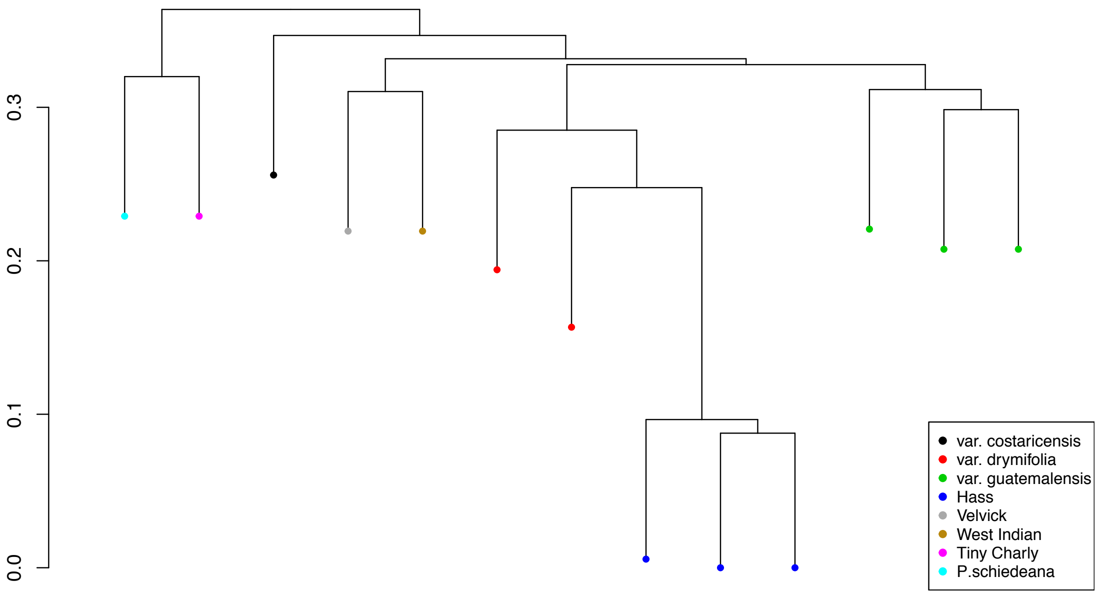

### 5. Population genomics

#### 5.1. Population structure

We evaluated the structure of the avocado varieties in terms of their potential admixtures. First, we ran NGSAdmix on the set of 6.7e6 identified polymorphic sites, with K values ranging from 1 to 6 and a maf threshold of 0.05 (Figure S17). As with the IBS analysis, the Akaike's information criterion selected K = 1 as the preferred number of populations. At K=3 however, two ancestral *P. americana* populations are represented in the Hass genome, as expected from a priori information on the origin of Hass as a hybrid between Mexican and Guatemalan varieties.

Figure S17. Population structure analysis (NGSadmix).

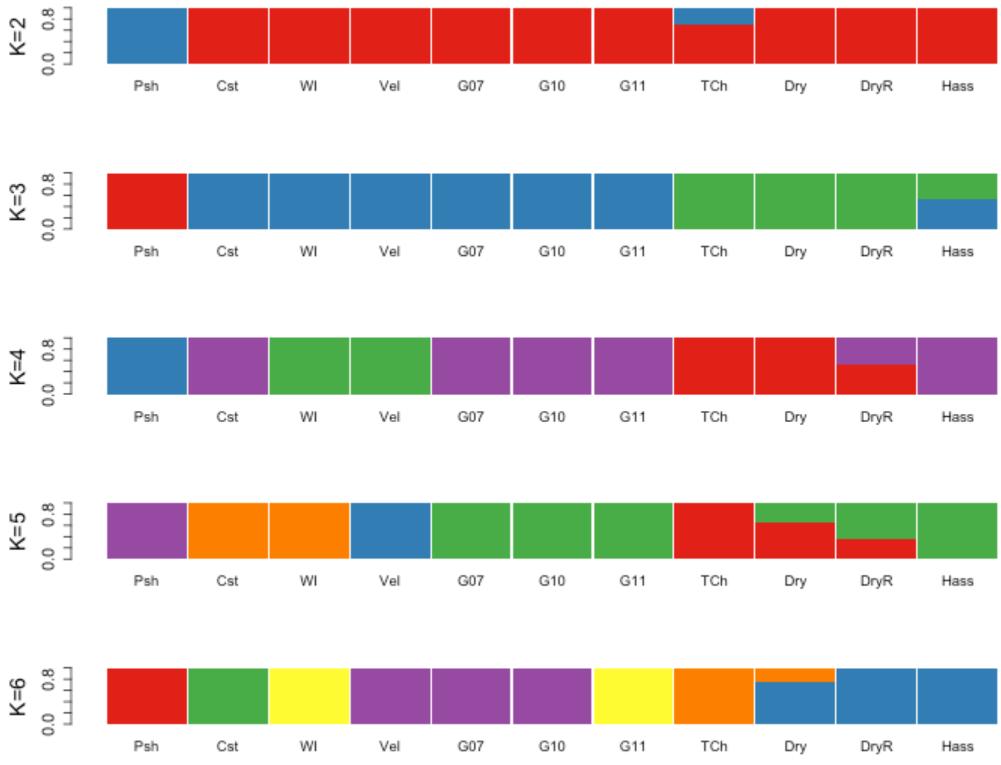

To reduce the effect of linkage disequilibrium, we calculated the Hass admixture proportions using EIGMIX as implemented in SNPRelate (43) on the set of MAF/LD pruned SNPs, leaving Tiny Charly out of the Mexican subpopulation (Figure S18; Table S13).

Figure S18. EIGMIX plots showing the admixture proportions considering three possible ancestral subpopulations from Guatemala (*guatemalensis*), Mexico (*drymifolia*) using *P. schiedeana* as the outgroup.

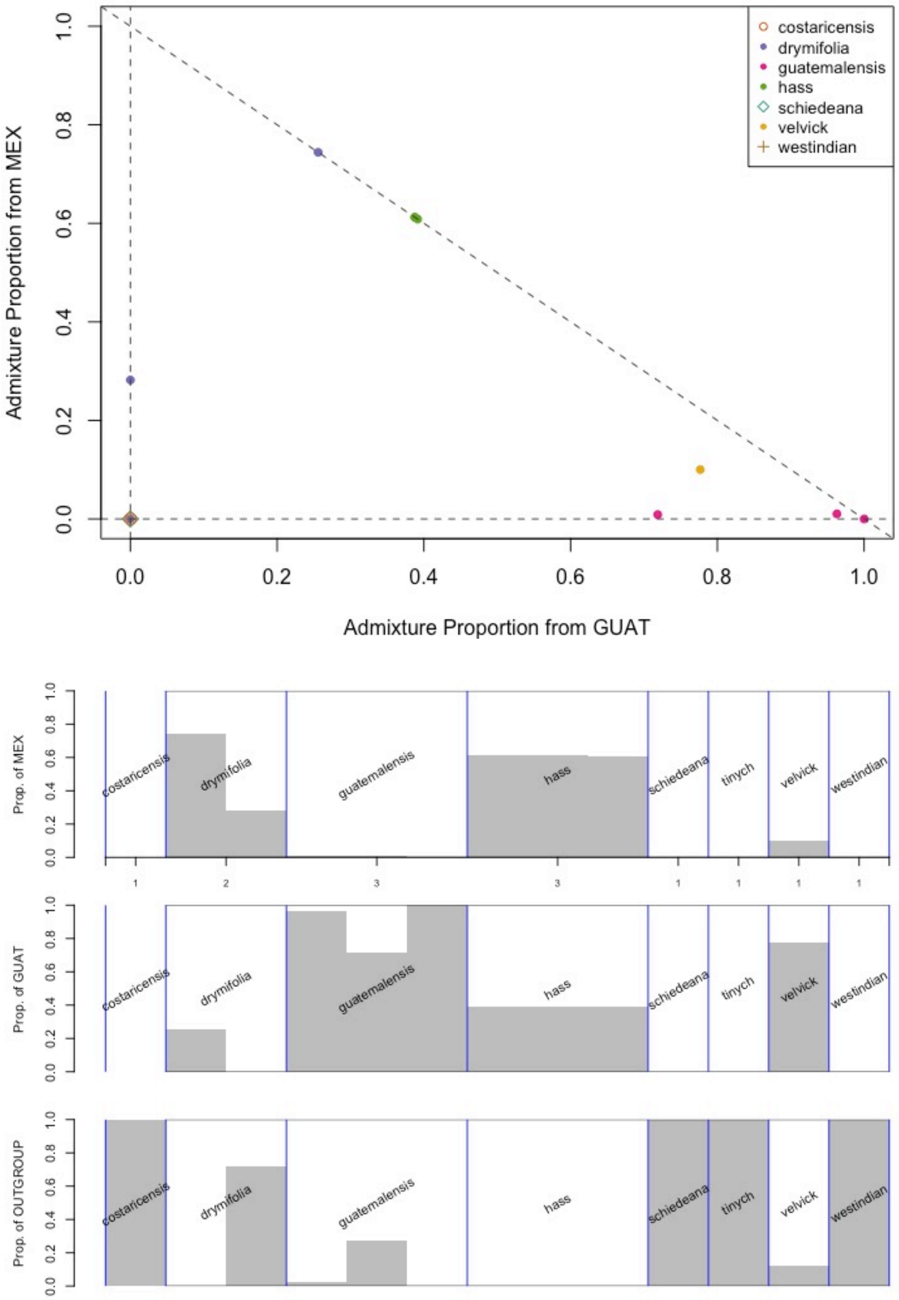

Table S13. Admixture proportions based on the eigen-analysis

**MEX (*drymifolia*)**

| group | num | mean | sd | min | max |
|---|---|---|---|---|---|
| Hass | 3 | 0.61 | 0.00 | 0.61 | 0.61 |
| drymifolia | 2 | 0.51 | 0.33 | 0.28 | 0.74 |
| Velvick | 1 | 0.10 | NA | 0.10 | 0.10 |
| guatemalensis | 3 | 0.01 | 0.01 | 0.00 | 0.01 |
| costaricensis | 1 | 0 | NA | 0 | 0 |
| P. schiedeana | 1 | 0 | NA | 0 | 0 |
| Tiny Charly | 1 | 0 | NA | 0 | 0 |
| West Indian | 1 | 0 | NA | 0 | 0 |

**GUAT (*guatemalensis*)**

| group | num | mean | sd | min | max |
|---|---|---|---|---|---|
| guatemalensis | 3 | 0.89 | 0.15 | 0.72 | 1.00 |
| Velvick | 1 | 0.78 | NA | 0.78 | 0.78 |
| Hass | 3 | 0.39 | 0.00 | 0.39 | 0.39 |
| drymifolia | 2 | 0.13 | 0.18 | 0.00 | 0.26 |
| costaricensis | 1 | 0 | NA | 0 | 0 |
| P. schiedeana | 1 | 0 | NA | 0 | 0 |
| Tiny Charly | 1 | 0 | NA | 0 | 0 |
| West Indian | 1 | 0 | NA | 0 | 0 |

**OUTGROUP (*P. schiedeana*)**

| group | num | mean | sd | min | max |
|---|---|---|---|---|---|
| costaricensis | 1 | 1 | NA | 1 | 1 |
| P. schiedeana | 1 | 1 | NA | 1 | 1 |
| Tiny Charly | 1 | 1 | NA | 1 | 1 |
| West Indian | 1 | 1 | NA | 1 | 1 |
| drymifolia | 2 | 0.36 | 0.51 | 0.00 | 0.72 |
| Velvick | 1 | 0.12 | NA | 0.12 | 0.12 |
| guatemalensis | 3 | 0.10 | 0.15 | 0.00 | 0.27 |
| Hass | 3 | 0 | 0 | 0 | 0 |

### 5.2. Genomic admixture

We looked for signals of hybridization events between three subpopulations of *P. americana* varieties (*guatemalensis, drymifolia* and Hass). We calculated $\hat{f}_d$, $\hat{f}_{dM}$ and $d_{XY}$ estimators of introgression and divergence according to Martin *et al*. 2015 and Malinsky *et al*. 2015 (44, 45) in non-overlapping 100Kb windows, controlling the directionality of gene-flow from *guatemalensis* (P3) to Hass (P2) and from *drymifolia* (P3) to Hass (P2), setting *P. shiedeana* as the outgroup (Figure S19, Table S14). The scripts for parsing the VCF file, calculating genome-wide allele frequencies and the ABBA-BABA statistics in sliding windows were obtained from https://github.com/simonhmartin/genomics_general.

Figure S19. Genome wide statistics of divergence and introgression between *P. americana* subpopulations.

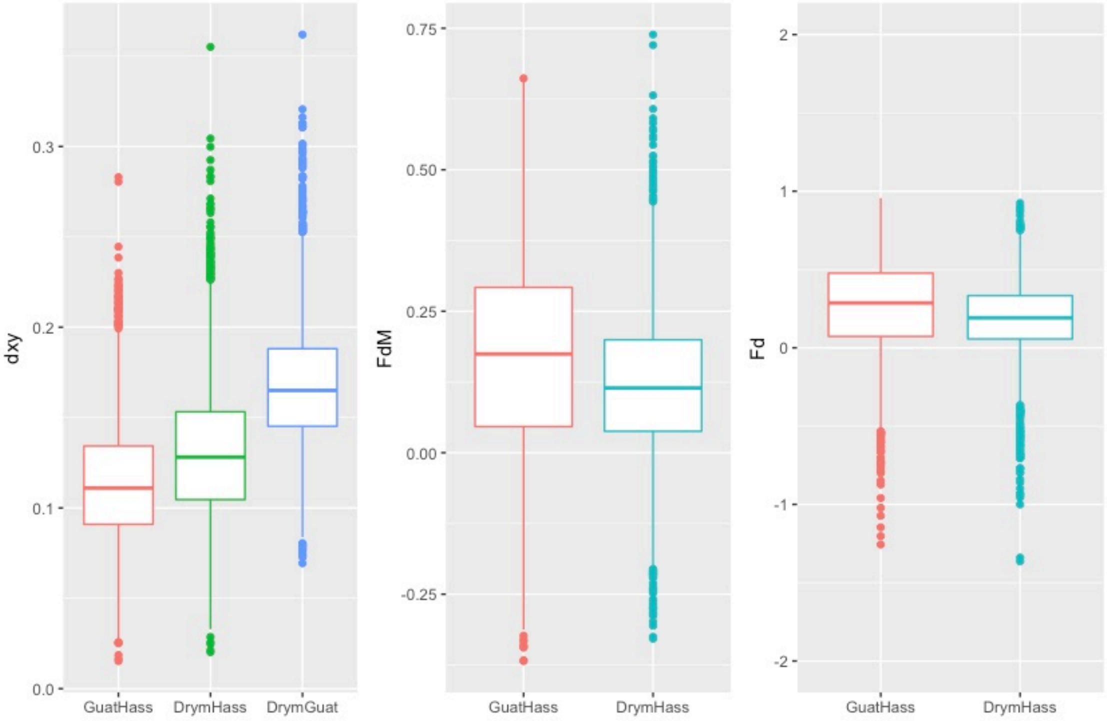

Genomic regions that behave as $\hat{f}_{dM}$ outliers can be distinguished as introgressed from ancestral variation if the absolute genetic distance $d_{XY}$ is also reduced between a donor (P3) and a receptor population (P2), given that in the presence of gene flow, genomic windows coalesce more recently than the species split, so the magnitude of reduction in P2-P3 $d_{XY}$ is greater than in the absence of recombination and hybridization. We evaluated several $\hat{f}_d$ cut-offs (Q50, 75, 90) and observed a remarkable reduction of genetic divergence in the scenario where gene flow occurs from *guatemalensis* into Hass (Figure S20).

Table S14. (See additional file.) Population genomic statistics.

Figure S20. Absolute divergence ($d_{XY}$) between background (green) vs. introgressed (purple) genomic windows at different $\hat{f}_{dM}$ cutoffs, with $d_{XY}$ in Q50, Q75, and Q90 windows.

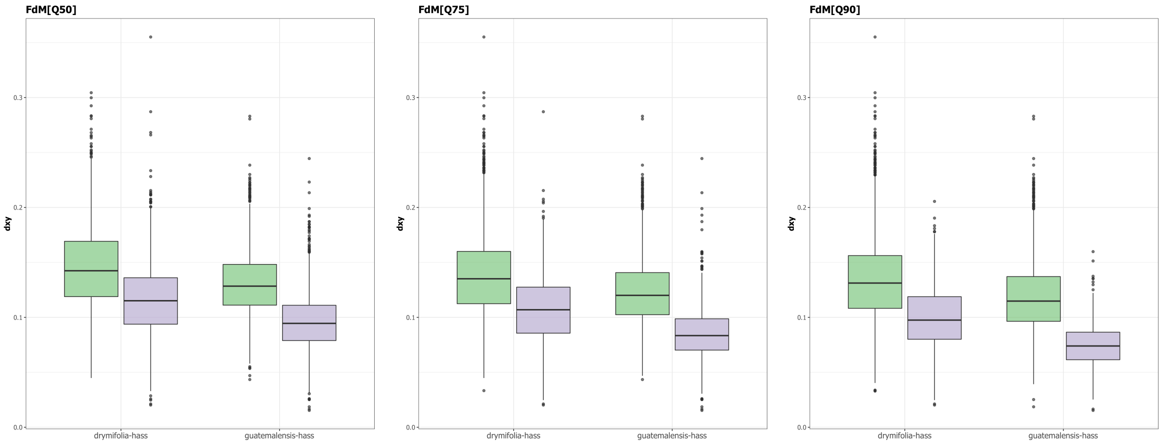

Based on these analyses we were able to define high-confidence regions of *guatemalensis* origin in each chromosome. By keeping those blocks with $\hat{f}_{dM[\text{Guat-Hass}]} > 0.174$ (Q50), $d_{XY[\text{Guat-Hass}]} < 0.113$ and $\hat{f}_{dM[\text{Drym-Hass}]} < 0.114$ (Q50), we obtained 840 windows of *guatemalensis* origin in the chromosomes (Figure S21).

Figure S21. Summary of divergence and introgression estimators in the *guatemalensis*-introgressed genomic windows.

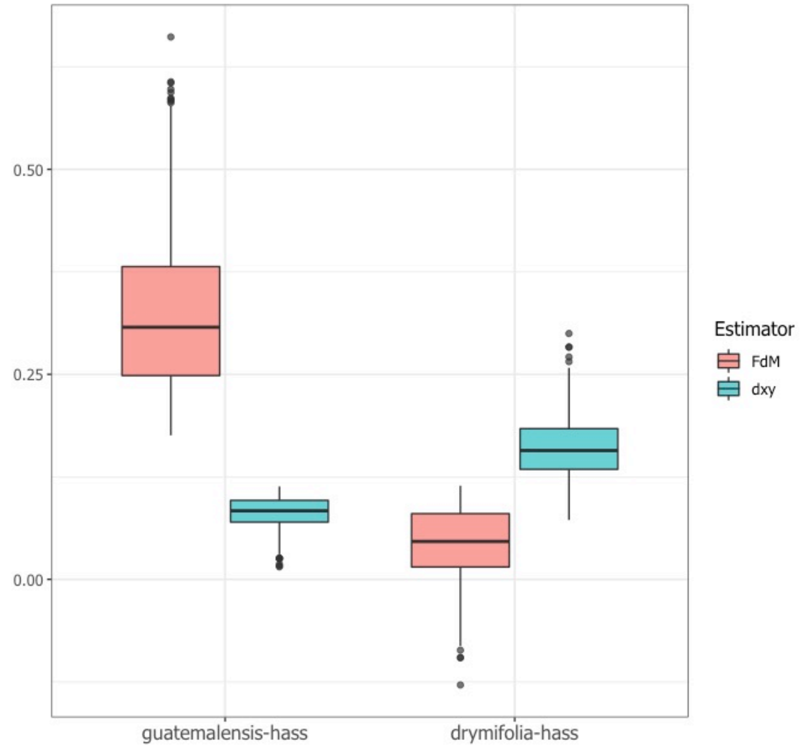

### 5.3. Selective pressures

We looked for signals of artificial selection that might reflect the avocado “domestication” process. We calculated the level of nucleotide diversity (π) in each population, the $F_{ST}$ index to determine regions of high differentiation between varieties, and Tajima’s D (Figures S22-34; Table S14) in order to evaluate any deviations from neutral evolution (https://github.com/simonhmartin/genomics_general/blob/master/popgenWindows.py; vcftools v0.1.13).

Figure S22. Genomic average of nucleotide diversity (π), Tajima’s D and differentiation index ($F_{ST}$) between *Persea* subpopulations.

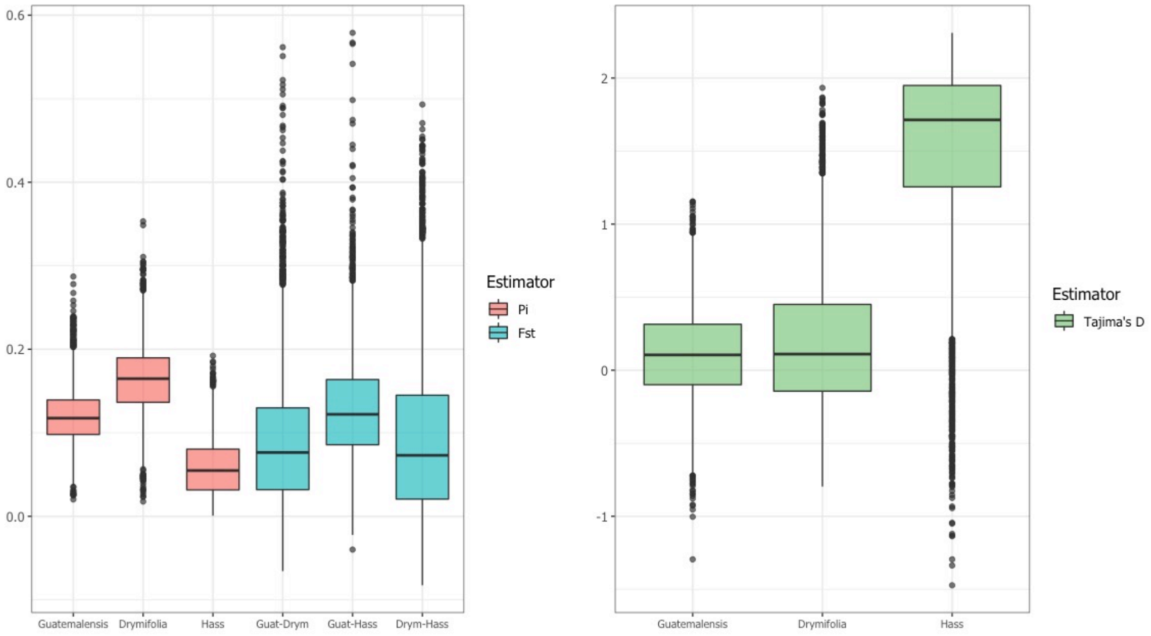

Figure S23. Population genomics statistics along chromosome 1. Each dot in the plots corresponds to statistics for SNP data in non-overlapping 100 Kb windows (level of confidence interval of 0.90 for graphical smoothed conditional means). For $F_{ST}$, $\hat{f}_{dM}$ and $d_{XY}$, the comparison Guatemalensis-Hass is colored in red, while Drymifolia-Hass is colored in blue. In the Tajima's D plot: guatemalensis, red; drymifolia, blue; Hass, green (same for figures S23-34)

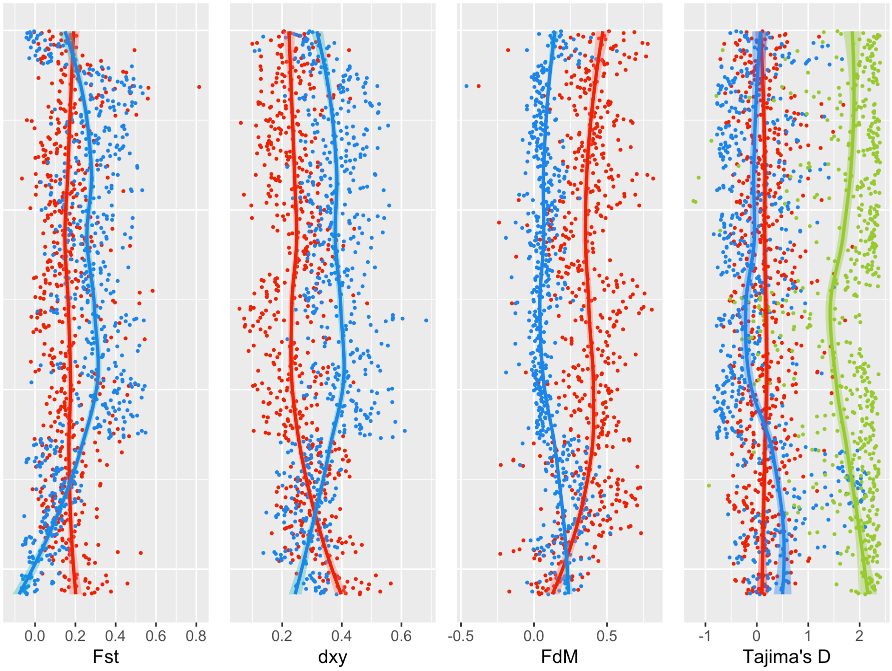

Figure S24. Population genomics statistics along chromosome 2.

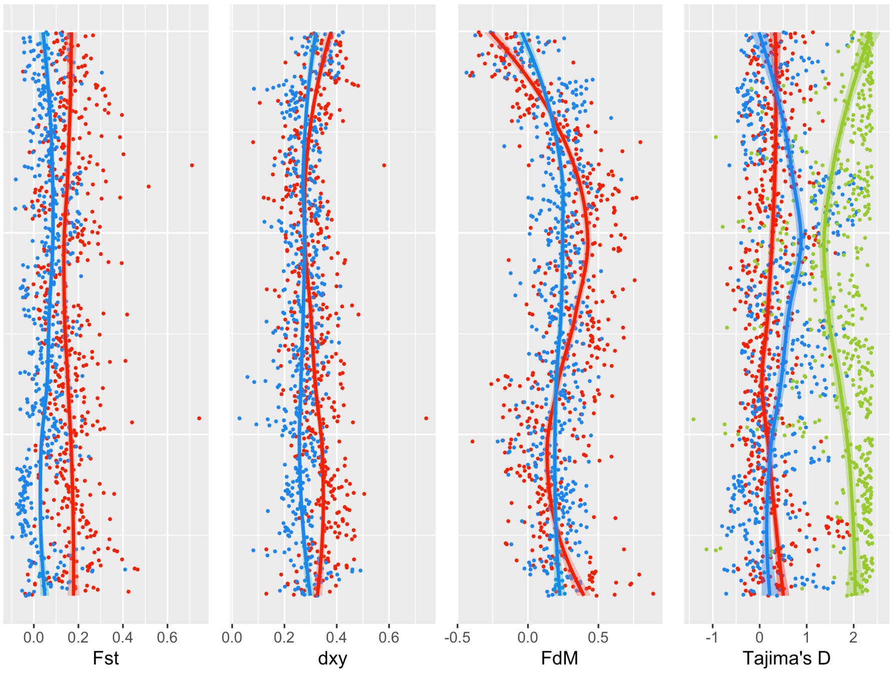

Figure S25. Population genomics statistics along chromosome 3.

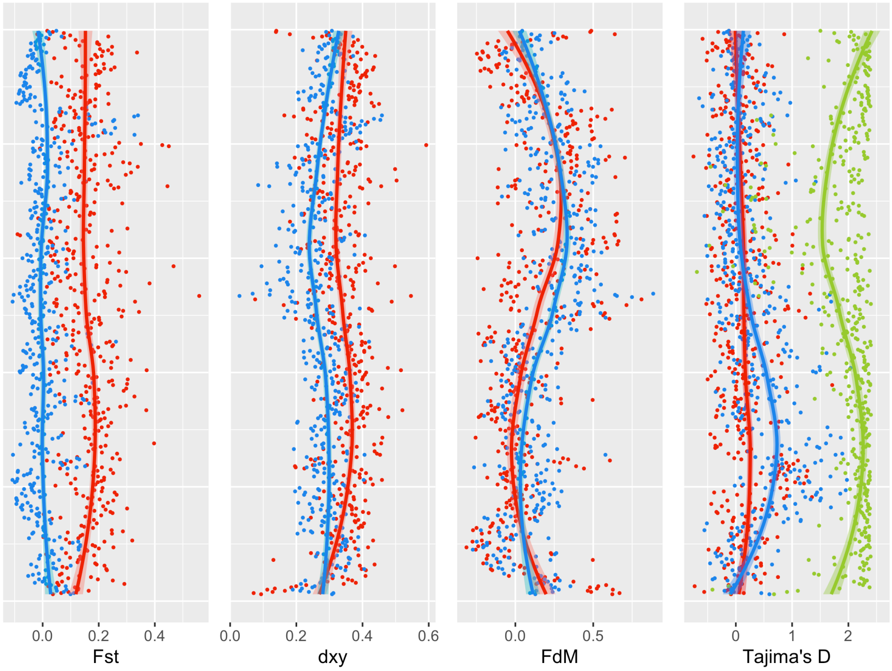

Figure S26. Population genomics statistics along chromosome 4.

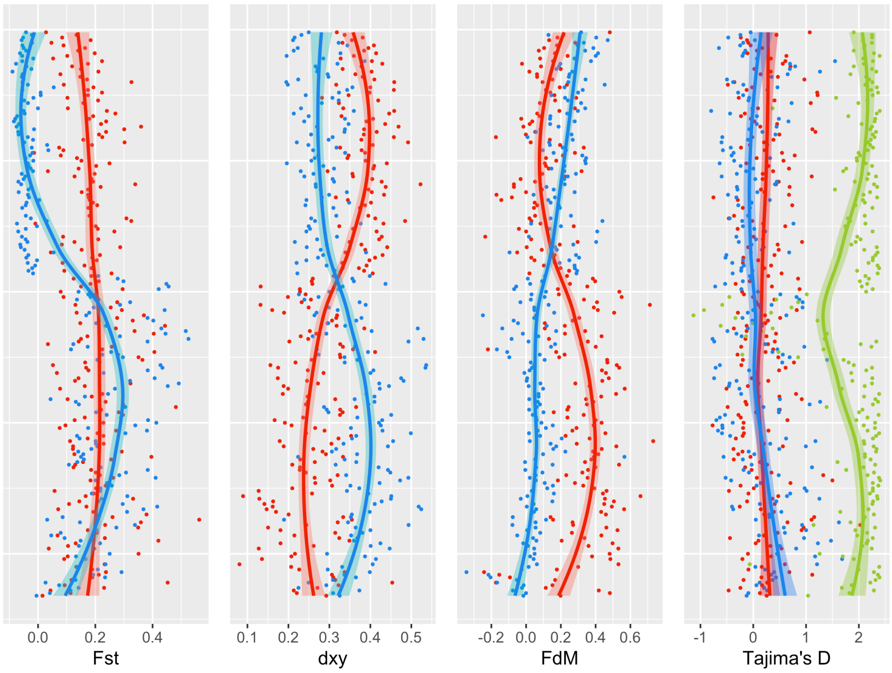

Figure S27. Population genomics statistics along chromosome 5.

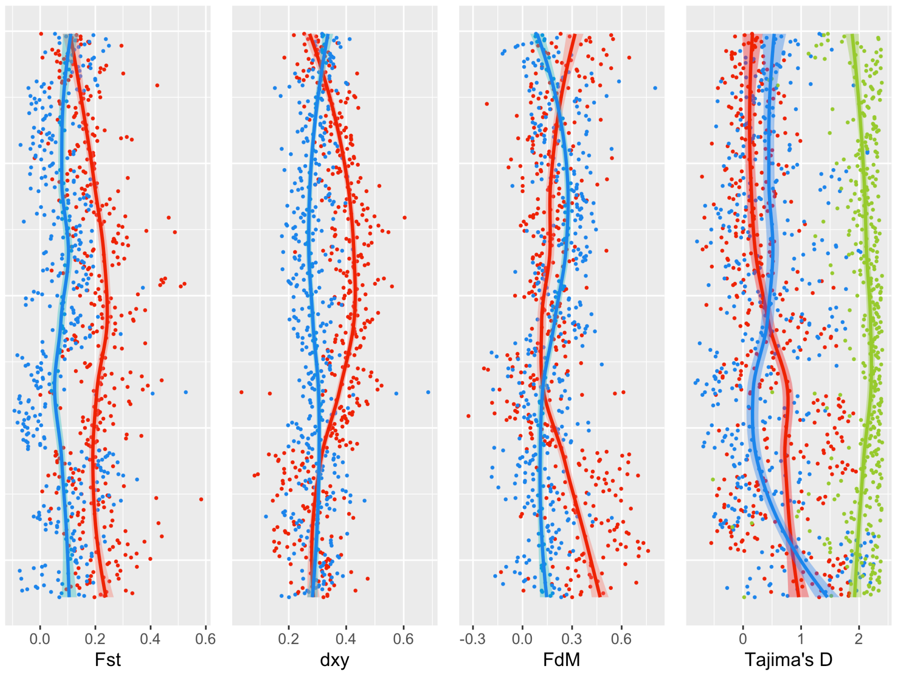

Figure S28. Population genomics statistics along chromosome 6.

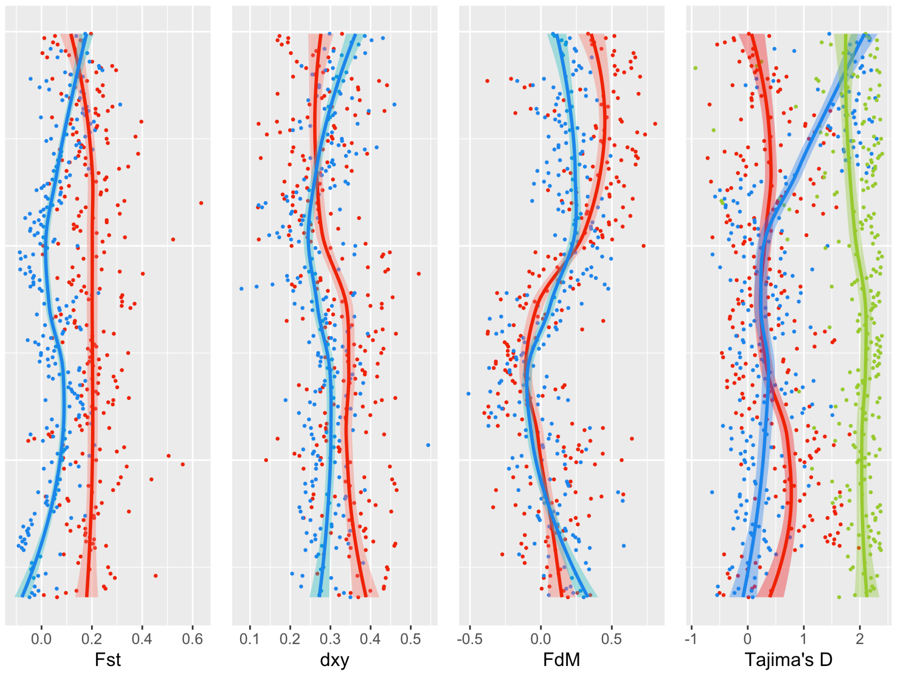

Figure S29. Population genomics statistics along chromosome 7.

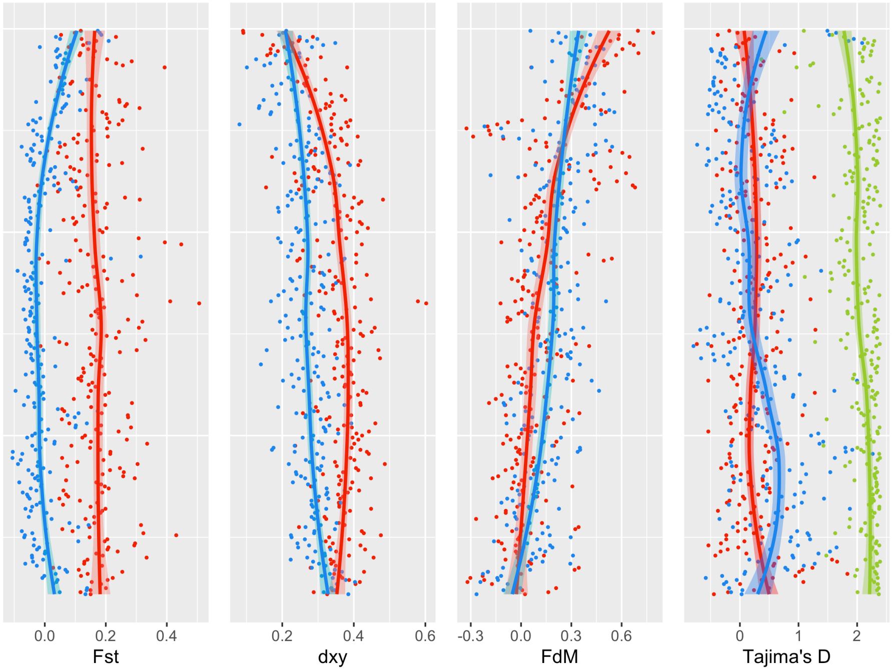

Figure S30. Population genomics statistics along chromosome 8.

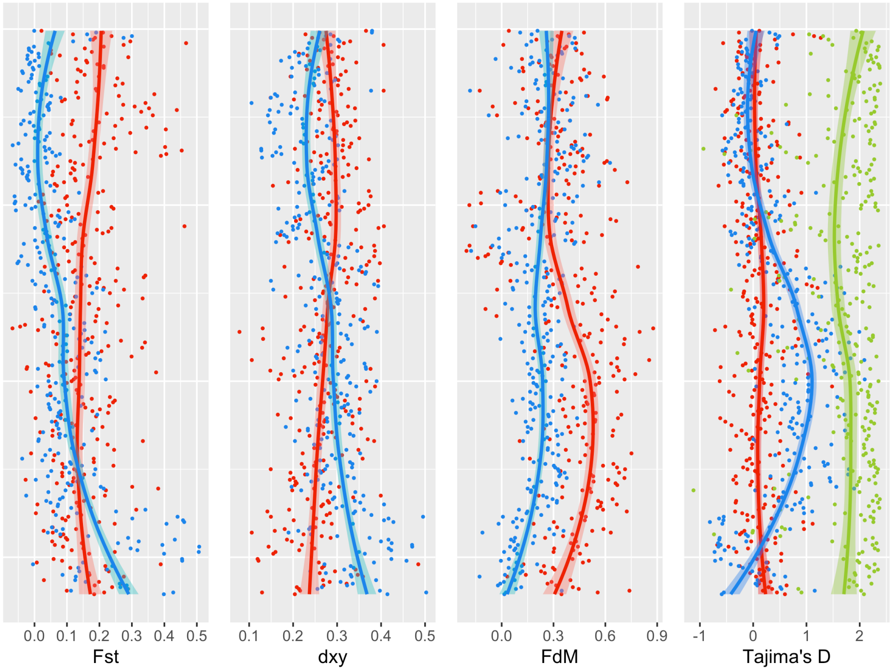

Figure S31. Population genomics statistics along chromosome 9.

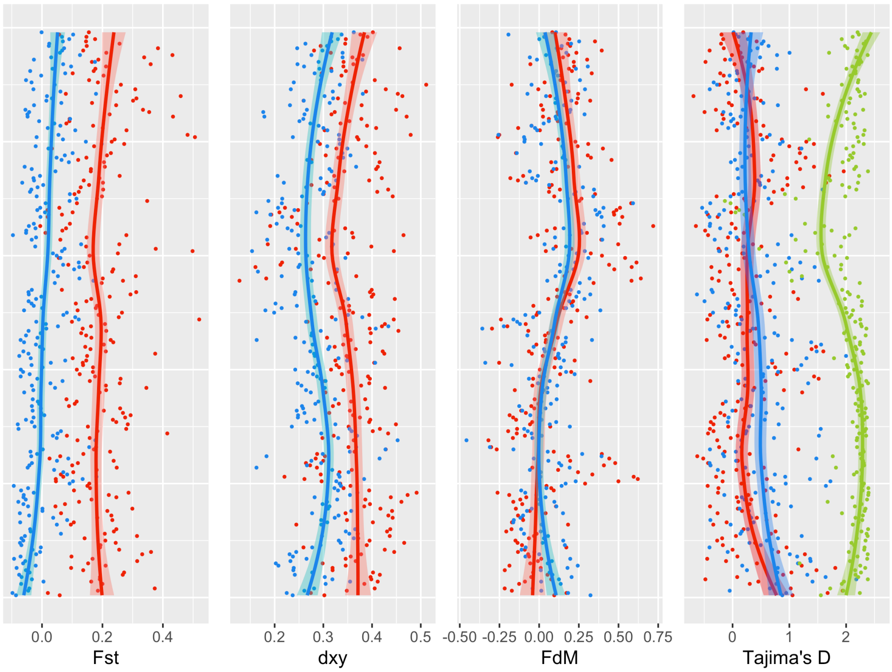

Figure S32. Population genomics statistics along chromosome 10.

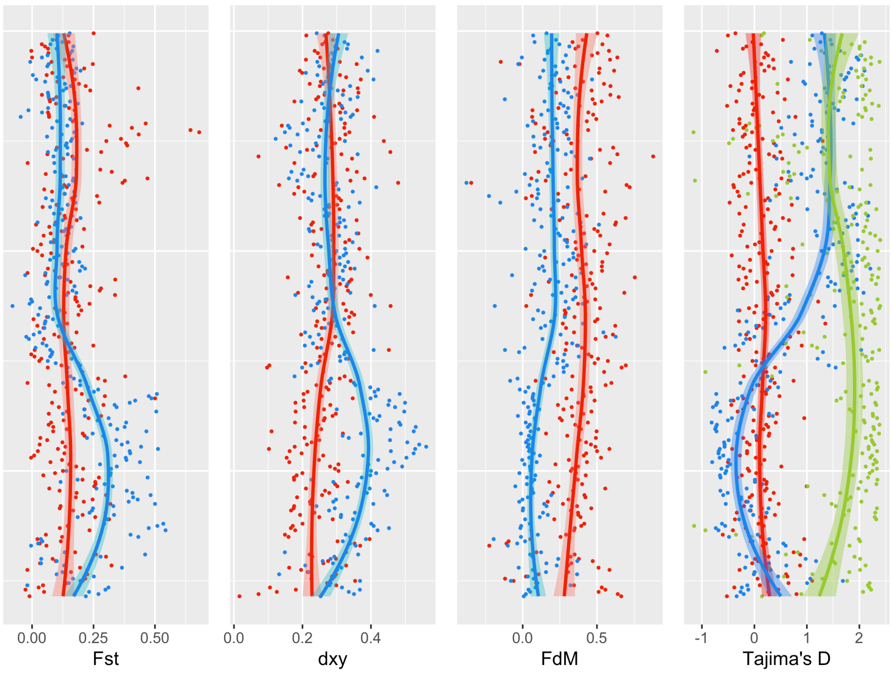

Figure S33. Population genomics statistics along chromosome 11.

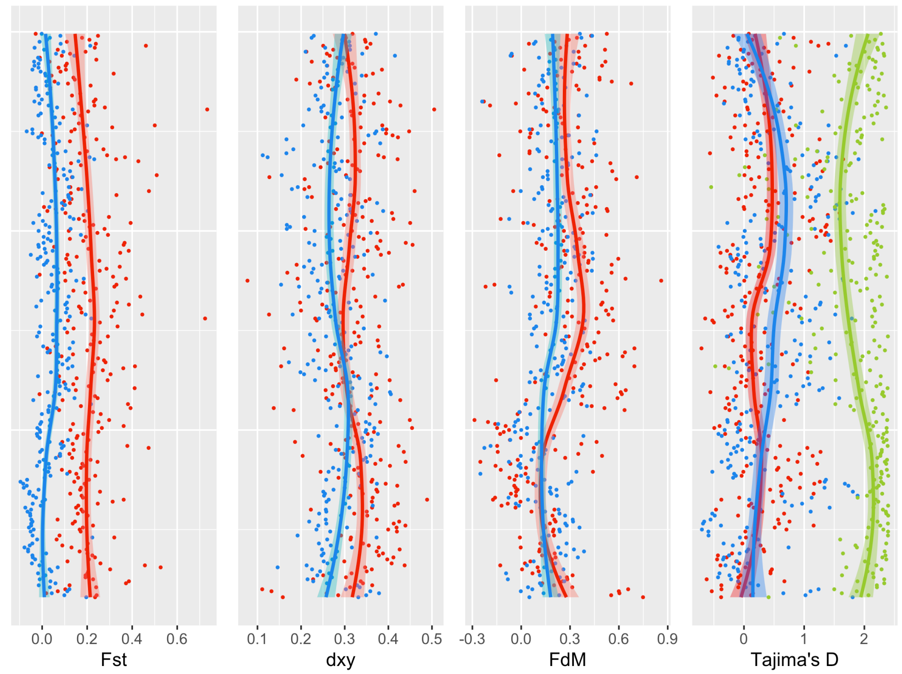

Figure S34. Population genomics statistics along chromosome 12.

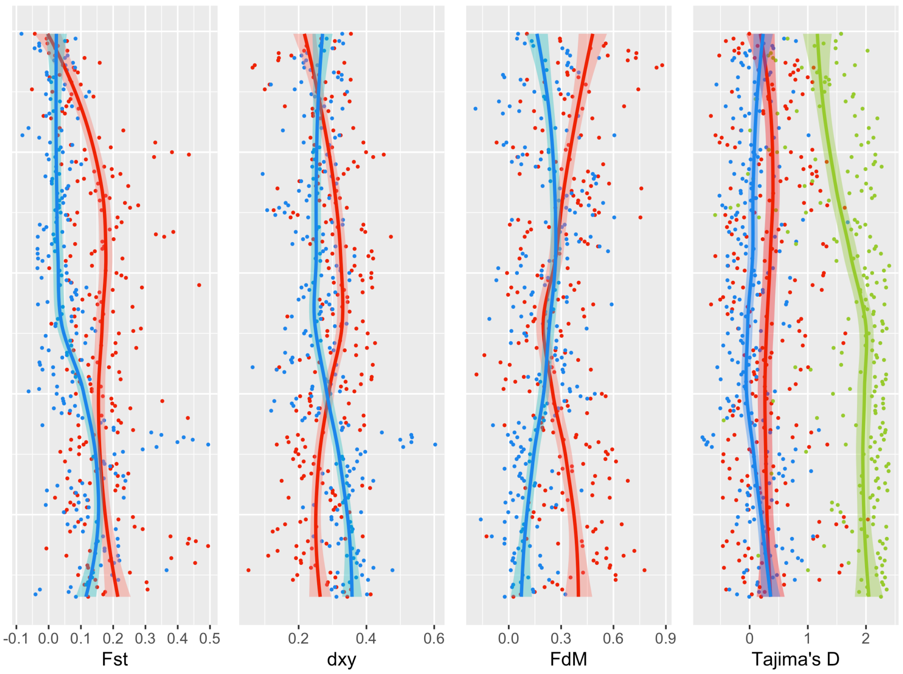

### 6. Whole genome duplication history

#### 6.1. CoGe analyses using the SynMap and GEvo tools

Whole-genome syntenic dotplots based on gene models were constructed using the SynMap tool in CoGe with default parameter settings and determination of synonymous substitution rates (Ks). Hass avocado was compared both with itself and against the *Amborella* and *Vitis* genome assemblies. Internal self:self synteny in Hass avocado revealed two ancient polyploid blocks, and these were mostly resolved in a 4:1 relationship to *Amborella* using the GEvo microsynteny tool in CoGe (main text Fig. 3C; see also Figure S35), suggestive of two whole genome duplications with avocado. MCScan [by Haibao Tang; https://github.com/tanghaibao/jcvi/wiki/MCscan-(Python-version)] was used to generate Fig. 3C. As shown elsewhere (46), *Amborella* is 1:3 relative to *Vitis*, given the presence of the gamma triplication event in the latter species. As such, *Amborella* represents the ploidy level of the angiosperm last common ancestor, and avocado is therefore doubly polyploid relative to it.

Figure S35. Microsynteny view from CoGe GEvo shows that a large 4 Mb block of the *Amborella* genome shows 1:4 synteny with avocado. Gene models hit by HSPs are shown in purple, green otherwise. Top view displays syntenic lines between HSPs, the bottom view without. The analysis can be regenerated at the following link:

https://genomevolution.org/coge//GEvo.pl?prog=blastz;iw=800;fh=15;padding=2;hsp_top=1;colorfeat=1;nt=0;cbc=0;spike_len=15;ca=1;skip_feat_overlap=1;skip_hsp_overlap=1;hs=0;bzW=8;bzK=3000;bzO=400;bzE=30;accn1=evm_27.model.AmTr_v1.0_scaffold00010.298;fid1=368884099;dsid1=80333;dsgid1=19514;chr1=AmTr_v1.0_scaffold00010;dr1up=2000000;dr1down=2000000;ref1=1;mask1=non-cds;accn2=augustus_masked-Ctg0027-processed-gene-5.3-mRNA-1;fid2=936749320;dsid2=99759;dsgid2=29302;chr2=Ctg0027;dr2up=2000000;dr2down=2000000;rev2=1;ref2=0;mask2=non-cds;accn3=augustus_masked-Ctg0035-processed-gene-5.1-mRNA-1;fid3=936750712;dsid3=99759;dsgid3=29302;chr3=Ctg0035;dr3up=2000000;dr3down=2000000;rev3=1;ref3=0;mask3=non-cds;accn4=augustus_masked-Ctg0043-processed-gene-2.0-mRNA-1;fid4=936752074;dsid4=99759;dsgid4=29302;chr4=Ctg0043;dr4up=2000000;dr4down=2000000;ref4=0;mask4=non-cds;accn5=maker-Ctg0098-augustus-gene-7.15-mRNA-1;fid5=936759076;dsid5=99759;dsgid5=29302;chr5=Ctg0098;dr5up=2000000;dr5down=2000000;rev5=1;ref5=0;mask5=non-cds;num_seqs=5;hsp_overlap_limit=0;hsp_size_limit=0

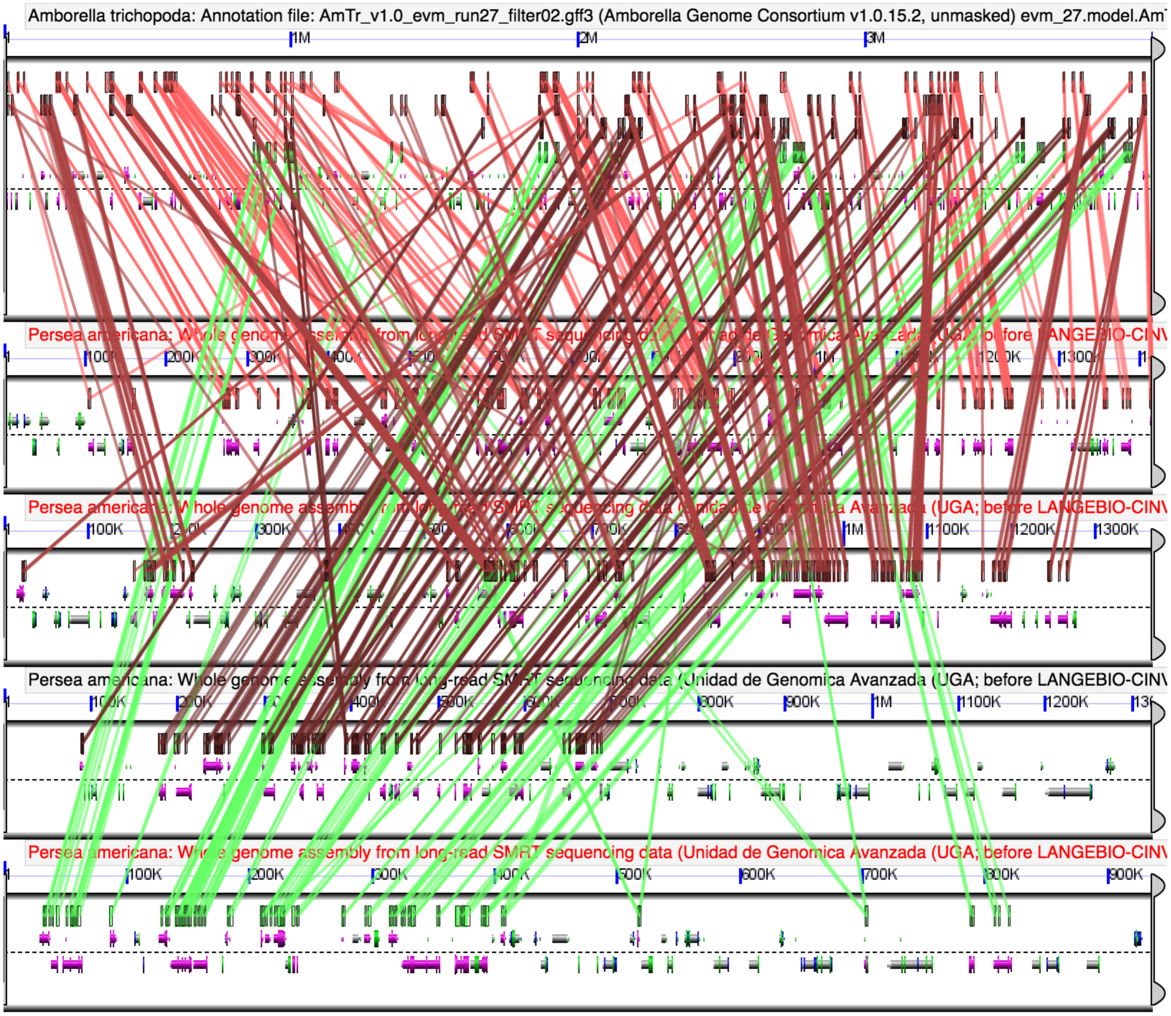
Amborella trichopoda: Annotation file: AmTr_v1.0_evm_run27_filter02.gff3 (Amborella Genome Consortium v1.0.15.2, unmasked) evm_27.model.Am
1M
2M
3M
Persea americana: Whole genome assembly from long-read SMRT sequencing data (Unidad de Genomica Avanzada (UGA; before LANGEBIO-CIN
100K
1100K
1200K
1300K
Persea americana: Whole genome assembly from long-read SMRT sequencing data (Unidad de Genomica Avanzada (UGA; before LANGEBIO-CIN
100K
300K
1M
1100K
1200K
1300K
Persea americana: Whole genome assembly from long-read SMRT sequencing data (Unidad de Genomica Avanzada (UGA; before LANGEBIO-CIN
400K
500K
800K
900K
1M
1100K
1200K
Persea americana: Whole genome assembly from long-read SMRT sequencing data (Unidad de Genomica Avanzada (UGA; before LANGEBIO-CIN
100K
200K
300K
500K
600K
700K
800K
900K

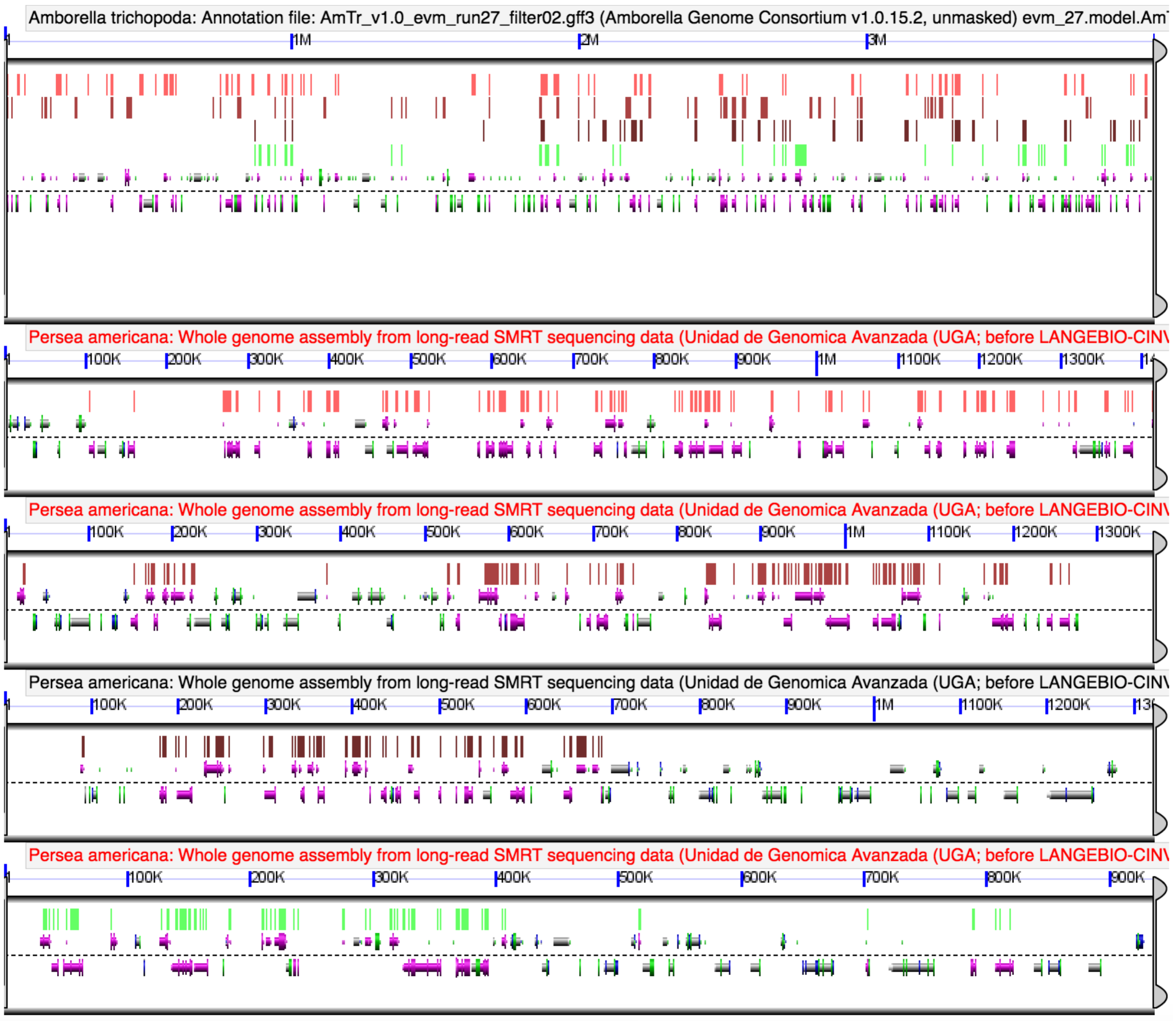

### 6.2. Gaussian mixture modeling of Ks distributions of syntenic homologous genes

Density plotting of Ks values (from Prank CDS alignments; see below) for each orthologous and paralogous gene pair among avocado, *Amborella* and *Vitis* clearly resolved the independence of the *Persea*-specific events from gamma, and therefore any other polyploidy events within core eudicots. Specifically, the two paralogous gene pair peaks in avocado postdate its species splits with *Amborella* and *Vitis* (Figure S36). Pairs of syntenic paralogs within the avocado Hass genome, and of syntenic orthologs between avocado Hass and *drymifolia* avocado, *Vitis vinifera* and *Amborella trichopoda* were extracted from syntenic genomic blocks, defined by genomic regions with a set of at least five collinear genes between the two genomes being compared. Syntenic genomic blocks were identified using the DAGChainer (47) algorithm as implemented in the SynMap (48) tool from the CoGe (49) platform, the Quota Align algorithm and the rest of settings as default. Estimates of $K_s$ were obtained for all pairs of syntenic paralogous and orthologous genes using the CODEML program (50) in the PAML package (v4.8, (51)) on the basis of codon sequence alignments. We employed the GY model with stationary codon frequencies empirically estimated by the F3×4 model. Codon sequences were aligned with PRANK (version 100701) using of the empirical codon model (52) (setting -codon) to align coding DNA, always skipping insertions (-F). Only gene pairs with $K_s$ values in the range of 0.1–5 were considered for further analyses. In order to identify peaks in the frequency distributions of $K_s$ values putatively corresponding to WGD events, we fitted Gaussian mixture models by means of the *densityMclust* function in the R mclust version 5.3 package (53). The Bayesian Information Criterion was used to determine the best fitting model to the data, including the optimal number of Gaussian components (peaks) to a maximum of nine.

Figure S36. $K_s$ plots for syntenic paralogs within Hass avocado, and for syntenic orthologs between Hass, *drymifolia* and other species. Top, density plot showing Hass:Hass syntenic paralogs (cyan), Hass:*drymifolia* syntenic orthologs (green), *Vitis*:Hass syntenic orthologs (purple), and *Amborella*:Hass syntenic orthologs (red). Bottom, histogram depiction of the same data (see also Fig. 3C).

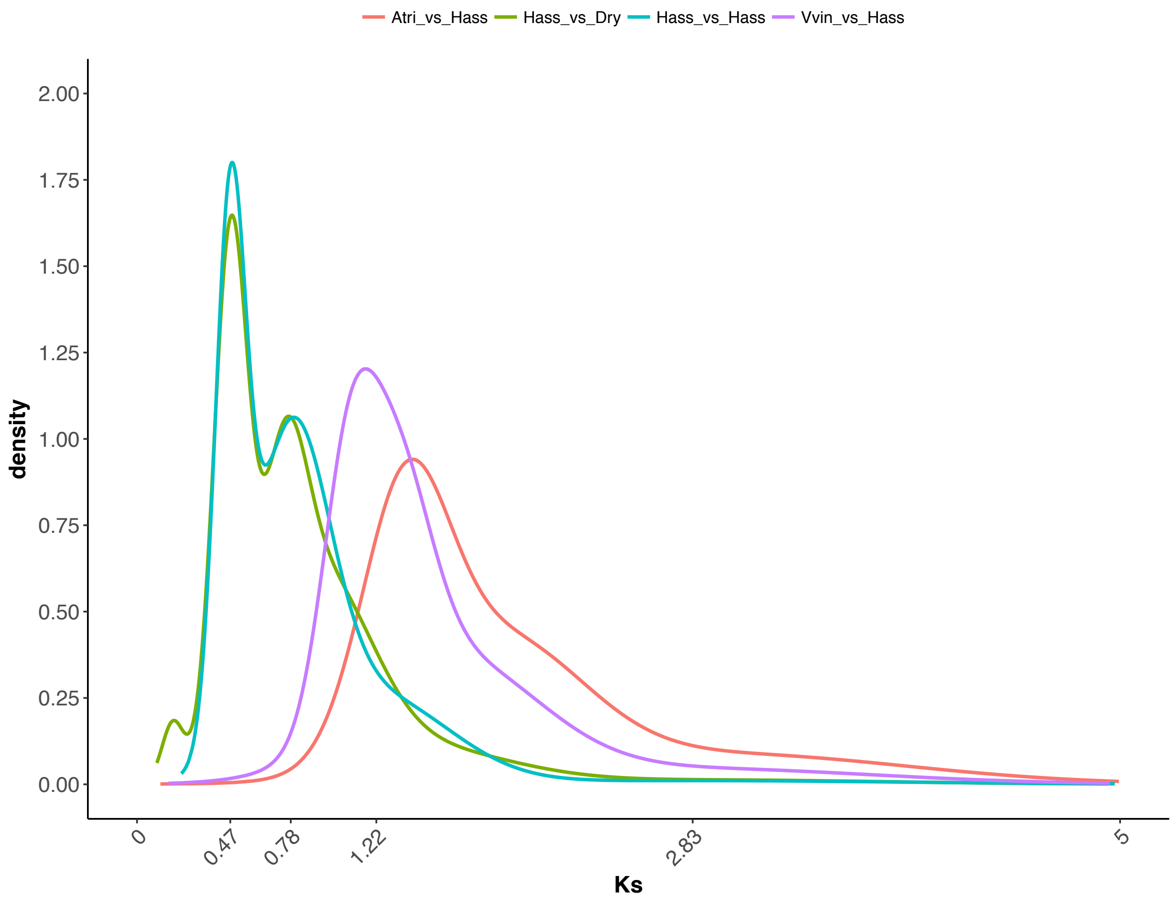

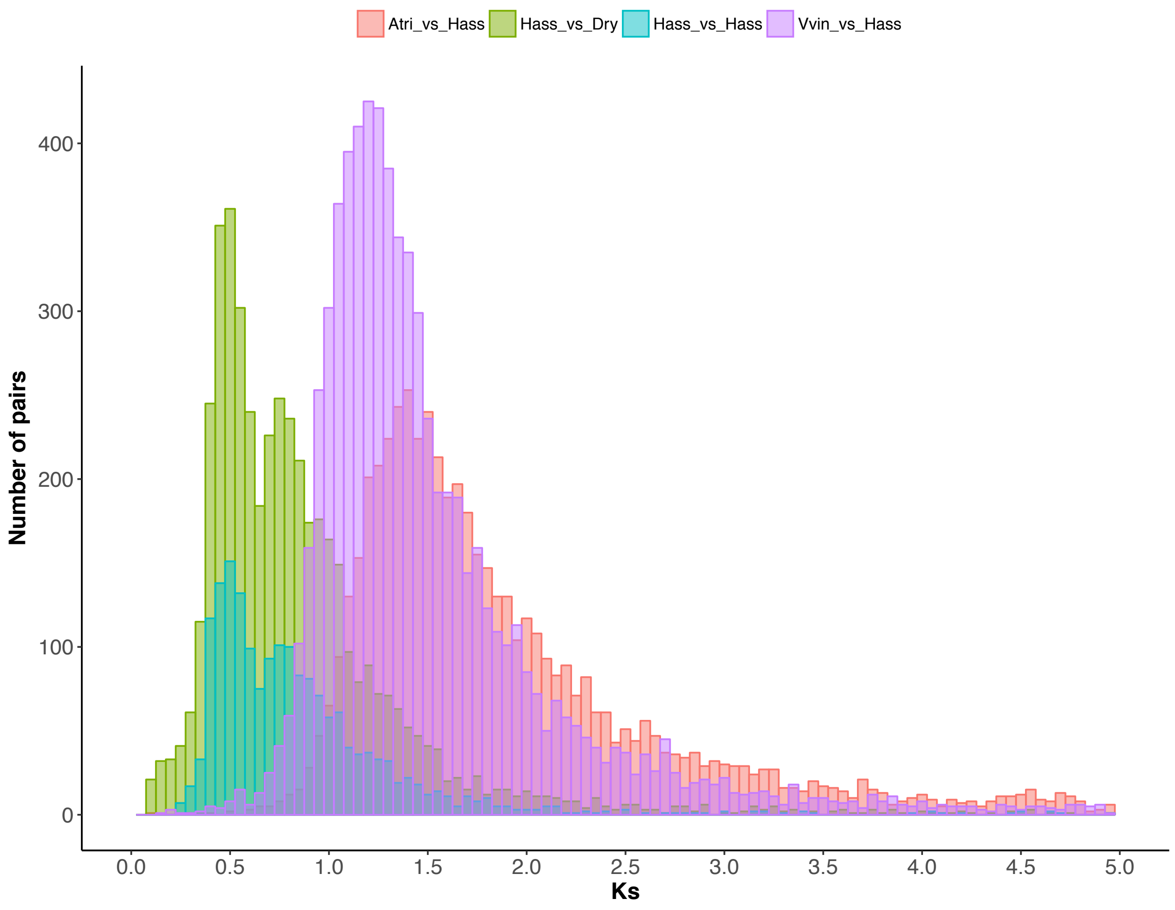

#### 6.3. Verification of avocado's 8:1 genome structural status relative to *Amborella*

We further investigated the polyploid status of avocado using a recently developed quantitative approach that optimizes discovery of orthologous syntenic blocks between species. Specifically, we wished to evaluate whether the most recent polyploidy event in avocado might represent a triplication instead of a simple duplication.

By studying triples of avocado chromosomal regions related through orthologous connections to other genomes, we can distinguish whether these triples are evidence for a recent genome tripling event or a doubling.

We are motivated to use comparative evidence in this way because relying solely on self-comparison to identify the paralogous gene pairs originating in a polyploidization event is subject to particularly high levels of noise. This is partly due to widely shared gene domains, random similarities between genes, transposed elements, other duplications of individual genes unconnected to the polyploidization, expansion of gene families, genome rearrangements and other factors, all of which degrade both paralogy and orthology. More important, however, is the process of fractionation, which operates only between paralogs, and not orthologs, whereby one of the genes in most paralog pairs is deleted. This drastically slashes the number of paralog pairs in the genome. Though it may also reduce the number of ortholog pairs between a polyploid and another genome, this will be quantitatively much less severe a loss.

The problems of validating homologous gene pairs can be attenuated somewhat through recourse to SynMap (49, 54), which retains only those pairs where the two genes are in similar syntenic context (*synteny block*), as defined by a fixed minimum number MinL of pairs of duplicated genes not interspersed with more than a fixed number of genes that are single-copy or have no duplicate within the corresponding block.

Figure S37. Paralogous gene pair similarities for *Persea americana* var. Hass. The total numbers of pairs are for minL=5, 4 and 3: 2038, 2760 and 4217, respectively.

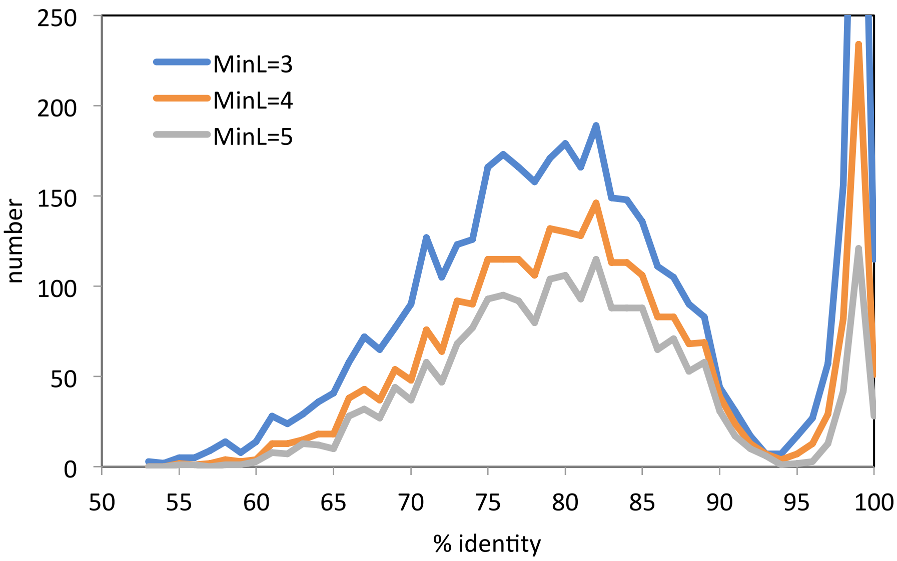

The gene pairs identified by SynMap can be evaluated in terms of sequence identity, as illustrated in Figure S37 for *Persea americana* var. Hass, and in Figure S38 for *Persea americana* var. *drymifolia.* In both cases, we observe a broad region from 75% to 82-84% where the distributions attain a fairly constant maximum sequence identity suggestive of two overlapping distributions with means around 77% and 82%, respectively.

Figure S38. Paralogous gene pairs for *Persea americana* var. *drymifolia*. The total numbers of pairs are for minL=5, 4 and 3: 2641, 3182 and 4221.

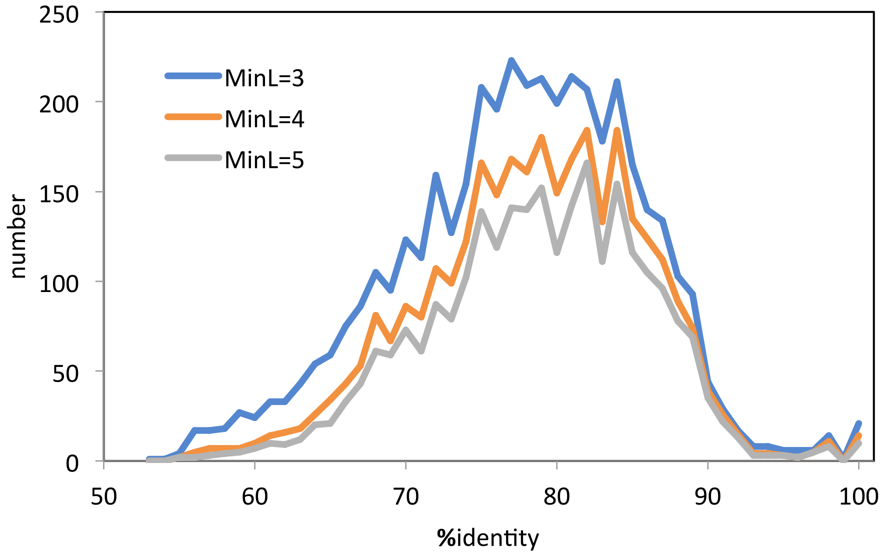

Using minL=4 as a compromise between too few data (minL=5) and data overly contaminated with non-WGD-origin duplicates (minL=3), the number of genes in WGD pairs or larger families surviving fractionation in Hass and *drymifolia* is 3898 (15%) and 5062 (22%) of the total number of genes 25211 and 22917, respectively.

In general, genome self-comparison is much less productive of paralogous pairs than the ortholog pairs resulting from a comparison of a WGD descendant *W* with a related genome *R*. More orthologous genes can be detected in a *W* x *R* comparison than paralogs in a *W* x *W* analysis because fractionation does not eliminate *both* genes of a paralogous pair in *W*. Thus the orthology still shows up with the one remaining paralog in *W* and its homolog in *R*, while the paralogy is completely destroyed by fractionation. Indeed, we may sometimes identify two genomic regions in *W* that were originally duplicates of each other but that retain few or no duplicates between them, simply by discovering that they both contain sufficient numbers of orthologs interleaved in a single region of *R*.

Thus in a sample of 15 diverse angiosperm reference genomes, the number of paralogy pairs in synteny blocks with avocado averages about 8100 (Hass) or 9800 (*drymifolia*) compared to 2344 (Hass) or 3147 (*drymifolia*) WGD paralog pairs within avocado.

We leverage this relatively high degree of orthology to circumvent the necessarily severe restrictions (e.g. MinL=4) on avocado x avocado syntenic block detection, by searching for two or more blocks in avocado whose orthologous blocks in the reference genome overlap by at least five genes, providing a “superblock” that potentially contains many more avocado duplicates than the paralogous syntenic blocks made without reference genomes. In other words, a superblock consists of two avocado regions, both of which are orthologous with some region (or two overlapping regions) on a chromosome in one of the reference genomes.

Figure S39. Relative frequencies of gene pair and syntenic superblock similarities. Block data derived from comparisons with 15 reference genomes. Pairs with sequence identity higher than 95% have been discarded (cf. Fig. S32), as being indicative of heterozygosity rather than paralogy. Blocks with average sequence identity less than 72% are not used in the subsequent analysis to avoid contamination from the gamma core eudicot whole genome triplication (31). AH: Hass, AD: *drymifolia*.

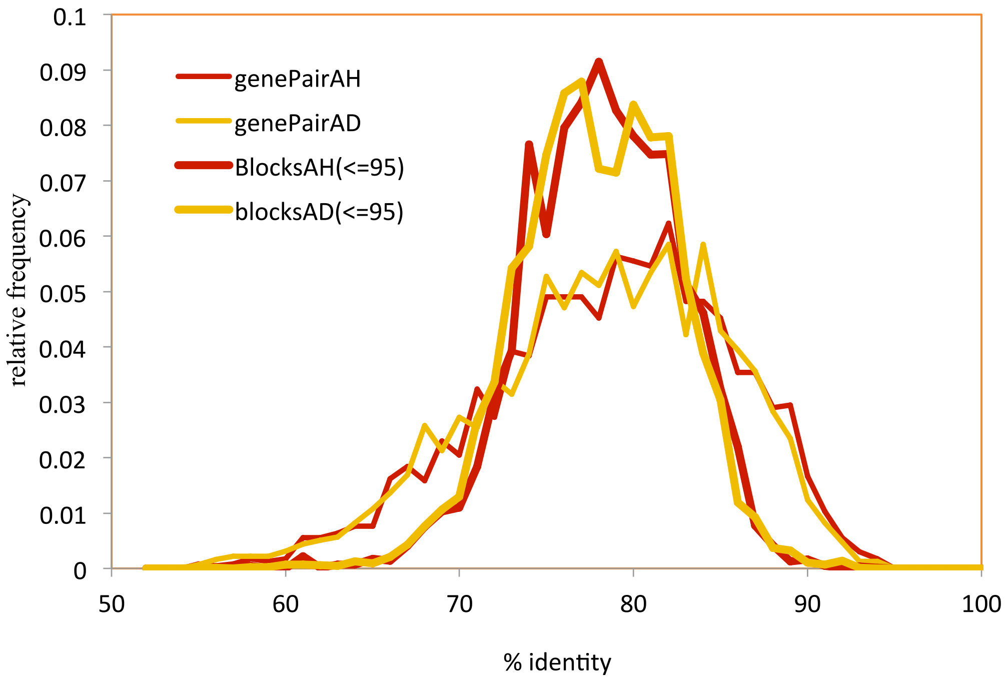

For each superblock we can calculate the average sequence identity of the duplicate gene pairs, generally much more numerous than the pairs in the paralogous blocks determined directly by SynMap These average similarities provide a more reliable indication of divergence time than the individual gene-pair scores, although a slight block-size bias is introduced, as discussed below.

It is important to note that the comparison of the two avocado regions making up a superblock has shed any explicit connection with the reference genomes. Moreover, the replicate construction for 15 different reference genomes accumulates many more superblocks than are obtainable from any one reference, due to the maximally independent fractionation and rearrangement histories of the organisms, including the earliest diverging angiosperm (*Amborella*), monocots (pineapple, duckweed, rice and sorghum), a basal eudicot (*Nelumbo*), various rosids (grape, watermelon, peach, poplar, *Arabidopsis*), a super-asterid (sugar beet) and asterids (*Mimulus*, tomato and coffee). Folding these 15 analyses into a single data sets yields three times as many pairs of blocks (9787 for Hass and 14636 for *drymifolia*) as the gene pairs discussed above (2344 and 3147, respectively). Just as important, the distribution of superblock similarities is considerably more compact than the distribution of gene pair similarities (standard deviations 4.4 and 4.5 for the blocks compared to 7.0 for both sets of gene pairs). This is evident in Figure S33, where there is also a slight shift in the mean from 78.5 and 78.2 for the genes to 78.2 and 78.0 with the blocks. (This bias is understandable in terms of recent blocks, having had less time to fractionate, tending to contain slightly more genes than earlier blocks, so there are proportionately fewer of the recent blocks).

Despite the sharpening up of the distributions of blocks compared to pairs in Figure S39, the bluntness of the peak, or broad range of maxima, persists, suggesting that the means of the two components of each distribution are at 76% and 81%. Although there is obviously much overlap, to distinguish between the earlier and more recent events, we establish a cut-off of 79%.

The proportion of superblocks with average gene pair sequence identity 79% or higher ("recent" pairs) is 0.5081 (Hass) and 0.4986 (*drymifolia*). We counted all triples of regions in the avocado genome, say A, B and C. where the three pairs of average sequence identity, AB, BC and CA were all between 62% and 95%. If the recent polyploidization event in avocado resulted in a whole genome triplication, we would expect the set of triples to be enriched for sets of three recent superblocks, i.e., AB, BC and CA all with average sequence identity above the cutoff, compared to three related superblocks chosen at random. If the recent polyploidization event resulted in a whole genome duplication, on the other hand, we would expect the set of triples to be enriched for only one of the superblocks AB, BC or CA to be recent, with the other two being of earlier vintage, compared to three superblocks chosen at random. The random distribution of high sequence identity superblocks in triples can be calculated via a binomial distribution with probability 0.5081 (Hass) and 0.4986 (*drymifolia*), e.g., the proportion of *drymifolia* triples with three similarities, AB, BC, CA all 79% or higher would be $(0.4986)^3=0.12395$. For 3731 triples this would produce around 463 such triples. The complete results are displayed in Table S15.

Table S15 Expected and observed numbers of triples of superblocks classified by recency of pairs of components.

| | pairs: triples | 3 recent | 2 recent | 1 recent | 0 recent | total |
|---|---|---|---|---|---|---|
| ***drymifolia*** | random | 462.5 | 1395.2 | 1403.0 | 470.3 | 3731 |
| | observed | 258 | 957 | 2190 | 326 | 3731 |
| **Hass** | random | 326.7 | 949.0 | 918.8 | 296.5 | 2491 |
| | observed | 201 | 691 | 1365 | 234 | 2491 |

Table S15 shows that there are far fewer triples with three recent superblocks than expected under the random model, and far more triples with only one recent superblock. Our main conclusion then is that the most recent polyploidization in avocado resulted in a whole genome duplication and not a whole genome triplication.

Triples with no recent superblocks could exceed expectations if the earlier event were a triplication. However, the results in Table S15 suggest an early whole genome duplication as well, based on this category of triples. Triples with two recent superblocks and one earlier one fit neither triplication nor duplication models. This results from the large overlap between the two components making up the distribution of average sequence identity of superblocks (cf. Figure S39) so that many superblocks originating in one polyploidization event are misclassified as originating in the other. The results in Table S15 on this category of triples suggest only that this misclassification is less frequent than a completely random assembly of superblock triples would produce.

The analysis described here is subject to some uncertainties. The proportions of recent superblocks (approximately 0.5) used to calculate the random results in Table S15 are subject to the block size bias mentioned above. However, this bias is much too small to account for the shortfall of observed recent triples compared to those predicted by the random model. In addition, the cutoff of 79% (or above) average sequence identity instead of 78.5% or 78% to assign superblocks to the more recent polyploidization event is somewhat subjective, but shifting the cut-off to 78%, say, which would increase the number of observed recent superblocks, would not materially affect the *contrast* between the observed and predicted number of triples in Table S15. Finally, avocado is a descendant of at least one earlier, pre-angiosperm plant polyploidization event, aside from the two in its own lineage, as well as other gene duplications, but judging from Figure S39 the cutoff to be included in our analysis, 72% for average superblock sequence identity, could only have allowed the inclusion of very few superblocks originating in these earlier events, in particular from the core eudicot gamma whole genome triplication.

### 7. Phylogenomic relationships of avocado to other angiosperms

#### 7.1. Phylogenetic relationships based on single-copy orthologs

We filtered our OrthoMCL orthogroups (section 3.1) to recover presumed gene families with single representative genes for every species compared and no missing representatives across all 19 species' proteomes. We generated gene alignments using MUSCLE (55) on amino acid sequences and filtered for alignment quality, requiring over 30% of the amino acid sequence alignment to be retained after using Gblocks (56) to remove poorly aligned positions with low stringency settings that permit smaller final blocks, gaps within final blocks, and less strict flanking positions; this resulted in 176 orthogroups remaining (Supplementary data 2). We thereafter concatenated each orthogroup alignment and generated phylogenetic trees using RAxML (57), based on either amino acid sequences or back-translated coding sequences. Models used for amino acid sequences and nuclear sequences were JTT+I+G and GTR, respectively, as determined using ProtTest3 (58) and jModelTest2 (59). We also ran coalescent-based species tree analyses; for this, we reconstructed gene trees for each of the individual gene alignments and fed the collection to ASTRAL (60) (using default parameters), which constructs a single species tree that reconciles relationships in the presence of gene tree conflict that can result from ILS or admixture. For coding sequences, each gene tree was run using the model prescribed by jModelTest2. For amino acid alignments, we used the best models automatically selected by RAxML for all genes, given the extreme computational requirements of model testing using ProtTest3. ASTRAL on either coding sequences or inferred amino acids produced the same topologies as the concatenated supermatrices analyzed using RAxML. Use of these alternative data transforms resulted in different resolutions of the three main clades of angiosperms represented among the 19 proteomes, as described in the main text. Based on protein sequences, avocado was resolved as sister to monocots plus eudicots (albeit with poor support on the single supermatrix tree of maximum likelihood; cf. (61, 62)), whereas from coding sequences, avocado was placed as sister to monocots only (cf. (63)) (Figures S40,41). All data sets and tree files are available as additional data files (Supplementary data 3).

Fig. S40. Phylogenetic trees based on amino acid alignments of 176 single-copy genes. Top, single tree of maximum likelihood from a concatenated set of alignments. Bootstrap supports are shown at nodes, indicating very poor support for early splits among the major angiosperm clades. Bottom, ASTRAL coalescence tree based on individual gene trees. Quadripartition supports are shown, which better support the resolution of avocado as sister to monocots+eudicots.

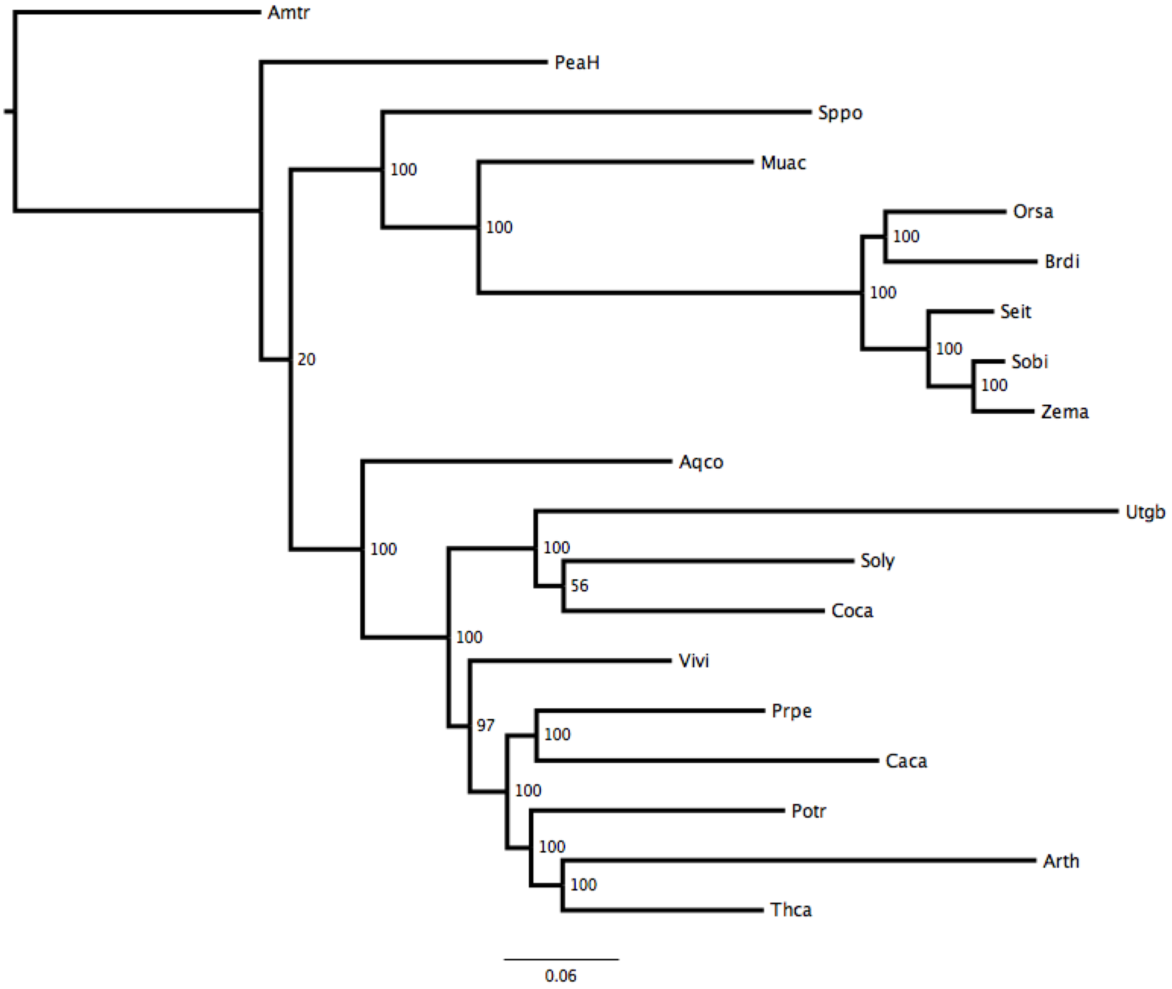

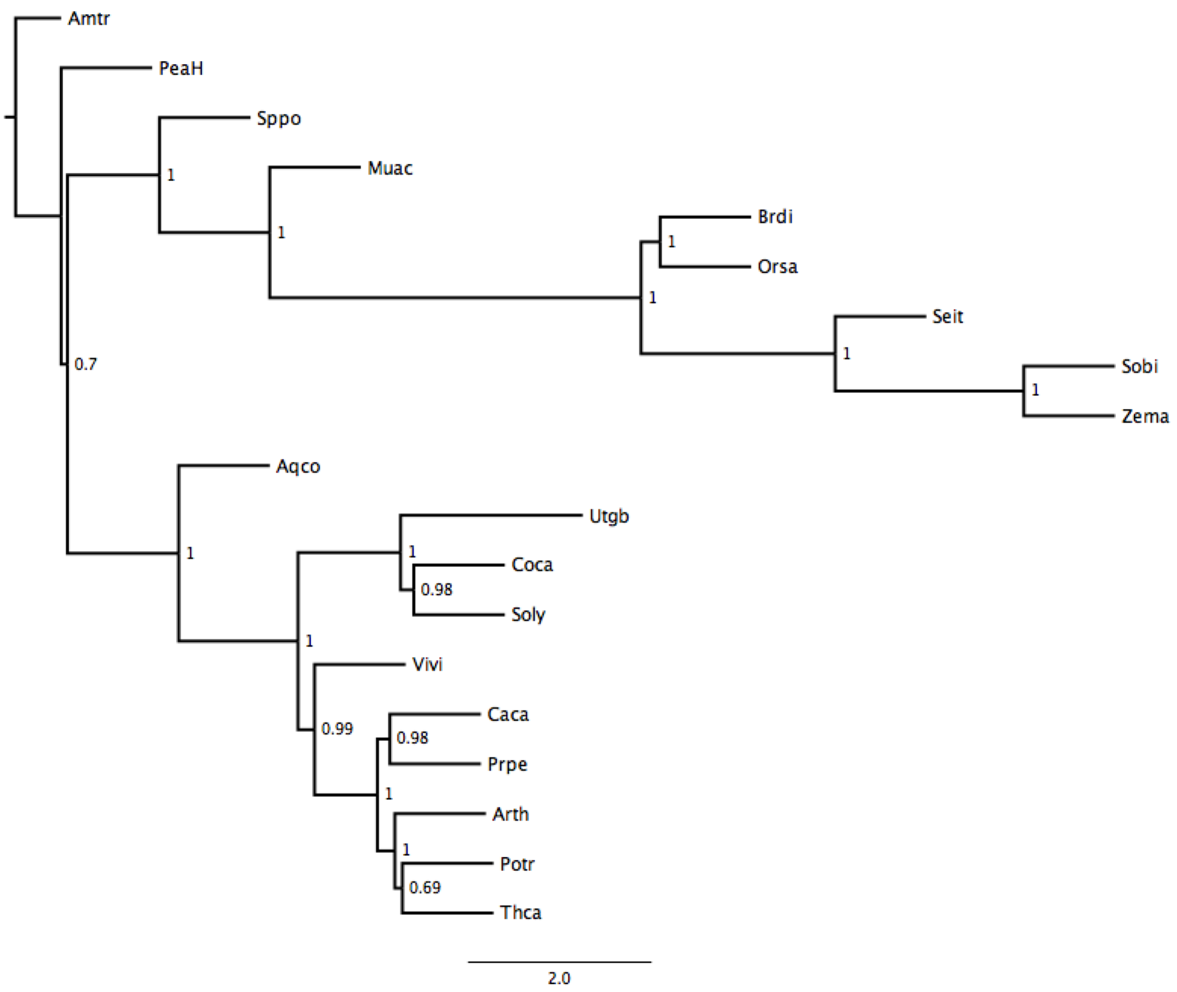
Amtr
PeaH
Sppo
1
Muac
1
Brdi
1
Orsa
1
Seit
1
Sobi
1
Zema
0.7
Aqco
1
Utgb
1
Coca
0.98
Soly
1
Vivi
0.99
Caca
0.98
Prpe
1
Arth
1
Potr
0.69
Thca
2.0

Fig. S41. Phylogenetic trees based on reverse-translated coding sequence alignments of 176 single-copy genes. Top, single tree of maximum likelihood from a concatenated set of alignments. Bootstrap supports are shown at nodes, indicating strong support for avocado as sister to monocots. Bottom, ASTRAL coalescence tree based on individual gene trees. Quadripartition supports are shown, which provide moderately good support for the resolution of avocado as sister to monocots.

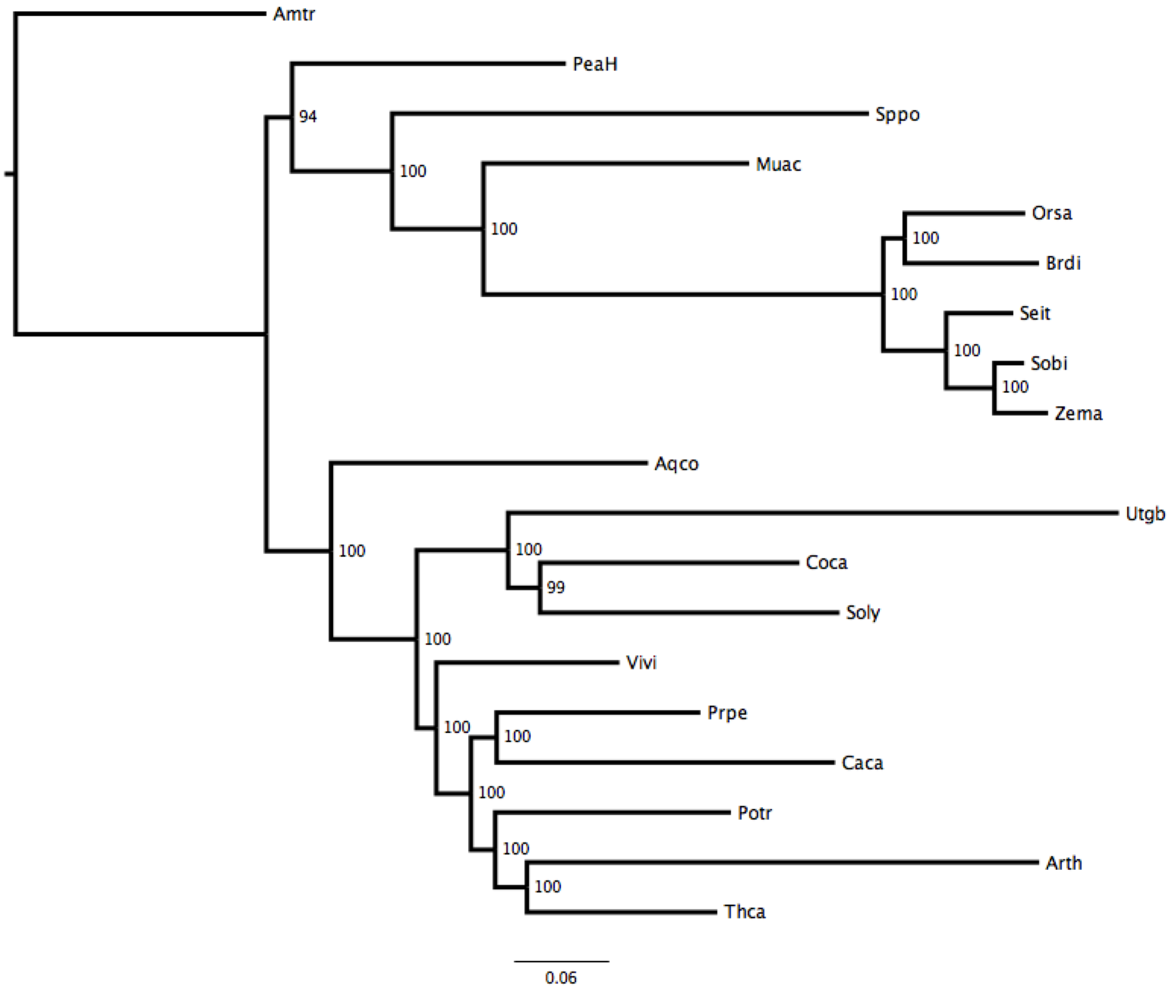

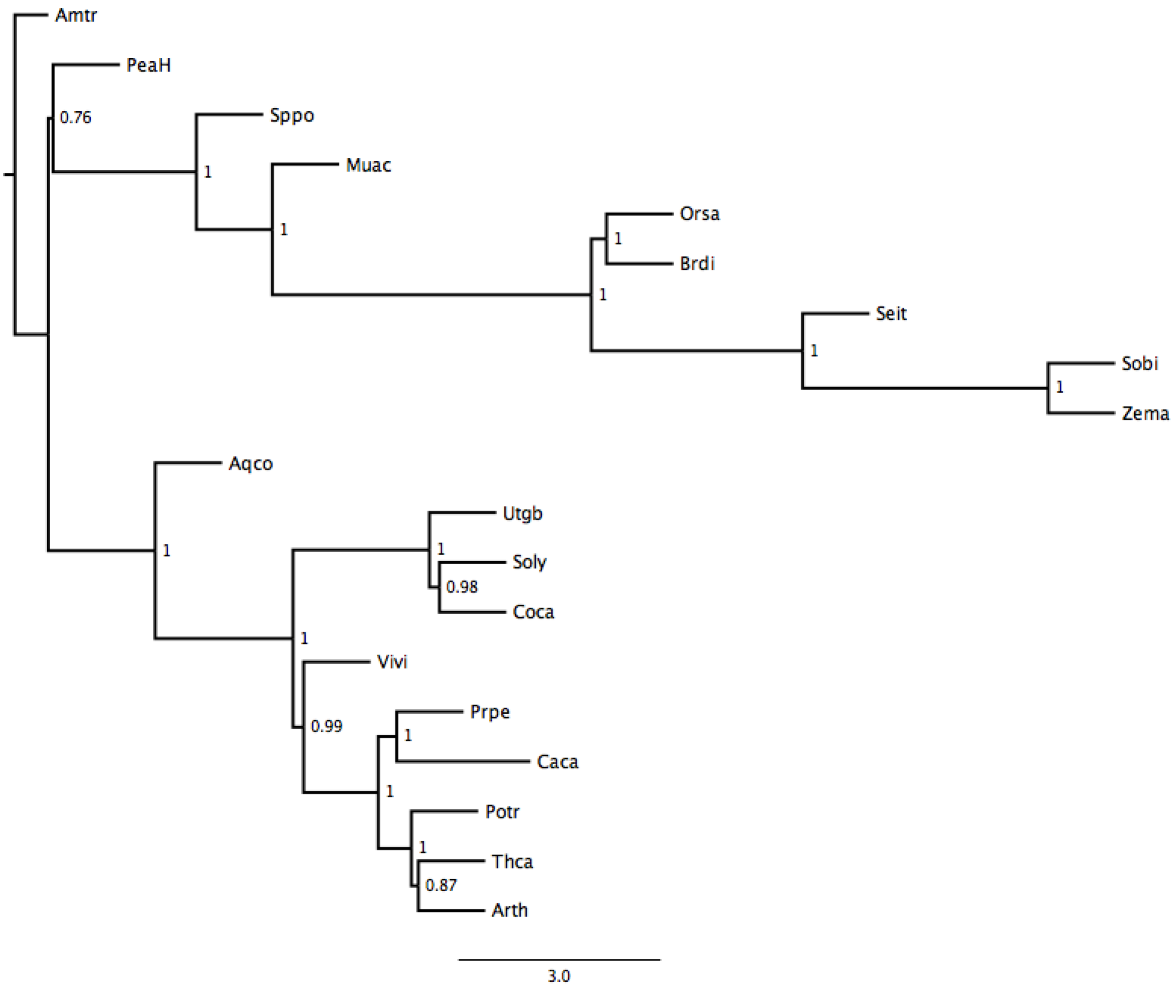

### 7.2. Phylogenomic analyses using the Orthofinder pipeline

In a different analysis we added to the 19 species *Gnetum* (a gymnosperm) and *Selaginella* (a non-seed plant) in orthogroup classification to generate a rooted species tree from all gene trees (4,694) that contained one or more (i.e., paralogous) gene copies from all species. OrthoFinder assigned 488282 genes (82.5% of total) to 17933 orthogroups. Fifty percent of all genes were in orthogroups with 34 or more genes (G50 was 34) and were contained in the largest 3966 orthogroups (O50 was 3966). As noted, there were 4694 orthogroups with all species present and 101 of these consisted entirely of single-copy genes. 644 well-supported, non-terminal duplications were observed. 629 support the best root and 15 contradict it. Note that aside from the unexpected phylogenetic resolution of *Arabidopsis* as sister to rosids, the topology (Figure S36) shows avocado placed sister to eudicots only, and concurs with expectation for *Gnetum* and *Selaginella* as successive sister taxa to angiosperms. Protein sequences for *Gnetum montanum* (64) were obtained from Data Dryad (https://doi.org/10.5061/dryad.0vm37.2), and primary transcripts for *Selaginella moellendorfii* were downloaded from Phytozome V12 (https://phytozome.jgi.doe.gov). Orthofinder v2.2.6 (65) was run with default settings to infer (66) and root (67) the species tree. Here, avocado was resolved as sister to eudicots only (Figure S42), a result similarly found in transcriptome-based analyses of large numbers of species (68, 69).

Fig. S42. Phylogenetic tree output from the Orthofinder pipeline, showing avocado sister to eudicots, as has similarly been shown in large transcriptomic datasets.

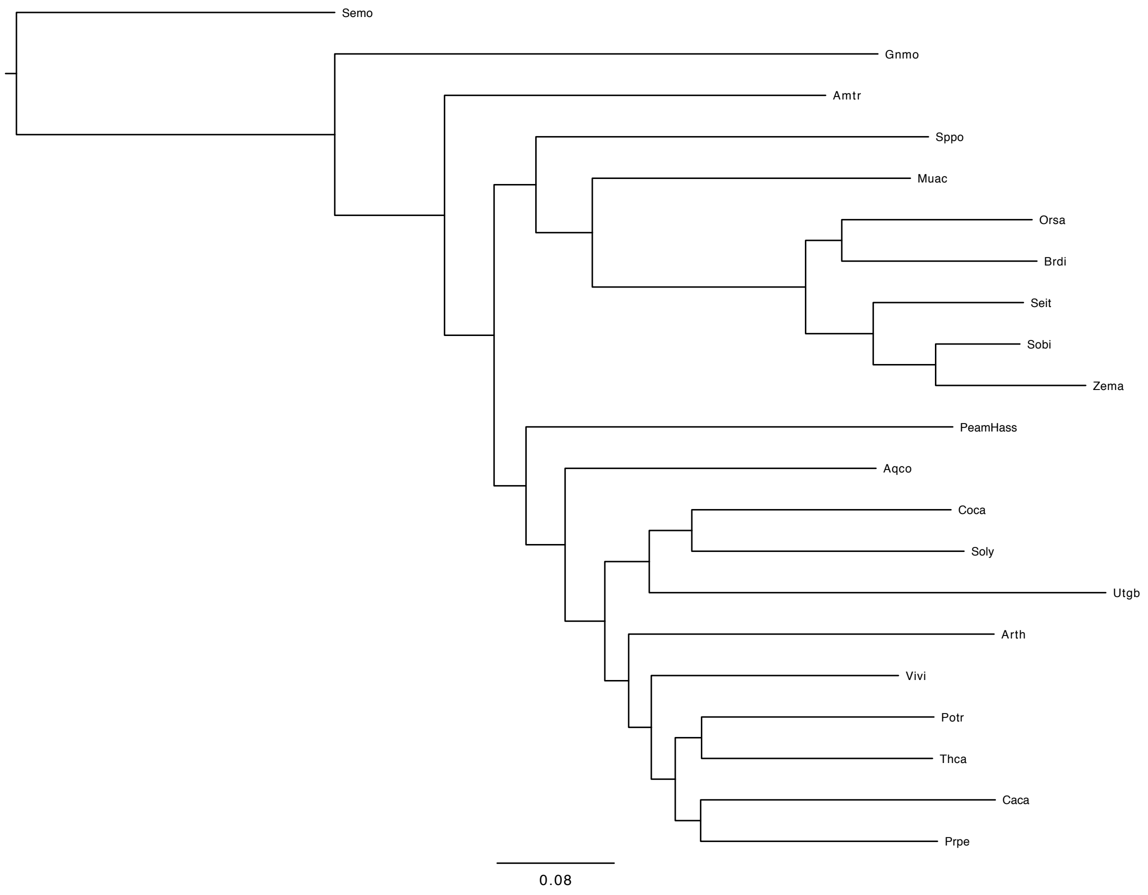

### 7.3. Phylogenomics using syntenic ortholog distances

In an altogether different approach (70, 71), we performed a phylogenomic analysis based on modal dissimilarity scores from syntenically-validated ortholog pairs (48).

Phylogenomic reconstruction based on dozens or hundreds of concatenated gene alignments may suffer from restrictions to widespread, single- or low-copy or best-versus-best genes and other biases in gene selection. As an alternative, we have proposed (72) to use neighbor-joining to construct an additive tree fit to a distance matrix whose elements are the modal dissimilarity scores of thousands of syntenically-validated ortholog pairs generated by the SynMap function on the CoGe platform (48, 49).

To illustrate, consider the distribution of syntenically valid ortholog dissimilarities between avocado and *Amborella* in Figure S43. (Syntenic validity refers to location of an ortholog pair adjacent to or in close chromosomal proximity to several other ortholog pairs in both genomes, with largely shared gene order.) This is calculated as the percentage of different bases in the two genomes over the entire CDS of the aligned genes.

Figure S43. Distribution of dissimilarity values of 7405 syntenically validated ortholog pairs in avocado and *Amborella.* Modal value is identified at dissimilarity = 24%.

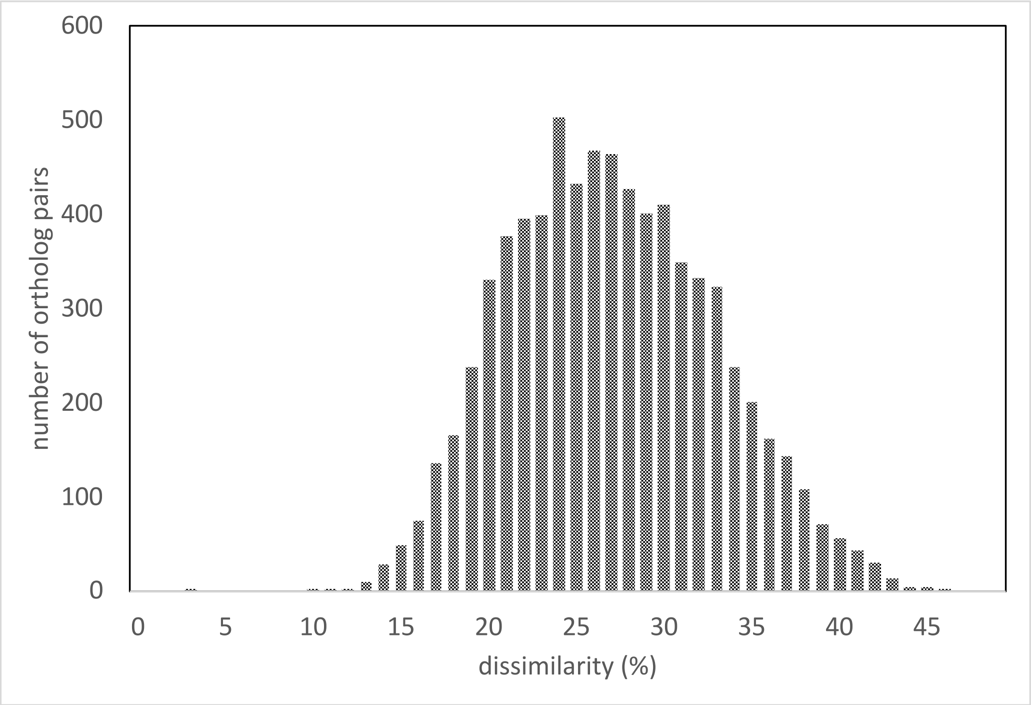

Including avocado and *Amborella*, we chose 14 species spanning the angiosperms, based on representing the major groupings, with a focus on genomes we had verified as useful in previous comparative studies. This included five monocots, five core eudicots and two basal eudicots. We compared each genome with every other genome, resulting in the construction of 14 x 13/2 = 91 distributions like the one in Figure S43. We identified the mode of each distribution as presented in Table S16.

Table S16. Location of mode in distributions of ortholog dissimilarities between pairs of 14 angiosperm genomes. Shaded blocks from left to right include the core eudicots, with the outlined subclades eurosids (including peach – a fabid, and cacao – a malvid), rosids, which also includes grape, and asterids, including coffee and tomato; the basal eudicots sacred lotus and columbine; and the monocots, including the early-branching alismatid duckweed, and the outlined commelinid clade containing banana from the order Zingiberales plus pineapple and the cereals from the order Poales. Also included are *Amborella*, constituting the earliest branching angiosperm lineage, and avocado as represented by the *P. drymifoli*a genome.

| | Peach | Cacao | Grape | Coffee | Tomato | Sacred lotus | Columbine | Duckweed | Banana | Pineapple | Rice | Sorghum | Avocado | Amborella |
|---|---|---|---|---|---|---|---|---|---|---|---|---|---|---|
| **Peach** | 0 | 22 | 22 | 24 | 26 | 25 | 24 | 29 | 27 | 29 | 32 | 28 | 26 | 27 |
| **Cacao** | 22 | 0 | 22 | 26 | 26 | 24 | 26 | 30 | 28 | 30 | 29 | 31 | 26 | 27 |
| **Grape** | 22 | 22 | 0 | 24 | 24 | 22 | 24 | 28 | 28 | 27 | 32 | 32 | 22 | 26 |
| **Coffee** | 24 | 26 | 24 | 0 | 25 | 27 | 27 | 29 | 27 | 30 | 33 | 32 | 26 | 28 |
| **Tomato** | 26 | 26 | 24 | 25 | 0 | 27 | 29 | 31 | 29 | 30 | 32 | 32 | 27 | 30 |
| **Sacred lotus** | 25 | 24 | 22 | 27 | 27 | 0 | 23 | 27 | 24 | 29 | 29 | 31 | 24 | 26 |
| **Columbine** | 24 | 26 | 24 | 27 | 29 | 23 | 0 | 34 | 29 | 26 | 31 | 30 | 26 | 26 |
| **Duckweed** | 29 | 30 | 28 | 29 | 31 | 27 | 34 | 0 | 27 | 26 | 31 | 28 | 29 | 28 |
| **Banana** | 27 | 28 | 28 | 27 | 29 | 24 | 29 | 27 | 0 | 22 | 28 | 27 | 27 | 28 |
| **Pineapple** | 29 | 30 | 27 | 30 | 30 | 29 | 26 | 26 | 22 | 0 | 24 | 25 | 27 | 30 |
| **Rice** | 32 | 29 | 32 | 33 | 32 | 29 | 31 | 31 | 28 | 24 | 0 | 15 | 30 | 32 |
| **Sorghum** | 28 | 31 | 32 | 32 | 32 | 31 | 30 | 28 | 27 | 25 | 15 | 0 | 29 | 33 |
| **Avocado** | 26 | 26 | 22 | 26 | 27 | 24 | 26 | 29 | 27 | 27 | 30 | 29 | 0 | 24 |
| **Amborella** | 27 | 27 | 26 | 28 | 30 | 26 | 26 | 28 | 28 | 30 | 32 | 33 | 24 | 0 |

In order to assess the phylogenetic origins of avocado with respect to *Amborella,* the monocots, the basal eudicots and the core eudicots, we first constructed a neighbor-joining tree, using the values in Table S16 as input, and rooted it to portray *Amborella* as the earliest branching lineage. The results confirmed all the groupings in Table S16, except for a three-way split between grape (known to evolve conservatively), the eurosids and the asterids within the core eudicots. The branching order was *Amborella*, avocado, the monocots, the basal eudicots and the core eudicots. One anomaly was the monophyletic grouping of the basal eudicots, instead of the expected sequential branching of columbine first, sacred lotus second, attributable to the lack of any other available sequenced genomes to include from the basal orders.

To test the robustness of this phylogenetic analysis, and not having available a bootstrap resampling of the primary data (frequency distributions over all ortholog pairs specific to each pair of genomes) we adopted a jackknife strategy, performing neighbor-joining analyses on each subset of 13 genomes, each subset omitting one of the full set of fourteen. *Amborella* was retained in each subset to provide a common rooting. For each tree, we decomposed it into 10 non-trivial bipartitions by removing one internal branch of the tree at a time.

In the 12 jackknife runs where avocado was present, it grouped with *Amborella,* and this is the key result for this study. The structure and position in the phylogeny of the monocot clade was retained in all runs. The ambiguous position of grape was resolved in favor of a grouping with the other rosids in a majority of the runs. In all of the runs containing both columbine and sacred lotus, they grouped together, and the phylogeny was simply the expected contraction of the full phylogeny with one genome removed. Only by omitting one of these two basal eudicots was the overall structure of the tree disrupted by anomalous placement of the other, either by sacred lotus grouping with the monocots or columbine with the rosids. This confirms the impression that the results of our method could only be improved with additional sequenced basal eudicot genomes.

The phylogeny constructed by assembling the bipartitions appearing in the majority of the jackknife runs is presented in Figure S44.

Figure S44. Phylogeny assembled from majority of bipartitions produced in one-genome-deleted-at-a-time jackknife taxon resampling and neighbor-joining. With *Amborella* as an outgroup to the rest of the angiosperms, avocado defines the earliest branch in all runs, followed by the monocots. Monophyletic basal eudicot group is likely an artifact of sparse taxon sampling of basal groups. Rosid grouping is blurred in a minority of samples.

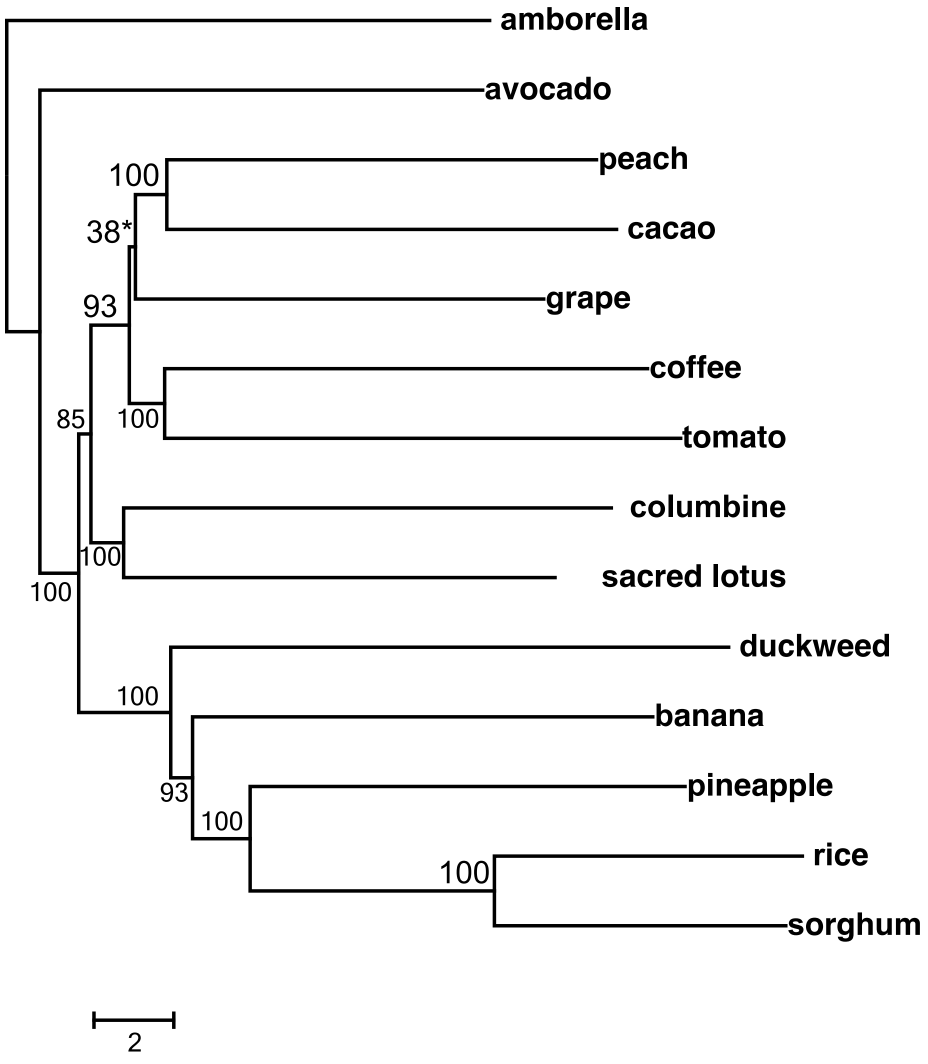

The method developed by Sankoff *et al.* (2016) (72) allows for the use of secondary modes in the distributions of gene pair similarities between genomes. These earlier modes represent whole genome duplication or triplication events in the shared early history of the two genomes. The addition of this information would lend a powerful boost to the discriminatory capacity of the methods. For example, columbine and sacred lotus both have whole genome duplication events in their history, but these are not shared between them. However, improvements in the detection and dating of common events are required before this methodology can be applied to data drawn from many genomes.

#### 7.4. Duplicate gene turnover analysis using BadiRate

Leveraging single-copy ortholog genes the phylogenetic position of avocado remained ambiguous, either as sister to monocots, sister to eudicots, or external to both. In order to provide complementary phylogenetic evidence, we apply BadiRate (73) to exploit multigene family data defined by OrthoMCL.

We then aligned the sequences of each orthogroup with the program M-Coffee (74) and used trimAl (75) to automatically remove poorly aligned regions. The best-fit amino acid substitution model for each multiple sequence alignment was selected using ProtTest (58) and specified in the RAxML analysis under a partitioned scheme. We finally used r8s to obtain the ultrametric trees required for the BadiRate (76) analysis, by applying the penalized likelihood algorithm (77) to the maximum-likelihood trees and fixing the age of the core eudicot node to 117 Mya (from TimeTree (78)). The three trees tested were:

(1) Avocado sister to monocots plus eudicots:

((PeaH:155.142880,((Sppo:131.526892,(Muac:111.383278,((Seit:23.464056,(Zema:12.795456,Sobi:12.795456):10.668600):15.854036,(Orsa:34.338245,Brdi:34.338245):4.979847):72.065186):20.143613):19.092998,(Aqco:134.027275,((((Prpe:90.160643,Caca:90.160643):8.850210,(Potr:92.938676,(Arth:86.697201,Thca:86.697201):6.241475):6.072177):9.012037,Vivi:108.022890):8.977110,((Coca:91.089278,Soly:91.089278):9.595857,Utgb:100.685135):16.314865):17.027275):16.592615):4.522990):49.711654,Amtr:204.854534);

(2) Avocado sister to monocots:

((((Sppo:129.665434,(Muac:109.873406,((Seit:23.183353,(Zema:12.646339,Sobi:12.646339):10.537013):15.645379,(Orsa:33.923527,Brdi:33.923527):4.905205):71.044675):19.792028):17.987786,PeaH:147.653220):3.809048,(Aqco:134.668594,((((Prpe:89.765241,Caca:89.765241):8.867591,(Potr:92.522586,(Arth:86.282914,Thca:86.282914):6.239672):6.110246):9.147039,Vivi:107.779870):9.220130,((Coca:90.876396,Soly:90.876396):9.647635,Utgb:100.524031):16.475969):17.668594):16.793674):51.149091,Amtr:202.611359);

(3) Avocado sister to eudicots:

(((Sppo:137.806675,(Muac:116.331068,((Seit:23.975138,(Zema:13.032166,Sobi:13.032166):10.942972):16.418656,(Orsa:35.219075,Brdi:35.219075):5.174719):75.937274):21.475607):18.511695,(PeaH:148.890366,(Aqco:133.665109,((((Prpe:90.522684,Caca:90.522684):8.845546,(Potr:93.306662,(Arth:87.054060,Thca:87.054060):6.252603):6.061568):8.914706,Vivi:108.282936):8.717064,((Coca:91.252387,Soly:91.252387):9.511621,Utgb:100.764008):16.235992):16.665109):15.225257):7.428004):50.870327,Amtr:207.188697);

We first contrasted four branch models to the three candidate topologies:

(i) **Global Rates model**: all branches have the same BDI (birth death, and innovation) rates.
(ii) **WGD/WGT model**: branches were grouped according to their corresponding number of WGD or WGT (whole genome triplication) events.
(iii) **WGD/WGT+Short model**: equivalent to WGD/WGT, with the addition that each branch shorter than 10 million years (my) was allowed to also have distinctive turnover rates.
(iv) **Free-Rates (FR) model**: BDI rates were allowed to vary in each phylogenetic branch.

The FR model ran for more than 350k hill-climbing iterations without reaching the strict convergence criteria implemented in BadiRate. Inspecting likelihood trajectories through iterations revealed however that likelihood values remained steady, with BadiRate spending a large number of iterations finely tuning parameters (Figure S45). We fitted a logistic growth model to these likelihood curves in order to predict the maximum likelihood values expected with further iterations. The predicted value was nevertheless lower than the ML score obtained by BadiRate (Table S18), which is only possible if the plateau reached by BadiRate's likelihood trajectory is stable, with an almost null gradient, and the predicted curve is fitted within the slightly erratic likelihood values obtained by BadiRate during the evaluation of new parameter proposals.

Table S17. (See additional file.) Statistics from BadiRate analyses of three different topological resolutions of avocado with respect to other major angiosperm lineages.

Figure S45. Maximum likelihood values for different topological placements of avocado from BadiRate under the FR model. The topology where avocado is sister to eudicots plus monocots is represented by the red curve, where it is sister to monocots in blue, and to eudicots in black.

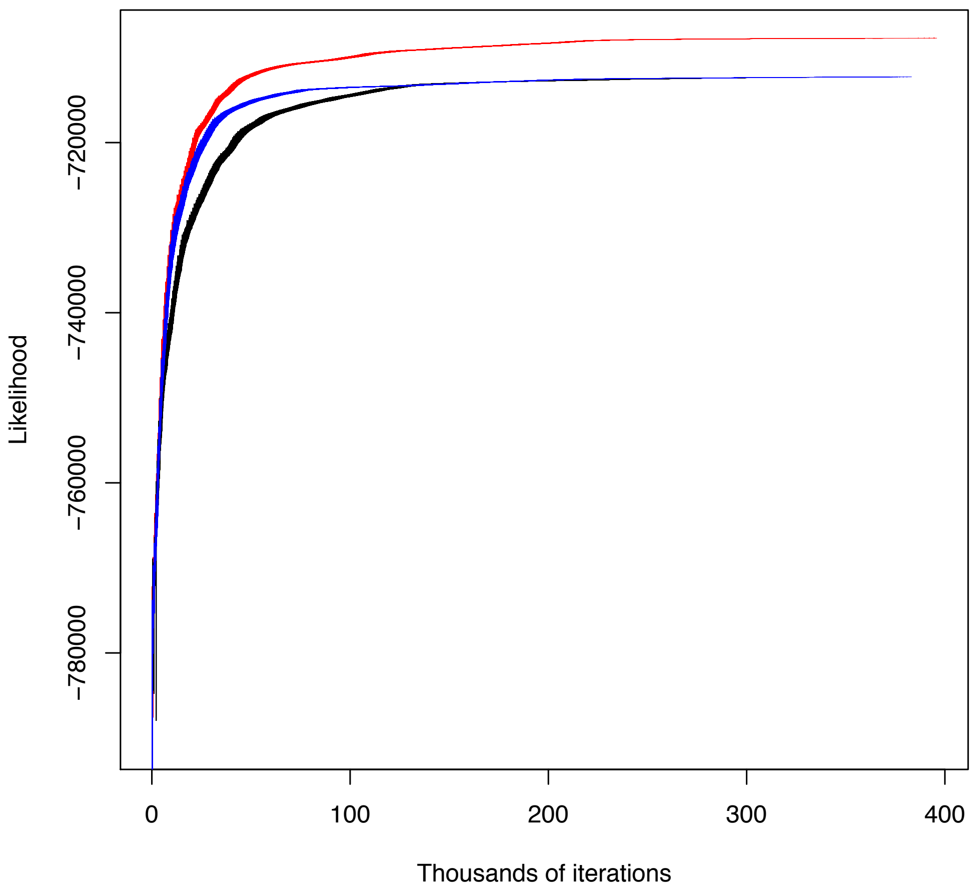

Despite slightly suboptimal FR likelihood values, the Akaike Information Criterion (AIC) clearly favored FR models, supporting heterogeneous rates of multigene family evolution across lineages (Table S17). Interestingly, such uneven rates of gene turnover cannot be entirely explained by lineage-specific WGD/WGT events, given that FR models fit multigene family data better than WGD/WGT models alone. Additionally, allowing for independent turnover rates in each short branch (<10 my) also improved likelihood and AIC values, although the fit was still worst than under the FR model.

Further exploration of FR estimates revealed inflated BDI turnover rates not only for short, but also for medium-length branches (20-50 my) (Table S18 and Figure S46). Since BDI rates are normalized by the corresponding branch lengths, this acceleration of the evolutionary rates at short time-scales cannot be due to methodological biases, but instead likely reflects incomplete lineage sorting. Indeed, gene copy number variation (CNV) within ancestral populations can lead to overestimates of gene family differences accumulated since the split of two recently-diverged species, unless ancestral polymorphisms are taken into account (79). This relation between branch lengths and gene turnover rates creates a BDI heterogeneity that can be only accommodated through the FR branch model.

Table S18. Correlation between branch length vs. turnover rates from BadiRate, on the tree where avocado is sister to monocots plus eudicots

| Correlation branch length vs. turnover rates | | | | | |
|---|---|---|---|---|---|
| Branch ID | Branch | Branch Length | Birth | Death | Innovation |
| 1 | 35->1 | 155.1429 | 0.0024022 | 0.0013356 | 0.0000116 |
| monocots+ eudicots, 2 | 35->34 | **4.523** | **0.0008563** | **0.0047665** | **0.0000002** |
| monocots, 3 | 34->14 | 19.093 | 0.0003971 | 0.0022256 | 0.0000022 |
| 4 | 14->2 | 131.5269 | 0.0011505 | 0.0015757 | 0.0000127 |
| 5 | 14->13 | 20.1436 | 0.0027527 | 0.0007067 | 0.0000028 |
| 6 | 13->3 | 111.3833 | 0.0042427 | 0.0009686 | 0.0000362 |
| 7 | 13->12 | 72.0652 | 0.0023985 | 0.0009779 | 0.0000159 |
| 8 | 12->8 | 15.854 | 0.0017812 | 0.0000002 | 0.0000161 |
| 9 | 8->4 | 23.4641 | 0.0163337 | 0.0072755 | 0.0001384 |
| 10 | 8->7 | 10.6686 | 0.0021833 | 0.0041048 | 0.0000157 |
| 11 | 7->5 | 12.7955 | 0.0215488 | 0.0080682 | 0.0002807 |
| 12 | 7->6 | 12.7955 | 0.0107149 | 0.0021493 | 0.000179 |
| 13 | 12->11 | 4.9798 | 0.0251492 | 0.0000045 | 0.0000354 |
| 14 | 11->9 | 34.3382 | 0.0025646 | 0.0030062 | 0.0001634 |
| 15 | 11->10 | 34.3382 | 0.0025709 | 0.0053834 | 0.0000576 |
| eudicots, 16 | 34->33 | 16.5926 | 0.0063922 | 0.0019342 | 0.0000049 |
| 17 | 33->15 | 134.0273 | 0.0038757 | 0.0018112 | 0.000006 |
| 18 | 33->32 | 17.0273 | 0.00075 | 0.0021454 | 0.0000051 |
| 19 | 32->26 | 8.9771 | 0.0000255 | 0.0021151 | 0.0000005 |
| 20 | 26->24 | 9.012 | 0.0018792 | 0.0000071 | 0.0000099 |
| 21 | 24->18 | 8.8502 | 0 | 0.0000583 | 0.0000021 |
| 22 | 18->16 | 90.1606 | 0.0035621 | 0.0014151 | 0.0000129 |
| 23 | 18->17 | 90.1606 | 0.0040874 | 0.0013231 | 0.0000157 |
| 24 | 24->23 | 6.0722 | 0.0011311 | 0.0025235 | 0.0000381 |
| 25 | 23->19 | 92.3787 | 0.0068664 | 0.0005692 | 0.0000305 |
| 26 | 23->22 | 6.2415 | 0.0048905 | 0.0067141 | 0.0000103 |
| 27 | 22->20 | 86.6972 | 0.0041158 | 0.0021948 | 0.0000209 |
| 28 | 22->21 | 86.6972 | 0.0026613 | 0.0010252 | 0.0000243 |
| 29 | 26->25 | 108.0229 | 0.0021567 | 0.001768 | 0.0000259 |
| 30 | 32->31 | 16.3149 | 0.00011 | 0.0014806 | 0.0000012 |
| 31 | 31->29 | 9.5959 | 0 | 0.0032281 | 0.0000012 |
| 32 | 29->27 | 91.0893 | 0.0028222 | 0.0014914 | 0.0000189 |
| 33 | 29->28 | 91.0893 | 0.0040253 | 0.0007225 | 0.0000382 |
| 34 | 31->30 | 100.6851 | 0.0021799 | 0.0027112 | 0.0000456 |

Figure S46. Relationship between birth, death, and innovation rates (y-axis) as a function of branch length (x-axis) on the tree where avocado is sister to monocots plus eudicots. Data from Table S18.

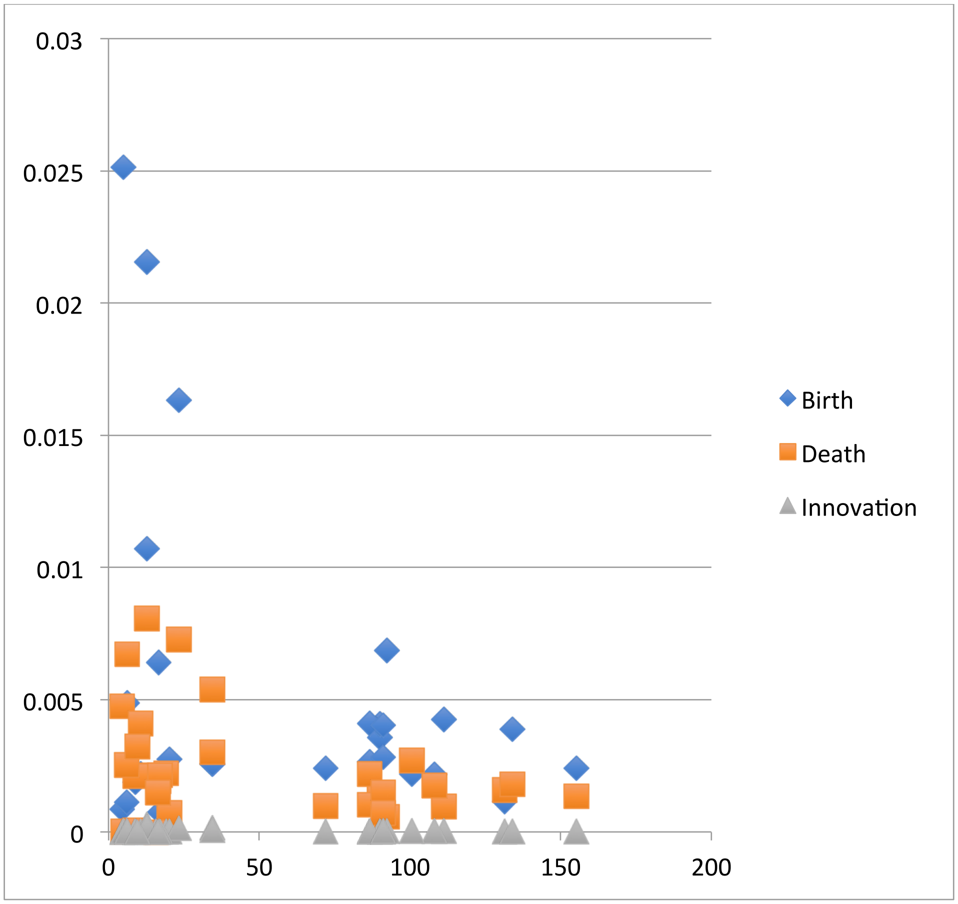

We next investigated whether multigene family data, evaluated under the FR model, preferentially supported one of the three candidate topologies. According to the AIC values, placing avocado as sister to both monocots and eudicots provided the greatest likelihood, in line with syntenic distance analyses (Section 7.3). Incongruence with phylogenetic reconstruction from 176 single-copy genes (Section 7.1) might reflect limited or biased information among 1:1 orthologs, due to the action of natural selection or ortholog misidentification. With large CNV variation in ancestral populations, as well as multiple WGD/WGT events followed by subsequent gene fractionation, the accurate identification of true orthologs based on all-against-all blast searches might be challenged.

The reconstruction of the ancestral gene content inferred under the best-fit model (FR and with avocado as sister to monocots plus eudicots) is illustrated in Figure S47.

Figure S47. Reconstruction of ancestral gene content inferred under the best-fit model (FR) and with avocado as sister to monocots plus eudicots. Also shown as main text Figure 3A.

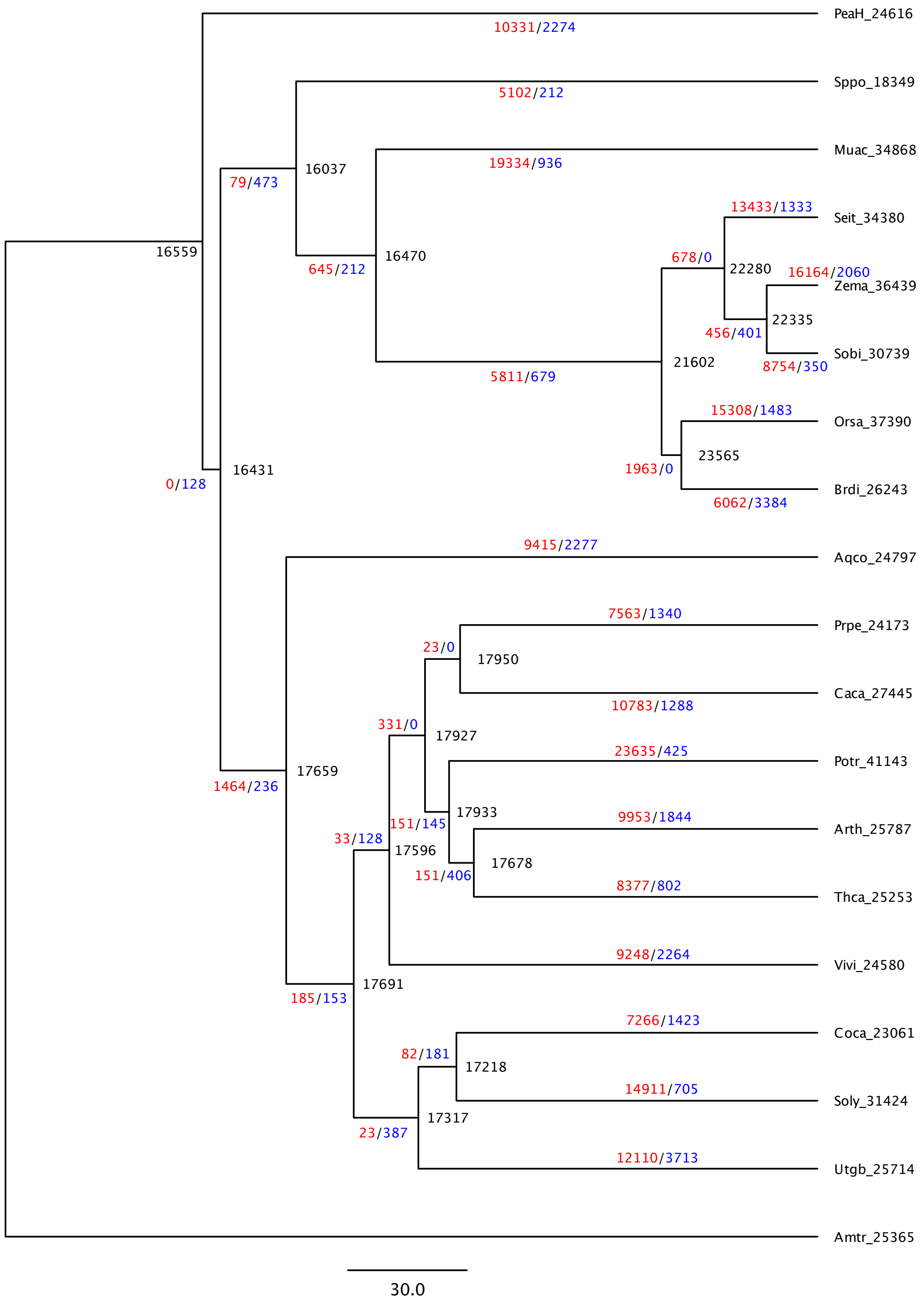

Three contentious phylogenetic topologies were investigated, where avocado is sister to eudicots and monocots, to monocots or to eudicots. The "GR", "WGD", "WGD short" and "FR" columns denote the likelihood values of the four branch models evaluated, and defined above. The "param" column indicates the number of parameters associated with each branch model. Given the likelihood and the number of parameters, the AIC columns report the Akaike Information Criterion. Lower AIC values indicate greater model support (best fit in red for each branch model). Since the FR model did not reach the strict convergence criteria implemented in BadiRate after ~400k hill-climbing iterations, a logistic growth function was fitted to these ~400k likelihood evaluations. The logistic regression predicted a maximum FR likelihood (column "FR predicted") that is on par with that obtained ~400k hill-climbing iterations (FR column), suggesting successful convergence. Therefore, the "FR_AIC" and "FR_AIC predicted" columns supported, consistently, that avocado is sister to both eudicots and monocots (Table S17).

### 8. Functional Enrichments in Duplicate Gene Space

Duplicate genes, in syntenic versus tandem bins (Table S19), were downloaded from CoGe self:self SynMap calculations. Gene models were annotated with the highest alignment score matches using tblastx versus the *Arabidopsis* coding sequences database v10.02 with an E-value cutoff of 1E-5. Generic gene ontology (GO) term annotations for *Arabidopsis* genes were downloaded from TAIR (http://arabidopsis.org/) and the avocado gene models were functionally annotated by assigning the GO terms from the best *Arabidopsis* gene hit. GO term enrichment analyses were carried out for subsets of foreground genes using all annotatable genes in the avocado genome as background and using Fisher's exact test implemented in GOATOOLS (https://github.com/tanghaibao/goatools) (Table S20). KEGG enrichment analysis was performed using the statistical framework from GOATOOLS using annotations from the KEGG pathway database downloaded from https://www.genome.jp/kegg/ (Table S21). The whole-genome background (Table S22) was custom-generated by selecting the set of avocado genes annotatable against *Arabidopsis* genes with E-value cutoff of 1E-05 and accepting the topmost hit as the match. Bonferroni correction for multiple tests was applied with $p < 0.05$ cutoff.

Table S19. (See additional file.) Syntenic and tandem duplicate gene bins in the Hass genome from CoGe SynMap default analysis, including their best *Arabidopsis* hit.

Table S20. (See additional file.) GO enrichment analysis of syntenic and tandem duplicates in the Hass genome.

Table S21. (See additional file.) KEGG enrichment analysis of syntenic and tandem duplicates in the Hass genome.

Table S22. (See additional file.) The whole-genome background used for enrichment analyses.

### 9. Differential Expression of Tandem Versus Polyploid Duplicates

Hass transcriptome reads for untreated control vs. pathogen-treated (80) were mapped to Hass gene models using Kallisto (81), normalized to transcript-per-million (TPM) values and thresholded by identifying genes with treatment/control log2 fold-change outside of the

[2,-2] interval. Fold changes were calculated from the averages of the 6/9 hr and 24 hr treatments (80). See Table S23. For each gene:

foldChange = mean(treatment)/mean(control)
if foldChange > 4:
gene is upregulated
if foldChange < 0.25:
gene is downregulated.

Functional enrichments were characterized as in section 8. Tandem duplicates showed significant enrichment among both up- and down-regulated genes ($p$ = 3.536e-09 and $p$ = 7.274e-07, Fisher's exact test, respectively), whereas polyploid duplicates did not show either pattern. Among tandem duplicates, we calculated functional enrichments within up- vs. down-regulated genes (Table S24). The only significantly enriched category was xyloglucan:xyloglucosyl transferase activity ($p$ = 0.038984; Fisher's exact test, Bonferroni corrected).

Table S23. (See additional file.) Data used for expression analysis and GO enrichment.

Table S24. Functional enrichments within up- vs. down-regulated genes.
